# Comparative Perturbation-Based Modeling of the SARS-CoV-2 Spike Protein Binding with Host Receptor and Neutralizing Antibodies : Structurally Adaptable Allosteric Communication Hotspots Define Spike Sites Targeted by Global Circulating Mutations

**DOI:** 10.1101/2021.02.21.432165

**Authors:** Gennady M. Verkhivker, Steve Agajanian, Deniz Yazar Oztas, Grace Gupta

## Abstract

In this study, we used an integrative computational approach focused on comparative perturbation-based modeling to examine molecular mechanisms and determine functional signatures underlying role of functional residues in the SARS-CoV-2 spike protein that are targeted by novel mutational variants and antibody-escaping mutations. Atomistic simulations and functional dynamics analysis are combined with alanine scanning and mutational sensitivity profiling for the SARS-CoV-2 spike protein complexes with the ACE2 host receptor are REGN-COV2 antibody cocktail (REG10987+REG10933). Using alanine scanning and mutational sensitivity analysis, we have shown that K417, E484 and N501 residues correspond to key interacting centers with a significant degree of structural and energetic plasticity that allow mutants in these positions to afford the improved binding affinity with ACE2. Through perturbation-based network modeling and community analysis of the SARS-CoV-2 spike protein complexes with ACE2 we demonstrate that E406, N439, K417 and N501 residues serve as effector centers of allosteric interactions and anchor major inter-molecular communities that mediate long-range communication in the complexes. The results provide support to a model according to which mutational variants and antibody-escaping mutations constrained by the requirements for host receptor binding and preservation of stability may preferentially select structurally plastic and energetically adaptable allosteric centers to differentially modulate collective motions and allosteric interactions in the complexes with the ACE2 enzyme and REGN-COV2 antibody combination. This study suggests that SARS-CoV-2 spike protein may function as a versatile and functionally adaptable allosteric machine that exploits plasticity of allosteric regulatory centers to fine-tune response to antibody binding without compromising activity of the spike protein.

## Introduction

The coronavirus disease 2019 (COVID-19) pandemic associated with the severe acute respiratory syndrome (SARS)^1–5^ has been at the focal point of biomedical research. SARS-CoV-2 infection is transmitted when the viral spike (S) glycoprotein binds to the host cell receptor leading to the entry of S protein into host cells and membrane fusion.^6–8^ The full-length SARS-CoV-2 S protein consists of two main domains, amino (N)-terminal S1 subunit and carboxyl (C)-terminal S2 subunit. The subunit S1 is involved in the interactions with the host receptor and includes an N-terminal domain (NTD), the receptor-binding domain (RBD), and two structurally conserved subdomains (SD1 and SD2). Structural and biochemical studies have shown that the mechanism of virus infection may involve spontaneous conformational transformations of the SARS-CoV-2 S protein between a spectrum of closed and receptor-accessible open forms, where RBD continuously switches between “down” and “up” positions where the latter can promote binding with the host receptor ACE2.^9–11^ The crystal structures of the S-RBD in the complexes with human ACE2 enzyme revealed structurally conserved binding mode shared by the SARS-CoV and SARS-CoV-2 proteins in which an extensive interaction network is formed by the receptor binding motif (RBM) of the RBD region.^12–16^ These studies established that binding of the SARS-CoV-RBD to the ACE2 receptor can be a critical initial step for virus entry into target cells. The rapidly growing body of cryo-EM structures of the SARS-CoV-2 S proteins detailed distinct conformational arrangements of S protein trimers in the prefusion form that are manifested by a dynamic equilibrium between the closed (“RBD-down”) and the receptor-accessible open (“RBD-up”) form required for the S protein fusion to the viral membrane.^17–34^ The cryo-EM characterization of the SARS-CoV-2 S trimer demonstrated that S protein may populate a spectrum of closed states by fluctuating between structurally rigid locked-closed form and more dynamic, closed states preceding a transition to the fully open S conformation.^26^ Conformational dynamics of SARS-CoV-2 trimeric spike glycoprotein in complex with receptor ACE2 suggesting considerable conformational heterogeneity of ACE2-RBD and continuous swing motions of ACE2-RBD in the context of SARS-CoV-2 S trimer. According to these experiments, the associated ACE2-RBD is relatively dynamic, showing three major conformational states with different angle of ACE2-RBD to the surface of the S trimer.^33^ Cryo-EM structural studies also mapped a mechanism of conformational events associated with ACE2 binding, showing that the compact closed form of the SARS-CoV-2 S protein becomes weakened after furin cleavage between the S1 and S2 domains, leading to the increased population of partially open states and followed by ACE2 recognition that can accelerate transformation to a fully open and ACE2-bound form priming the protein for fusion activation.^34^ These studies confirmed a general mechanism of population shifts between different functional states of the SARS-CoV-2 S trimers, suggesting that RBD epitopes can become stochastically exposed to the interactions with the host receptor ACE2.

The biochemical and functional studies using a protein engineering and deep mutagenesis have quantified binding mechanisms of SARS-CoV-2 interactions with the host receptor.^35, 36^ Deep mutational scanning of SARS-CoV-2 RBD revealed constraints on folding and ACE2 binding showing that many mutations of the RBD residues can be well tolerated with respect to both folding and binding. A surprisingly large number of amino acid modifications could even improve ACE2 binding, including important binding interface positions that enhance RBD expression (V367F and G502D) or enhance ACE2 affinity (N501F, N501T, and Q498Y)^35^. This comprehensive mutational scanning of the SARS-CoV-2 RBD residues suggested the evolutionary potential for compensation of deleterious mutations in the ACE2 interface reminiscent of multi-step escape pathways and highlighted the energetic plasticity of the SARS-CoV-2 interaction networks in which mutations may enhance binding affinity thus providing a roadmap for quantifying map immune-escape mutations. Using deep mutagenesis, it was also demonstrated that human ACE2 is only suboptimal for binding of the SARS-CoV-2 RBD as ACE2 variants near the interface can result in the improved binding and simultaneously enhance folding stability.^36^ Mutational landscape analysis showed a significant number of ACE2 mutations at the interface that enhance RBD binding, and the molecular basis for affinity enhancement can be rationalized from the structural analysis.^36^ Functional studies characterized the key amino acid residues of the RBD for binding with human ACE2 and neutralizing antibodies, revealing two groups of amino acid residues to modulate binding, where the SARS-CoV-2 RBD mutations N439/R426, L452/K439, T470/N457, E484/P470, Q498/Y484 and N501/T487 can result in the enhanced binding affinity for ACE2.^37^ Additionally, A475 and F486 in the SARS-CoV-2 RBD were identified as the key residues for the recognition of both their common functional receptor ACE2 and neutralizing antibodies, suggesting structural and energetic plasticity of the RBM residues involved in ACE2 recognition may induce mutational escape from the neutralizing antibodies targeting the RBD regions.

The rapidly growing structural studies of SARS-CoV-2 antibodies (Abs) have delineated molecular mechanisms underlying binding competition with the ACE2 host receptor, showing that combinations of Abs can provide a broad and efficient cross-neutralization effects through synergistic targeting of conserved and variable SARS-CoV-2 RBD epitope.^38–49^ Structural studies confirmed that the SARS-CoV-2 S protein can feature distinct antigenic sites and some specific Abs may allosterically inhibit the ACE2 receptor binding without directly interfering with ACE2 recognition.^44^ The SARS-CoV-2 Abs are divided into several main classes of which class 1 and class 2 antibodies target epitopes that overlap with the ACE2 binding site.^50, 51^ The structural studies revealed binding epitopes and binding mechanisms for a number of newly reported SARS-CoV-2 Abs targeting RBD regions and competing with ACE2 include B38 and H14 Abs,^52^ P2B-2F6,^53^ CA1 and CB6,^54^ CC12.1 and CC12.3,^55^ C105,^56^ and BD-23 Ab.^57^ The crystal structure of a neutralizing Ab CR3022 in the complex with the SARS-CoV-2 S-RBD revealed binding to a highly conserved epitope that is located away from the ACE2-binding site but could only be accessed when two RBDs adopt the “up” conformation.^42^ Subsequent structural and surface plasmon resonance studies confirmed that CR3022 binds the RBD of SARS-CoV-2 displaying strong neutralization by allosterically perturbing the interactions between the RBD regions and ACE2 receptor.^43^ The crystal structure of an RBD-EY6A Fab complex identified the highly conserved epitope located away from the ACE2 binding site, showing that EY6A can compete with CR3022 by targeting residues that are important for stabilizing the pre-fusion S conformation.^58^

The B.1.1.7 variant of the SARS-CoV-2, a descendant of the D614G lineage, has originated in UK and spread to 62 countries showing the increased transmissibility. 8 of the 17 mutations observed in this variant are accumulated in the S protein, featuring most prominently N501Y mutation that can increase binding affinity with ACE2 while eliciting immune escape and reduced neutralization of RBD-targeting Abs.^59–61^ A new SARS-CoV-2 lineage (501Y.V2) first detected in South Africa is characterized by 21 mutations with 8 lineage-defining mutations in the S protein, including three at important RBD residues (K417N, E484K and N501Y) that have functional significance and often induce significant immune escape.^62, 63^ Finally, the recently discovered new lineage, named P.1 (descendent of B.1.1.28), was observed in December in Brazil and contains a constellation of lineage defining mutations, including several mutations of known biological importance such as E484K, K417T, and N501Y mutations.^64, 65^

Functional mapping of mutations in the SARS-CoV-2 S-RBD that escape antibody binding using deep mutational scanning showed that the escape mutations cluster on several surfaces of the RBD and have large effects on antibody escape while a negligible negative impact on ACE2 binding and RBD folding.^66^ This illuminating study demonstrated that escape sites from antibodies can be constrained with respect to their effects on expression of properly folded RBD and ACE2 binding, suggesting that escape-resistant antibody cocktails can compete for binding to the same RBD region but have different escape mutations which limits the virus ability to acquire novel sites of immune escape in the RBD without compromising its binding to ACE2.^66^ Comprehensive mapping of mutations in the SARS-CoV-2 RBD that affect recognition by polyclonal human serum antibodies revealed that mutations in E484 site tend to have the largest effect on antibody binding to the RBD,^67^ and various functional neutralization assay experiments indicated that E484 modifications can reduce the neutralization potency by some antibodies by >10 fold.^67–69^ These studies also indicated that K417N and N501Y mutants can escape neutralization by some monoclonal antibodies but typically only modestly affected binding.^67, 70^ At the same time, mutations in the epitope centered around E484 position (G485, F486, F490) or in the 443–455 loop (K444, V445, L455, F456 sites) strongly affected neutralization for several Abs.^67–71^ Functional mapping of the SARS-CoV-2 RBD residues that affect binding of the REGN-COV2 cocktail showed that REGN10933 and REGN10987 are escaped by different mutations as mutation at F486 escaped neutralization only by REGN10933, whereas mutations at K444 escaped neutralization only by REGN10987, while E406W escaped both individual REGN-COV2 antibodies.^70^ This study confirmed that escape mutations at Q493, Q498 and N501 sites may enhance binding affinity with ACE2 and that escape mutations can also emerge in positions distant from the immediate proximity of the binding epitope, highlighting structural and energetic plasticity of the RBD regions and potential allosteric-based mechanism of immune escape.^70^ The REGN-COV2 cocktail (REG10987+REG10933) demonstrated significant potential in preventing mutational escape,^72^ several other antibody cocktails such as COV2-2130+COV2-2196,^73^ BD-368-2+BD-629^74^ and B38+H4^52^ displayed promising neutralization activities. Analysis of the molecular determinants and mechanisms of mutational escape showed that SARS-CoV-2 virus rapidly escapes from individual antibodies but doesn’t easily escape from the cocktail due to stronger evolutionary constraints on RBD-ACE2 interaction and RBD protein folding.^75^ According to this study, the key RBD positions critical for escape of antibody combinations include K444 which is important epitope residue for CoV2-06, P2B-2F6, S309 and REG10987 Abs as well as E484/F486 sites that are central for binding of CoV2-14 and REG10933. Functional analysis validated that mutations of these residues are responsible for viral escape from the individual Abs and in combination with other currently circulating variants (N501Y, K417N, E484K) may induce the reduced neutralization by the antibody cocktails.^75^ The SARS-CoV-2 501Y.V2 lineage that includes one cluster in NTD with four substitutions and a deletion (L18F, D80A, D215G, Δ242–244, and R246I), and another cluster of three substitutions in RBD (K417N, E484K, and N501Y) can confer neutralization escape from SARS-CoV-2 directed monoclonal antibodies and significantly increased neutralization resistance from individuals previously infected with SARS-CoV-2 virus.^76^ Moreover, three class 1 antibodies (CA1, CB6 and CC12.1) that target the ACE2-binding RBM region showed a complete lack of binding for 501Y.V2 variant, suggesting that mutations in the RBD and NTD clusters may to amplify the mutational escape from RBD-targeted Abs.

Structural and functional studies showed that the activity of mRNA vaccine-elicited antibodies to SARS-CoV-2 and circulating variant encoding E484K or N501Y or the K417N/E484K/N501Y combination can be reduced by a small but significant margin, suggesting that these mutations in individuals with compromised immunity may erode the effectiveness of vaccine elicited immunity.^77^ Importantly, it was also shown that neutralization by 14 of the 17 most potent tested mAbs can be partly reduced or even abolished by either K417N, or E484K, or N501Y mutations. Another latest study reported the preserved neutralization of N501Y, Δ69/70 + N501Y + D614G and E484K + N501Y + D614G viruses by BNT162b2 vaccine-elicited human sera.^77^ Consistent with other studies, it was shown that the neutralization against the virus with three mutations from the SA variant (E484K + N501Y + D614G) was slightly lower than the neutralization against the N501Y virus and the virus with three UK mutations (Δ69/70 + N501Y + D614G), but these differences were relatively small.^78^ SARS-CoV-2-S pseudo-viruses bearing either the reference strain or the B.1.1.7 lineage spike protein with sera of 40 participants who were vaccinated with the mRNA-based vaccine BNT162b2 showed largely preserved neutralization, indicating that the B.1.1.7 lineage will not escape BNT162b2-mediated protection.^79^ The recent data demonstrate reduced but still significant neutralization against the full B.1.351 variant following mRNA-1273 vaccination.^80^ New SARS-CoV-2 variants that resist neutralizing antibodies are now emerging in low frequencies in circulating SARS-CoV-2 populations. In particular, recent reports presented evidence of circulating SARS-CoV-2 spike N439K variants evading antibody-mediated immunity, particularly N439K mutation that confers resistance against several neutralizing monoclonal antibodies and reduces the activity of mRNA vaccine-elicited antibodies.^81^ Computational modeling and molecular dynamics (MD) simulations have been instrumental in predicting conformational and energetic determinants of SARS-CoV-2 mechanisms and the binding affinity and selectivity with the host receptor ACE2.^82–93^ Molecular mechanisms of the SARS-CoV-2 binding with ACE2 enzyme were analyzed in our recent study using coevolution and conformational dynamics.^94^ Using protein contact networks and perturbation response scanning based on elastic network models, we recently discovered existence of allosteric sites on the SARS-CoV-2 spike protein.^95^ By using molecular simulations and network modeling we recently presented the first evidence that the SARS-CoV-2 spike protein can function as an allosteric regulatory engine that fluctuates between dynamically distinct functional states.^96^

Coarse-grained normal mode analyses combined with Markov model and computation of transition probabilities characterized the dynamics of the S protein and the effects of mutational variants D614G and N501Y on protein dynamics and energetics.^97^ Using time-independent component analysis (tICA) and protein networks, another computational study identified the hotspot residues that may exhibit long-distance coupling with the RBD opening, showing that some mutations may allosterically affect the stability of the RBD regions.^98^ Molecular simulations reveal that N501Y mutation increases ACE2 binding affinity, and may impact the collective dynamics of the ACE2-RBD complex while mutations K417N and E484K reduce the ACE2-binding affinity by abolishing the interfacial salt bridges.^99^ The growing body of computational modeling studies investigating dynamics and molecular mechanisms of SARS-CoV-2 mutational variants produced inconsistent results that propose different mechanisms. The development of a more unified view and a working theoretical model which can explain the diverse experimental observations is an important area of current efforts in the field.

In this study, we used an integrative computational approach to examine molecular mechanisms underlying the functional role of K417, N439, E484 and N501 positions targeted by novel mutational variants in the SARS-CoV-2 S protein. We combined coarse-grained (CG) simulations and atomistic reconstruction of dynamics trajectories with dynamic fluctuation communication analysis, mutational sensitivity analysis and network community modeling to examine complexes of the SARS-CoV-2 S-RBD and dissociated S1 domain of the S protein formed with the ACE2 host receptor. Using distance fluctuations communication analysis and functional dynamics analysis, we determine and compare the distribution of regulatory centers in the RBD complexes with ACE2 and REGN-COV2 antibody cocktail (REG10987+REG10933). Using alanine scanning and mutational sensitivity analysis, we show that K417, E484 and N501 residues correspond to key interacting centers with a significant degree of structural and energetic plasticity that allow mutants in these positions to afford the improved binding affinity with ACE2. Through perturbation-based network modeling and community analysis of the SARS-CoV-2 RBD complexes with ACE2 we demonstrate that E406, N439, K417 and N501 residues serve as effector centers of allosteric interactions and anchor major inter-molecular communities that mediate long-range communication in the complexes. The results of the comparative network analysis with antibody complexes, we show that mutations in these positions can alter structural arrangements with antibodies and compromise their neutralization effects. These results suggest that antibody-escaping mutations target allosteric mediating hotspots with sufficient plasticity and adaptability to modulate and improve binding and allosteric signaling functions with the host receptor activity while reducing efficiency of antibody recognition and long-range communications. This analysis suggests that SARS-CoV-2 S protein may function as a versatile and functionally adaptable allosteric machine that exploits plasticity of allosteric regulatory centers to generate escape mutants that fine-tune response to antibody binding without compromising activity of the spike protein.

## Materials and Methods

### Coarse-Grained Molecular Simulations

Coarse-grained (CG) models are computationally effective approaches for simulations of large systems over long timescales. In this study, CG-CABS model^100–104^ was used for simulations of the crystal structures of the SARS-CoV-2 RBD complex with ACE2 (pdb id 6M0J)^15^ and complexes formed by the dissociated S1 domain of SARS-CoV-2 Spike bound to ACE2 (pdb id 7A91, 7A92)^34^ (Figure 1A-C). We also simulated the cryo-EM structure of the SARS-CoV-2 RBD complex with the Fab fragments of two neutralizing antibodies, REGN10933 and REGN10987 (pdb id 6XDG)^105^ (Figure 1D-F). In this high-resolution model, the amino acid residues are represented by Cα, Cβ, the center of mass of side chains and another pseudoatom placed in the center of the Cα-Cα pseudo-bond. In this model, the amino acid residues are represented by Cα, Cβ, the center of mass of side chains and the center of the Cα-Cα pseudo-bond. The CABS-flex approach implemented as a Python 2.7 object-oriented standalone package was used in this study to integrate a high-resolution coarse-grained model with robust and efficient conformational sampling proven to accurately recapitulate all-atom MD simulation trajectories of proteins on a long time scale.^104^ Conformational sampling in the CABS-flex approach is conducted with the aid of Monte Carlo replica-exchange dynamics and involves local moves of individual amino acids in the protein structure and global moves of small fragments.^100–102^

**Figure 1.**
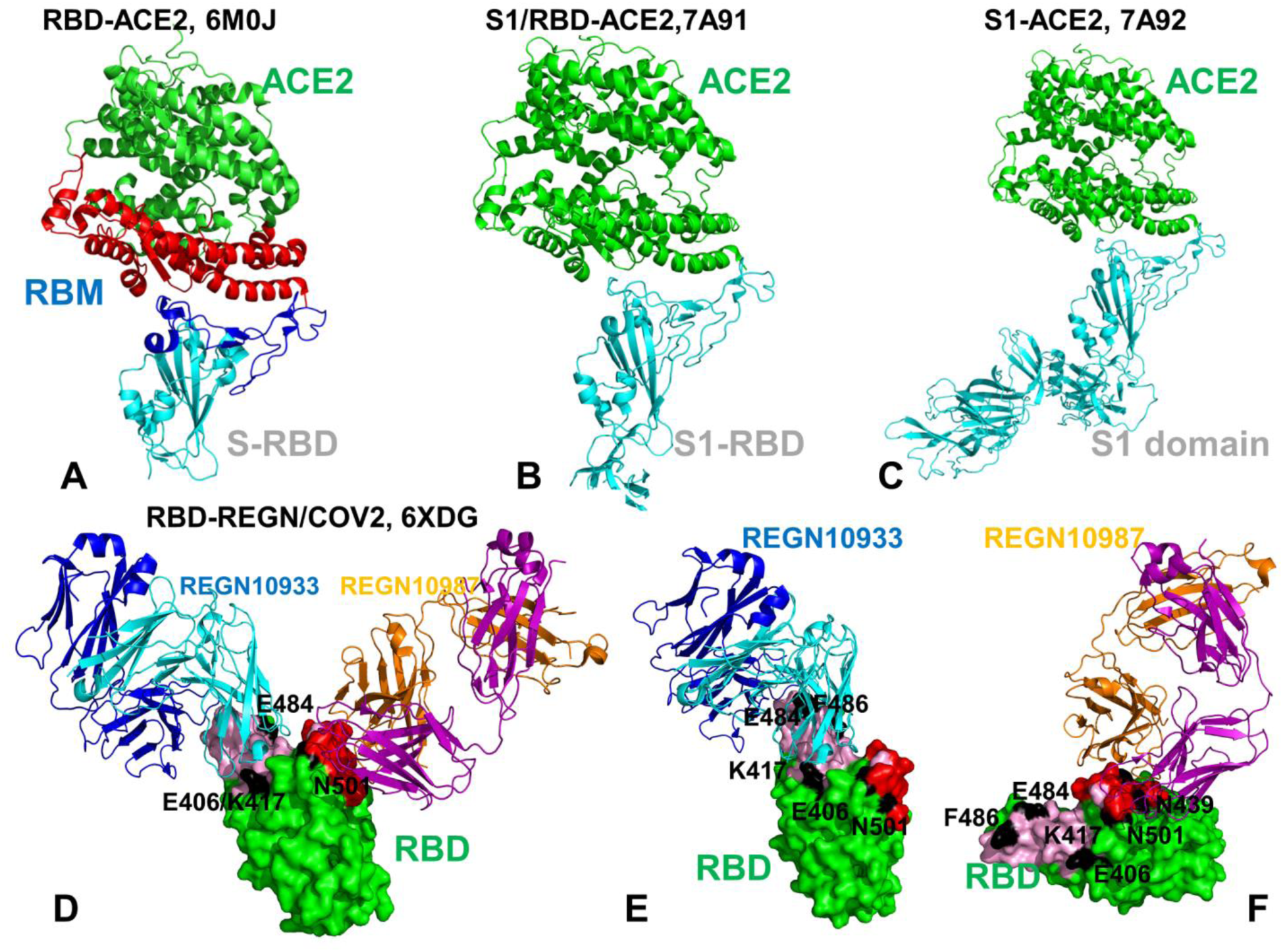
Crystal structures of the SARS-CoV-2 RBD and S1 domain complexes with ACE enzyme and REGN-COV2 antibody cocktail. (A) Structural overview of the SARS-CoV-2 RBD complex with ACE2 (pdb id 6M0J). The SARS-CoV RBD is shown in cyan ribbons and the RBM region is in blue ribbons. The subdomain I of human ACE2 is shown in red ribbons and the subdomain II is shown in green ribbons. The structure of ACE2 consists of the N-terminus subdomain I (residues 19-102, 290-397, and 417-430) and C-terminus subdomain II (residues 103-289, 398-416, and 431-615) that form the opposite sides of the active site cleft. (B) The crystal structure of the dissociated S1 domain form in the complex with ACE2 (pdb id 7A91). S1-RBD is in cyan ribbons and ACE2 is in green ribbons. (C) The crystal structure of the fully dissociated S1 domain in the complex with ACE2 (pdb id 7A92). S1 domain of the SARS-CoV-2 S protein is in cyan ribbons and ACE2 is in green ribbons. (D) The cryo-EM structure of the SARS-CoV-2 RBD in the complex with REGN10933/REGN10987 antibody cocktail. The RBD region is shown in green surface. REGN10933 Fab fragment is shown in ribbons with heavy chain in cyan and light chain in blue. REGN10987 is in ribbons with heavy chain in orange and light chain in purple. The positions of functional residues targeted by mutational variants and antibody-escaping mutations are E406, K417, E484 and N501 are annotated and highlighted as black patches on the RBD surface. (E) A close-up of the SARS-CoV-2 RBD interactions with REGN10933. The RBD is shown in green surface. REGN10933 Fab fragment is shown in ribbons with heavy chain in cyan and light chain in blue. The REGN10933 antibody epitope on RBD is highlighted in cyan patches on the surface. The positions of E406, K417, E484, F486, N501 are shown as black surface patches on the RBD. (F) A close-up of the SARS-CoV-2 RBD interface with REGN10987. The red patches correspond to the REGN10987 epitope. The positions of E406, K417, N439, E484, F486, N501 are shown as black surface patches on the RBD.

The default settings were applied in which soft native-like restraints are imposed only on pairs of residues fulfilling the following conditions : the distance between their *C*^α^ atoms was smaller than 8 Å, and both residues belong to the same secondary structure elements. A total of 1,000 independent CG-CABS simulations were performed for each of the studied systems. In each simulation, the total number of cycles was set to 10,000 and the number of cycles between trajectory frames was 100. MODELLER-based reconstruction of simulation trajectories to all-atom representation provided by the CABS-flex package was employed to produce atomistic models of the equilibrium ensembles for studied systems.

### Structure Preparation and Analysis

All structures were obtained from the Protein Data Bank.^106, 107^ During structure preparation stage, protein residues in the crystal structures were inspected for missing residues and protons. Hydrogen atoms and missing residues were initially added and assigned according to the WHATIF program web interface.^108, 109^ The structures were further pre-processed through the Protein Preparation Wizard (Schrödinger, LLC, New York, NY) and included the check of bond order, assignment and adjustment of ionization states, formation of disulphide bonds, removal of crystallographic water molecules and co-factors, capping of the termini, assignment of partial charges, and addition of possible missing atoms and side chains that were not assigned in the initial processing with the WHATIF program. The missing loops in the studied crystal structures of the dissociated S1 domain complexes with ACE2 (residues 556-573, 618-632) were reconstructed and optimized using template-based loop prediction approaches ModLoop,^110^ ArchPRED server^111^ and further confirmed by FALC (Fragment Assembly and Loop Closure) program.^112^ The side chain rotamers were refined and optimized by SCWRL4 tool.^113^ The shielding of the receptor binding sites by glycans is an important common feature of viral glycoproteins, and glycosylation on SARS-CoV proteins can camouflage immunogenic protein epitopes.^114, 115^ The atomistic structures from simulation trajectories of the dissociated S1 domain complex with ACE2 (pdb id 7A92) were elaborated by adding N-acetyl glycosamine (NAG) glycan residues and optimized. The glycosylated microenvironment for atomistic models of the simulation trajectories was mimicked by using the structurally resolved glycan conformations for most occupied N-glycans as determined in the cryo-EM structures of the SARS-CoV-2 spike S trimer in the closed state (K986P/V987P,) (pdb id 6VXX) and open state (pdb id 6VYB).

### Functional Dynamics and Collective Motions Analysis

We performed principal component analysis (PCA) of reconstructed trajectories derived from CABS-CG simulations using the CARMA package^116^ and also determined the essential slow mode profiles using elastic network models (ENM) analysis.^117^ Two elastic network models: Gaussian network model (GNM)^117, 118^ and Anisotropic network model (ANM) approaches^119^ were used to compute the amplitudes of isotropic thermal motions and directionality of anisotropic motions. The functional dynamics analysis was conducted using the GNM in which protein structure is reduced to a network of *N* residue nodes identified by *C_α_* atoms and the fluctuations of each node are assumed to be isotropic and Gaussian. Conformational mobility profiles in the essential space of low frequency modes were obtained using ANM server^119^ and DynOmics server.^120^

### Local Structural Parameters : Relative Solvent Accessibility

We have computed the relative solvent accessibility parameter (RSA) that is defined as the ratio of the absolute solvent accessible surface area (SASA) of that residue observed in a given structure and the maximum attainable value of the solvent-exposed surface area for this residue.^121^ According to this model, residues are considered to be solvent exposed if the ratio value exceeds 50% and to be buried if the ratio is less than 20%. Analytical SASA is estimated computationally using analytical equations and their first and second derivatives and was computed using web server GetArea.^121^

### Mutational Sensitivity Analysis and Alanine Scanning

To compute protein stability and binding free energy changes in the SARS-CoV-2 RBD structures upon complex formation with ACE2 receptor and REGN-COV2 antibody cocktail, we conducted a systematic alanine scanning of protein residues in the SARS-CoV-2 RBD and S1 domain. In addition, a complete mutational sensitivity analysis was done for binding free energy hotspots and residues E406, K417, N439, K444, E484, F486, and N501 targeted by widely circulating and antibody-escaping mutations. Alanine scanning and mutational sensitivity profiling of protein residues was performed using BeAtMuSiC approach.^122, 123^ If a free energy change between a mutant and the wild type (WT) proteins ΔΔG= ΔG (MT)-ΔG (WT) > 0, the mutation is destabilizing, while when ΔΔG <0 the respective mutation is stabilizing. BeAtMuSiC approach is based on statistical potentials describing the pairwise inter-residue distances, backbone torsion angles and solvent accessibilities, and considers the effect of the mutation on the strength of the interactions at the interface and on the overall stability of the complex.^122, 123^ The reported protein stability and binding free energy changes are based on the ensemble averages of BeAtMuSiC values using equilibrium samples from reconstructed simulation trajectories.

### Perturbation Response Scanning

Perturbation Response Scanning (PRS) approach^124, 125^ was used to estimate residue response to external forces applied systematically to each residue in the protein system. This approach has successfully identified hotspot residues driving allosteric mechanisms in single protein domains and large multi-domain assemblies.^126–131^ The implementation of this approach follows the protocol originally proposed by Bahar and colleagues^126, 127^ and was described in details in our previous studies.^96^ In brief, through monitoring the response to forces on the protein residues, the PRS approach can quantify allosteric couplings and determine the protein response in functional movements. In this approach, it 3N × 3*N* Hessian matrix ***H*** whose elements represent second derivatives of the potential at the local minimum connect the perturbation forces to the residue displacements. The 3*N*-dimensional vector Δ***R*** of node displacements in response to 3*N*-dimensional perturbation force follows Hooke’s law ***F*** = ***H*** ∗ ***ΔR***. A perturbation force is applied to one residue at a time, and the response of the protein system is measured by the displacement vector Δ***R***(*i*) = ***H***^−**1**^***F***(***i***) that is then translated into *N*×*N* PRS matrix. The second derivatives matrix ***H*** is obtained from simulation trajectories for each protein structure, with residues represented by *C_α_* atoms and the deviation of each residue from an average structure was calculated by Δ**R**_*j*_(*t*) = **R**_*j*_(*t*) − 〈**R**_*j*_(*t*)〉, and corresponding covariance matrix C was then calculated by Δ**R**Δ**R**^*T*^. We sequentially perturbed each residue in the SARS-CoV-2 spike structures by applying a total of 250 random forces to each residue to mimic a sphere of randomly selected directions. The displacement changes, Δ***R***^***i***^ is a *3N-*dimensional vector describing the linear response of the protein and deformation of all the residues.

Using the residue displacements upon multiple external force perturbations, we compute the magnitude of the response of residue 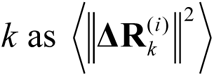 averaged over multiple perturbation forces **F**^(*i*)^, yielding the *ik*^th^ element of the *N*×*N* PRS matrix.^126, 127^ The average effect of the perturbed effector site *i* on all other residues is computed by averaging over all sensors (receivers) residues *j* and can be expressed as〈(Δ***R***^***i***^)^2^〉_*effector*_. The effector profile determines the global influence of a given residue node on the perturbations in other protein residues and can be used as proxy for detecting allosteric regulatory hotspots in the interaction networks. In turn, the *j* ^th^ column of the PRS matrix describes the sensitivity profile of sensor residue *j* in response to perturbations of all residues and its average is denoted as 〈(Δ***R***^***i***^)^2^〉_*sensor*_. The sensor profile measures the ability of residue *j* to serve as a receiver (or transmitter) of dynamic changes in the system.

### Network Modeling and Community Analysis

A graph-based representation of protein structures^132, 133^ is used to represent residues as network nodes and the inter-residue edges to describe residue interactions. The details of graph construction using residue interaction cut-off strength (*I*_min_) were outlined in our previous studies.^96^ We constructed the residue interaction networks using both dynamic correlations^134^ and coevolutionary residue couplings^135^ that yield robust network signatures of long-range couplings and communications. The details of this model were described in our previous studies.^135–137^ The ensemble of shortest paths is determined from matrix of communication distances by the Floyd-Warshall algorithm.^138^ Network graph calculations were performed using the python package NetworkX.^139^

The Girvan-Newman algorithm^140–142^ is used to identify local communities. An improvement of Girvan-Newman method was implemented where all highest betweenness edges are removed at each step of the protocol. The algorithmic details of this modified scheme were presented in our recent study.^143, 144^ The network parameters were computed using the python package NetworkX^139^ and Cytoscape package for network analysis.^145, 146^

## Results and Discussion

### Conformational Dynamics Profiles of the SARS-CoV-2 S RBD and Dissociated S1 Domain Binding with ACE2 : Balancing Structural Rigidity and Plasticity at Binding Interfaces

Multiple CG-CABS simulations of the SARS-CoV-2 RBD and S1 domain complexes with the ACE2 host receptor were performed to analyze similarities and differences in th conformational dynamics profiles of the RBD regions and specifically binding interface residues (Figure 2). We combined CG-CABS simulations with atomistic reconstruction of simulation trajectories to characterize regions of structural stability and plasticity at the ACE2 binding interfaces and determine the effect of the complete S1 domain in the complex on flexibility of the interacting residues and functional RBD regions. To our knowledge this is the first comparative computational analysis of S1 domain binding with ACE2 initiated to understand structural and dynamic rearrangements of the S1 domain to form a stable monomeric complex with ACE2. The conserved core of SARS-CoV-RBD consists of five antiparallel β strands with three connecting α-helices (Figures 1,2). The central β strands (residues 354-363, 389-405, 423-436) in SARS-CoV-2 RBD are stable and, as expected, only small thermal fluctuations were observed in these regions (Figure 2A). The anti-parallel β-sheets (β5 and β6) (residues 451-454 and 492-495 in SARS-CoV-RBD) that anchor the RBM region to the central core also displayed a significant stabilization in the complex with ACE2. The small α-helical segments of the RBD (residues 349-353, 405-410, and 416-423) also displayed a significant stability in simulations. These regions become even more rigidified in the complex formed by the dissociated S1 domain (Figure 2B). Of special interest were the ACE2 induced changes in the RBM region (residues 437–508) and particularly in a stretch of residues 471-503 involved in multiple contacts with the ACE2 receptor. The overall similar RBM profiles in all systems were seen but the greater stabilization of the interfacial residues in the ACE2 complex was observed for the bound S1 domain structure (Figure 2A,B). The interfacial loop residues 436-455 containing an important motif 444-KVGGNYNY-451 displayed significantly reduced fluctuations. Among residues that experience a more pronounced stabilization in the complex are K417, G446, Y449, Y453, L455, F456, Y473, A475, and G476 positions in the middle segment of the RBM (Figure 2C-E). The analysis of the intermolecular contacts in the SARS-CoV-2 RBD and S1 complexes with ACE2 (Supporting Information, Tables S1-S3) indicated a significant number of RBM interactions formed by the middle segment of the interface (K417, Y453, L455, F456, and Q493) with K31 and E35 of ACE2 which may explain a pronounced stabilization of these positions in simulations. Indeed, K417 forms contacts with D30 and H34 hotspot ACE2 residues, L455 is involved in interactions with K31 and D30, and F456 forms stabilizing contacts with D30, K31, and T27 ACE2 hotspots (Supporting Information, Tables S1-S3). Another group of residues in the RBM ridge involved in multiple binding contacts included E484 and F486 sites that interact with K31, Q24, M82, and L79 of ACE2 (Supporting Information, Tables S1-S3). Notably, this analysis showed that the interacting RBD motif 495-YGFQPTNG-502 is involved in the most persistent interaction contacts with ACE2 that is exemplified by stabilization of Y489, F490, Q493, Y495, G496, Q498, T500, and Y505 residues (Figure 2). These RBD residues form the largest number of contacts with ACE2 (Supporting Information, Tables S1-S3) and experienced the most significant stabilization in the complex (Figure 2A,B). Importantly, not only these residues showed markedly reduced fluctuations, but the large interfacial stretch of residues across entire binding interface (residues 486-FNCYFPLQSYGFQ-498) including key Q493 and Q498 interacting sites exhibited even a stronger stabilization in the complex formed by the dissociated S1 domain (Figure 2B).

**Figure 2.**
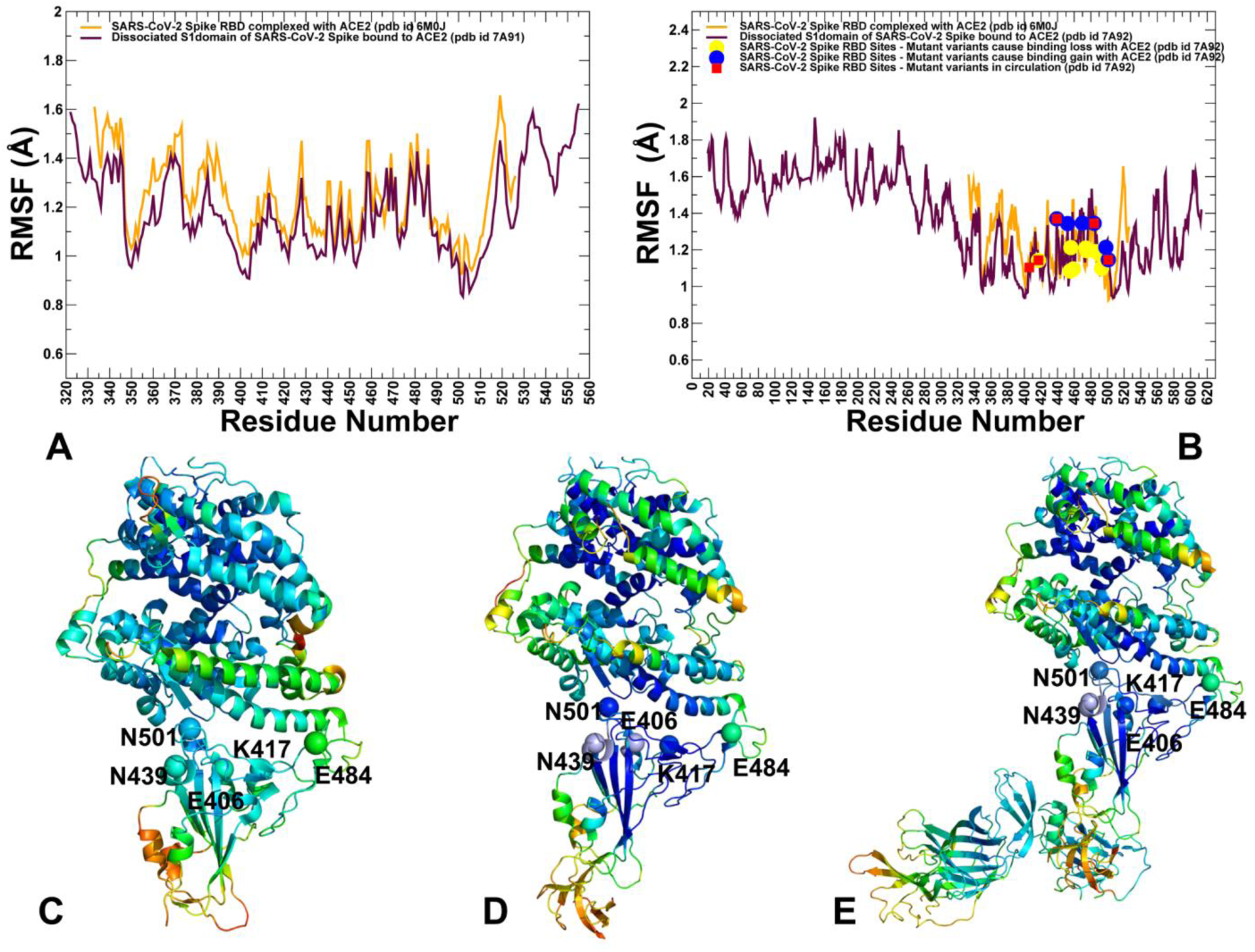
CABS-GG conformational dynamics of the SARS-CoV-2 S-RBD and S1 domain complexes with ACE2. (A) The root mean square fluctuations (RMSF) profiles from simulations of the structures of the SARS-CoV-2 S-RBD complex with ACE2, pdb id 6M0J (in orange lines) and S1 domain complex with ACE2, pdb id 7A91 (in maroon lines). (B) The RMSF profiles from simulations of the structures of the SARS-CoV-2 S-RBD complex with ACE2, pdb id 6M0J (in orange lines) and complete S1 domain complex with ACE2, pdb id 7A92 (in maroon lines). The S-RBD sites L455, F456, S459, Q474, A475, F486, F490, Q493 and P499 whose mutations can abolish binding affinity with ACE2 are shown in yellow filled circles. The S-RBD sites N439, L452, E484/P470, Q498, and N501 whose mutations could enhance binding affinity for ACE2 are shown in blue filled circles. The S-RBD sites E406, N439, K417, E484, and N501 targeted by novel circulating mutations and antibody-escaping mutations are highlighted in red filled squares. (C) Structural mapping of the conformational dynamics profiles in the SARS-CoV-2 S –RBD complex with ACE2 (pdb id 6M0J). A ribbon-based protein representation is used with coloring (blue-to-red) according to the protein residue motilities (from more rigid-blue regions to more flexible-red regions). (D,E) Structural mapping of the conformational dynamics profiles in the S1 domain complexes with ACE2 (pdb id 7A91, 7A92 respectively). The stability profile for protein residues is shown as in (C) using a coloring spectrum from blue to red to highlight changes from rigid to more flexible regions.

Structural maps of the conformational dynamics profiles highlighted these observations showing a more uniform and broad stabilization of the RBD regions in the complex formed by the dissociated S1 domain (Figure 2C-E). Although the overall fluctuation profile of the RBM residues remained largely unchanged, we noticed small but important differences pointing to the greater stability of the 495-YGFQPTNG-502 loop in the ACE2 complex with the dissociated S1 domain. These observations are consistent with the latest structural studies showing that ACE2 binding can induce disassembly of the SARS-CoV-2 S trimer and promote formation of a stable dissociated monomeric S1 complex with ACE2 receptor.^147^ At the same time, a modest mobility of the RBM positions N439, L452, T470, E484, Q498 and N501 was seen in the S1-ACE2 complex (Figure 2B). Importantly, several of these positions N49, E484, and N501 correspond to sites that confer mutational variants with the increased binding to ACE2 and elevated level of transmission and infectivity. The observed partial flexibility of this SARS-CoV-2 RBM motif in the complex formed by the dissociated S1 domain may allow for tolerance and adaptability of these sites to specific modifications resulting in the improved binding affinity with ACE2.

We also report the relative solvent accessibility (RSA) ratio in the SARS-CoV-2 RBD and S1 domain complexes with ACE2 that were obtained by averaging the SASA computations over the simulation trajectories (Supporting Information, Figure S1). The anti-parallel β-sheet regions in the SARS-CoV-2 (residues 451-454 and 492-495) are deeply buried at the interface. The key RBM residues in the central segment (K417, L456, F456, Y473, F490 and Q493) also showed small RSA values, indicating that these positions are buried in the ACE2 complex. Of particular interest were the average RSA values for functional sites E406, K417, N439, E484, and N501 that are central focus of our investigation. We found that E406, K417 and N439 showed moderate RSA values (∼20-30%) indicating that these positions could retain a certain degree of plasticity in the RBD and S1 domain complexes with ACE2 (Supporting Information, Figure S1). The more extreme cases were exemplified by E484 that maintains significant solvent exposure (RSA ∼ 65%) in the ACE2 complexes, while N501 is largely buried with very small RSA values (Supporting Information, Figure S1). However, conformational dynamics profiles indicated that N501 may still maintain some level of plasticity in the ACE2 complex.

To compare the differences in the local flexibility with experimental functional data, we specifically analyzed a group of SARS-CoV-2 RBD residues L455, F456, S459, Q474, A475, F486, F490, Q493 and P499 whose mutations to their SARS-CoV RBD counterpart positions resulted in the abolished binding affinity.^37^ It could be noticed that in the S1 domain complex with ACE2 these residues become appreciably more stable than residues from another group (N439, L452, T470, E484, Q498, N501) that are more susceptible to affinity-improving mutations. Functional studies showed that N439/R426, L452/K439, T470/N457, E484/P470, Q498/Y484 and N501/T487 modifications of these SARS-CoV-2 RBD residues to their respective position in SARS-CoV-RBD can in fact result in the enhanced binding affinity for ACE2.^37^ Our analysis allowed to capture these subtle differences showing that this group of RBD residues may experience larger fluctuations (Figure 2A,B). Interestingly, these differences become more evident only in the ACE2 complex with the dissociated S1 domain, suggesting that the partial redistribution of mobility in the S1-ACE2 complex could provide more room for structural adaptation of N439, E484, and N501 positions (Figure 2B). Hence, a moderate level of residual fluctuations can be preserved even when RBM residues are involved in strong stabilizing contacts with ACE2. Although E484 interacts with the K31 interaction hotspot residue of hACE2, this residue retains a more significant degree of mobility and plasticity in the RBM region which may be associated with the mutational variability and emergence of E484K variant that can improve binding affinity with the host receptor. Interestingly, we found that escape mutations and variants improving binding affinity with the ACE receptor may emerge in sites that are moderately flexible in the S1-ACE2 complex. This suggested that although some of these positions such as K417 and N501 are involved in multiple contacts with ACE2 there should be a substantial energetic plasticity in the interaction network. According to our findings, there may be more room for tolerant modifications of N439 and E484 positions, while potential for favorable mutations at K417 and N501 sites could be more limited.

To summarize, the central finding of this analysis is that conformational dynamics profile for the SARS-CoV-2 S-RBD residues remained largely conserved among ACE2 complexes of the S1-RBD and fully dissociated S1 domain. Another important observation is a consistent trend for moderate residual mobility of RBD residues whose mutations may often lead to the enhanced binding with ACE2. Conformational dynamics analysis also indicated that RBM residues targeted by novel mutational variants may be adaptable and display a range of flexibility – from more dynamic positions at N439 and E486 to more constrained K417 and N501 residues.

### Essential Dynamics of the SARS-CoV-2 S Complexes with ACE2 and the REGN-COV2 Cocktail Unveils Regulatory Roles of Functional Sites Targeted by Mutational Variants

To characterize collective motions and determine the distribution of hinge regions in the SARS-CoV-2 S-RBD and SARS-CoV-2 S1 domain complexes with ACE2 (Figure 3) and REGN-COV2 antibody combination (Figure 4), we performed PCA of trajectories derived from CABS-CG simulations and also determined the essential slow modes using ENM analysis. The reported functional dynamics profiles were averaged over the first three major low frequency modes. For comparison of ACE2-induced changes in the functional dynamics, we first characterized the slow mode profiles for the unbound forms of the RBD and S1 domain extracted from the crystal structures of the complexes (Figure 3A-C). For the unbound RBD structure the local minima associated with local hinge points corresponded to W353, F374, F400, L452, R466, Q493 and V510 residues. Some of these residues L452 and Q493 are involved in the interactions in the complex and their immobilized hinge position in the unbound form may be important to induce the optimal inter-molecular association with ACE2. Interestingly, none of the functional positions targeted by novel mutational variants that promote infectivity and antibody resistance (E409, K417, N439, E484, and N501) corresponded to hinge positions in the unbound RBD form. In fact, E484 residue is located in the moving region of the unbound RBD structure in slow modes (Figure 3A). The unbound form of the dissociated S1 domain featured local hinge positions in residues F518 and V539, C538 and F592 (Figure 3B,C). Strikingly, these findings are consistent with our recent analysis of collective dynamics in the open and closed forms of the SARS-CoV-2 S trimer structures showing that these residues correspond to major regulatory centers of functional motions and coordinate global displacements of the S1 regions with respect to more rigid S2 subunit in distinct timer states.^148^

**Figure 3.**
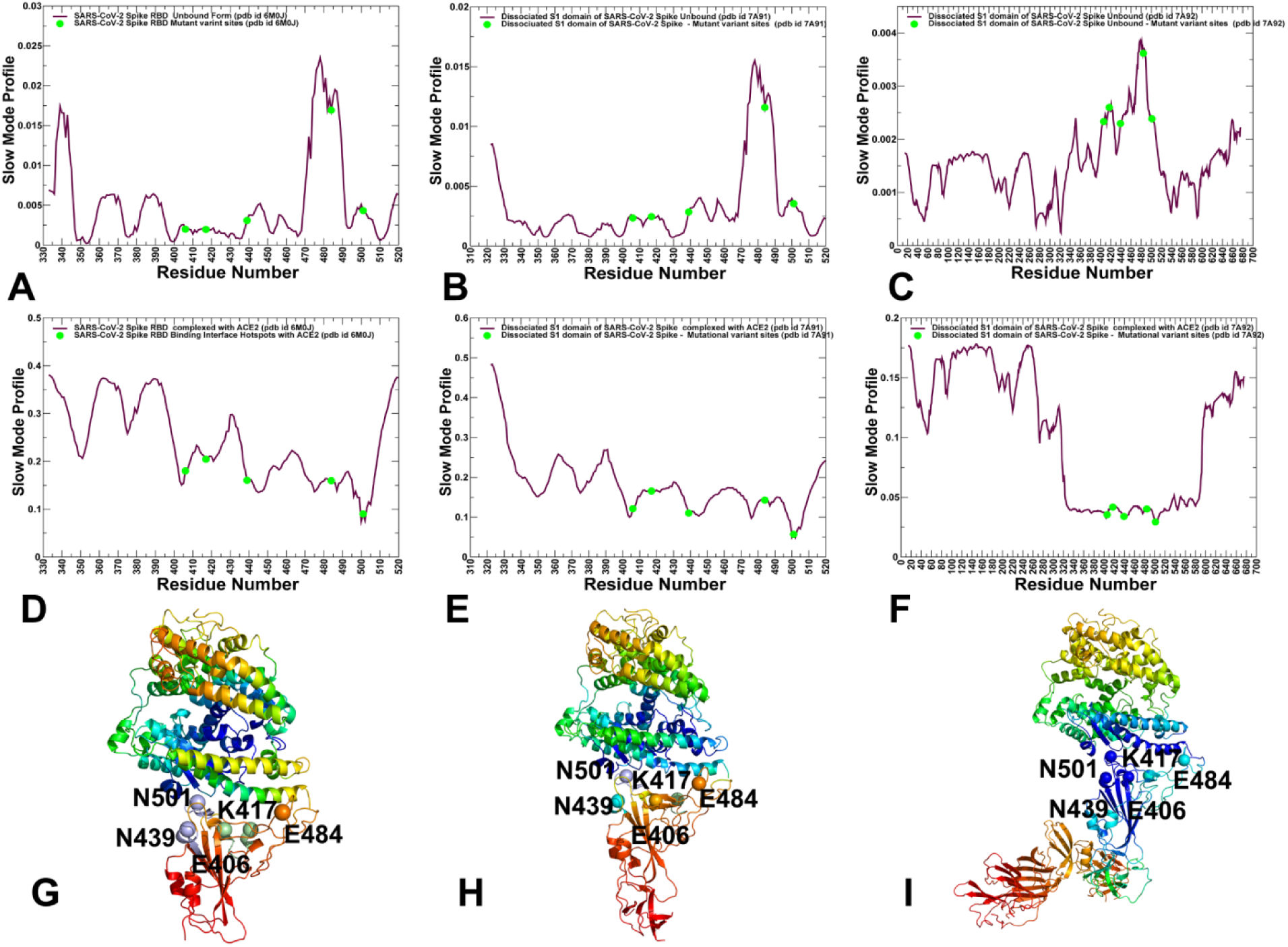
Functional dynamics of the SARS-CoV-2 S-RBD and S1 domain complexes with ACE2. The mean square displacements in functional motions are averaged over the three lowest frequency modes. The essential mobility profiles for the unbound forms of the SARS-CoV-2 S-RBD (A) and dissociated forms of the S1 domain (B,C). (D) The slow mode profile of the bound SARS-CoV-2 S-RBD structure in the complex with ACE2 (pdb id 6M0J). (E,F) The slow mode profiles of the dissociated S1 domain complexed with ACE2. The essential mobility profiles are shown in maroon lines and positions of key functional residues E406, K417, N439, E484, and N501 are highlighted by filled green circles. Structural maps of the essential mobility profiles for the SARS-CoV-2 S-RBD complex with ACE2 (G) and complexes formed by the dissociated S1 domain with ACE2 (H,I). Structural maps of collective dynamics are derived from fluctuations driven by the slowest three modes. The color gradient from blue to red indicates the decreasing structural stability (or increasing conformational mobility) of protein residues. The key functional residues E406, K417, N439, E484, and N501 are shown in spheres colored according to the level of mobility in the low frequency slow modes.

**Figure 4.**
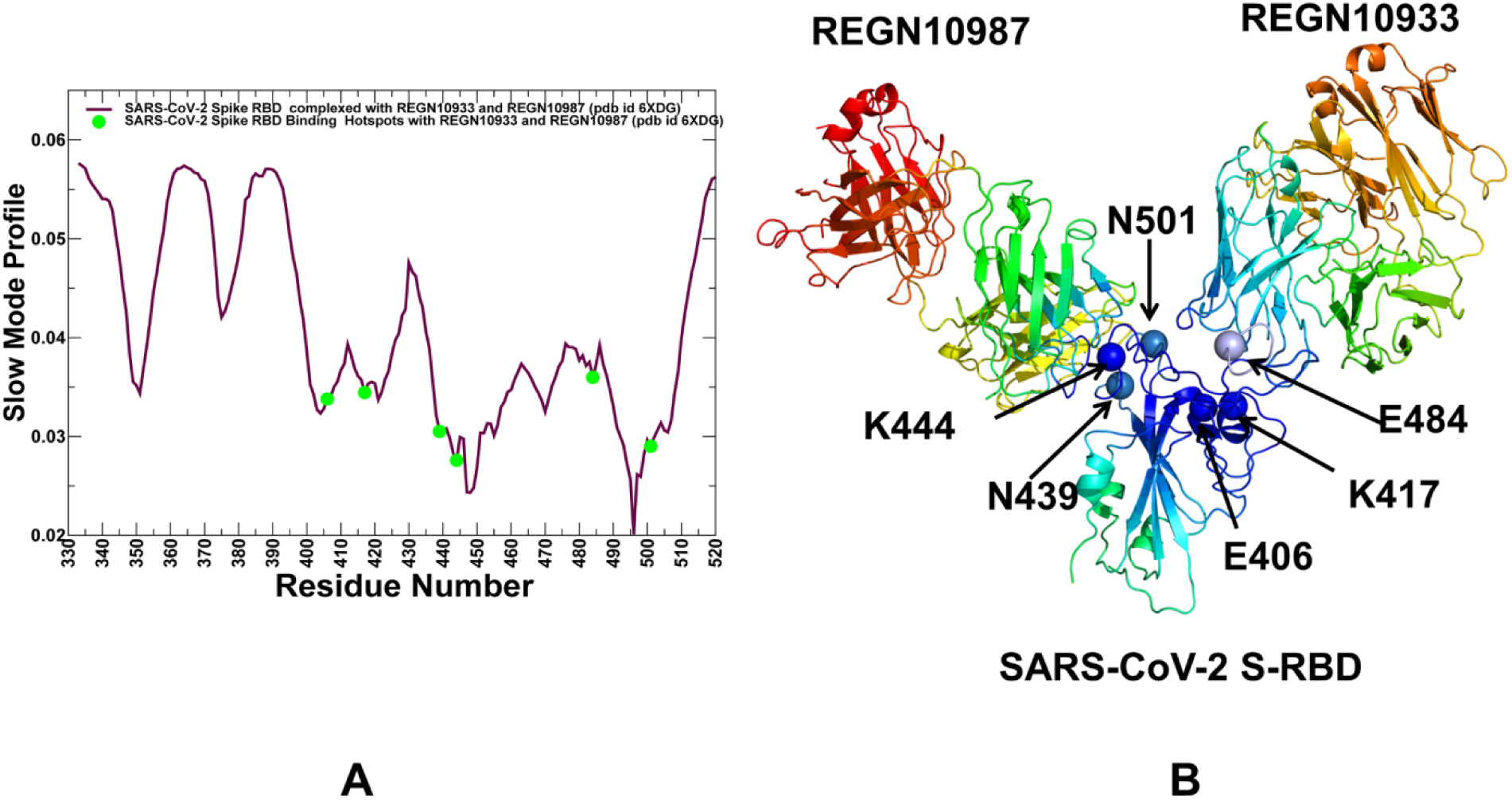
Functional dynamics of the SARS-CoV-2 S-RBD complex with REGN-COV2 antibody cocktail (REGN10933 and REGN10987). The mean square displacements in functional motions are averaged over the three lowest frequency modes. (A) The slow mode profile of the bound SARS-CoV-2 S-RBD structure in the complex with REGN-COV2 antibody combination (pdb id 6XDG). The essential mobility profiles are shown in maroon lines and positions of E406, K417, N439, E484, and N501 are highlighted by filled green circles. (B) Structural map of the essential mobility profiles for the SARS-CoV-2 S-RBD complex with REGN-COV2 antibody cocktail. The color gradient from blue to red indicates the decreasing structural stability (or increasing conformational mobility) of protein residues. The key functional residues E406, K417, N439, E484, and N501 are annotated and shown in colored spheres according to the level of their respective mobility in the low frequency slow modes.

Consistent with these studies, the functional movements of RBDs can be determined by the main hinge centers located near F318, S591, F592, V539 residues. These results highlighted the conserved nature of global hinges in the dissociated S1 domain and in the SARS-CoV-2 S complete trimer. Notably, the RBD residues targeted by novel mutational variants are located in the flexible moving regions of the unbound S1 domain. The distribution of hinge sites is altered in the SARS-CoV-2 RBD complex with ACE2, and strikingly the sites subjected to circulating variants and escape mutations often coincided with hinge clusters anchored by E406, K417, N439 and N501 residues (Figure 3D). E484 is located near F486 that is aligned with another local hinge position, while N501 together with T500 and Y505 can form a dominant hinge center in the complex. Hence, a group of functional residues that include N439, F486, T500, N501 and Y505 sites may form a network of regulatory centers coordinating global movements of the RBD and ACE2 molecules. These findings are in line with studies of conformational dynamics of the SARS-CoV-2 trimeric spike glycoprotein in complex with ACE2 revealing these positions may be involved in regulation of continuous swing motions of ACE2-RBD relative to the SARS-CoV-2 S trimer.^33^

The slow mode profiles obtained for complexes of the dissociated S1 domain with ACE2 further highlighted the role of these functional residues in collective motions (Figure 3E,F). The entire RBD region and structurally conserved C-terminal domain 1, CTD1 (residues 528-591) become largely immobilized in the S1 domain complex with ACE2 complex, while C-terminal domain 2, CTD2 region (residues 592-686) can undergo large movements (Figure 3F). By zooming on the RBD regions, one could see that local minima and corresponding hinge sites are almost precisely aligned with residues E406, K417, N501 and Y505. Hence, the RBD residues targeted by mutant variants may play an important role in coordinating the relative orientation and approach angle of the ACE2 receptor in the complex, and consequently affect recognition and signal transmission in the functional complex. Structural maps of functional dynamics profiles illustrated these findings showing that the RBD regions that include functional positions E406, N439 and N501 are aligned with immobilized in slow motions hinge centers, while K417 and E484 residues could be less constrained during collective movements (Figure 3G-I). The central finding of this analysis is the unique role that positions targeted by novel variants (N439, N501) and escape mutations (E406) could play in concerted functional movements of S-RBD and S1 domain when bound to ACE2. Based on the results we argue that functional role of these sites in controlling global motions and long-range interactions in ACE2 complexes could be an important reason for mediating escape from antibody binding while maintaining and enhancing binding with the host receptor.

To compare the slow mode profiles of the SARS-CoV-2 S1/RBD complexes with ACE2 and neutralizing antibodies, we performed ENM-based modeling of slow mode profiles in the S-RBD complex with REGN-COV2 cocktail of two antibodies REGN-10933 and REGN10987 (Figure 4). This analysis showed that sites E406 and K417 corresponded to local hinge positions, while N439 and K444/G446 residues are now aligned with the dominant hinge center of the SARS-CoV-2 RBD complex with the REGN-COV2 cocktail (Figure 4A).

Importantly, antibody binding altered dynamic role of residues E484 and N501 that become aligned with moving regions in the collective dynamics of the complex (Figure 4A). Structural mapping of the essential profiles further illustrated this point, showing that functionally immobilized in collective motions hinge centers are localized near E406, N439 and K444 sites, while E484 and N501 positions could undergo some movements in the complex (Figure 4B). In this context it is particularly interesting to compare our observations with functional studies showing that K417 and F486 are sites of escape from RERGN10933, while mutations in K444 and G446 escape neutralization by REGN10987 and E406 is a unique site susceptible to mutations escaping both antibodies.^70^ In line with these experiments, we found that E406 and K444 positions may correspond to the antibody-specific unique hinge centers of collective motions that control relative orientation and rigid body movements of REGN10933 and REGN10987 molecules (Figure 4B). This is in some contrast to SARS-CoV-2 RBD complexes with ACE2 in which N439 and N501 form the major hinge center of functional dynamics. As a result, it is possible that mutations in K444 and E406 positions may perturb not only local interactions with antibody molecules but alter the global collective movements and long-range communication which may be sufficient to trigger mutational escape from antibody binding. At the same time, these mutations could only moderately change the SARS-CoV-2 RBD local interactions with ACE2 without affecting the collective movements in the complex.

To summarize, this analysis suggested that mutational variants and escape mutations may preferentially target specific positions involved in regulation and coordination of functional dynamics motions and allosteric changes in the SARS-CoV-2 complexes with ACE2 and REGN-COV2 antibody cocktail.

### Mutational Sensitivity Profiling of the SARS-CoV-2 RBD Binding Interfaces Reveals Energetic Plasticity in Sites Susceptible to Circulating Mutational Variants and Antibody Escaping Modifications

We first performed a systematic alanine scanning of the SARS-CoV RBD S protein residues (Figure 5A) and residues from the dissociated S1 domain in the complexes with the ACE2 host receptor (Figure 5B,C). Using the equilibrium ensembles obtained from simulation trajectories, we evaluated the average cumulative mutational effect of alanine substitutions on protein stability and binding affinity with the host receptor. The alanine scanning of the SARS-CoV-2 RBD residues highlighted a significant destabilization effect caused by mutations of G446, Y453, L455, F456, F486, Y489, Y495, T500, and Y505 residues (Figure 5A). These residues also corresponded to the binding free energy hotspots in the complexes formed by the dissociated S1 domain with ACE2 (Figure 5B,C). In particular, large destabilization effects were observed upon mutations of Y453, L455, F456, Y489, and F490 residues in the S1-ACE2 complexes. Notably, the largest destabilization changes were produced by alanine mutations of F456 and Y489 residues, displaying clear and pronounced peaks of the profile and pointing to these positions as key binding affinity hotspots in the S1-ACE2 complexes. (Figure 5B,C). Several key binding energy hotspot sites (Y453, Y489, and Y505) are conserved between SARS-CoV and SARS-CoV-2 proteins and are located in the central segment of the interface. A detailed analysis of the intermolecular contacts in the SARS-CoV-2 RBD and S1 complexes with ACE2 aided in understanding of the binding energy preferences of RBD residues (Supporting Information, Tables S1-S3). This analysis is particularly instructive by considering contact distributions with two virus-binding hotspots on ACE2 formed by interacting residues K31 and E35 as well as K353 and D38.

**Figure 5.**
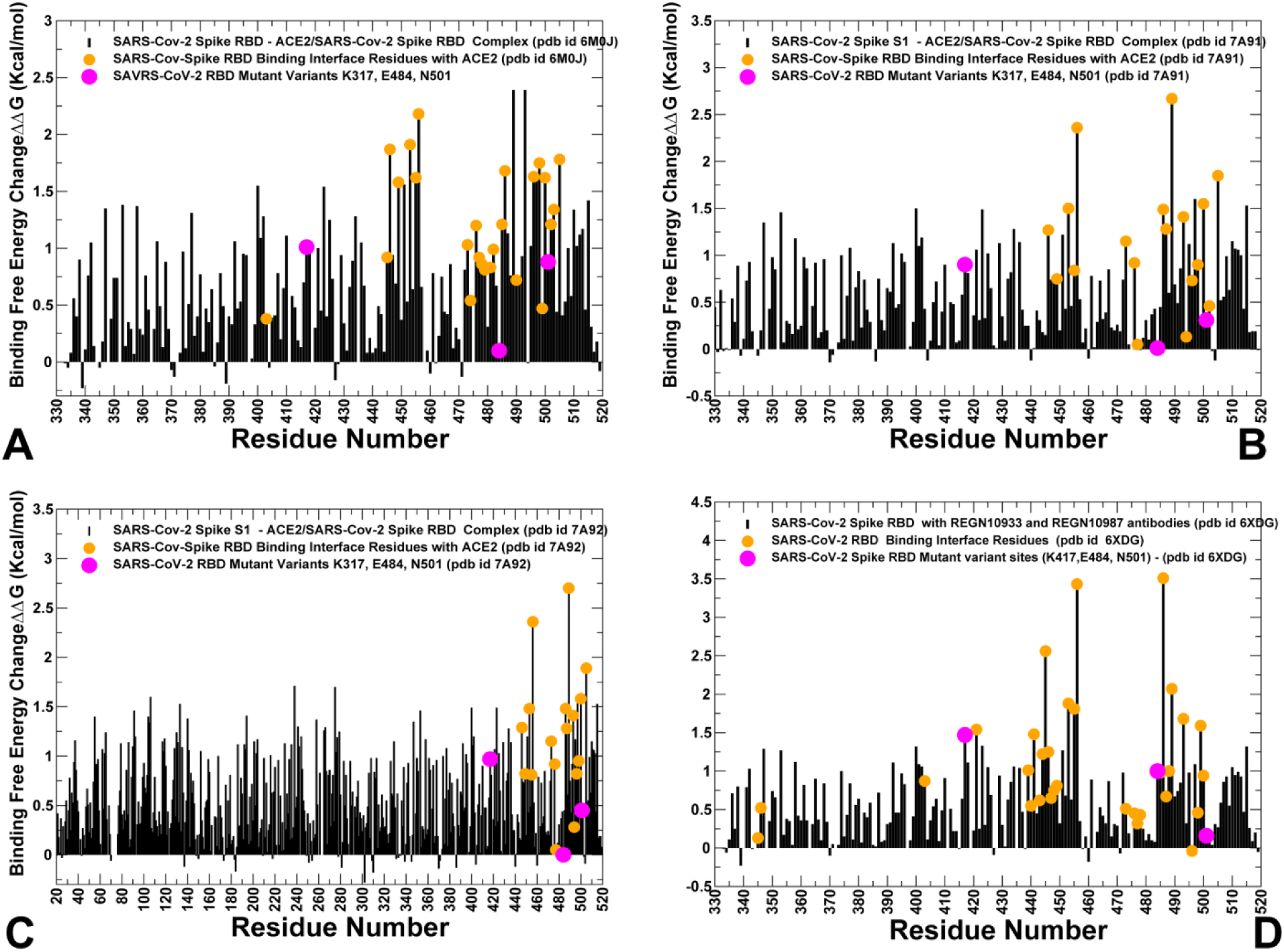
Alanine scanning of the RBD residues in the SARS-CoV-2 S-RBD and S1 domain complexes with ACE2 and REGN-COV2 antibody cocktail. (A) The binding free energy changes upon alanine mutations for the RBD residues in the SARS-CoV-2 S-RBD complex with ACE2 (pdb id 6M0J). (B) The binding free energy changes upon alanine mutations for the S1-RBD residues in the S1 domain complex with ACE2 (pdb id 7A91). (C) The binding free energy changes upon alanine mutations for the dissociated S1 domain residues in the complex with ACE2 (pdb id 7A92). (D) The binding free energy changes upon alanine mutations for the SARS-CoV-2 S-RBD residues in the complex with REGN-COV2 cocktail (pdb id 6XDG). The binding energy changes for the protein residues are shown in maroon bars. The binding interface residues are depicted in orange filled circles and functional residues K417, E484 and N501 targeted by mutational variants are highlighted in magenta filled circles.

In particular, Y489 residue makes numerous favorable contacts with multiple ACE2 residues K31, F28, Y873, L79 and T27 residues, while another hotspot position F456 forms interactions with T27, D30 and K31 positions on ACE2 (Supporting Information, Tables S1-S3). L455 residue of the RBD makes favorable contacts with the key hotspots on ACE2 K31, H34, and D30 while Q493 contacts K31, H34 and E35 positions. The hydrophobic residue F486 forms multiple interactions with M82, L79, Y83, Q24, while another hydrophobic RBD site F490 interacts with K31 and Y473 with T27 (Supporting Information, Tables S1-S3). Overall, the alanine scanning highlighted the importance of the SARS-CoV-2 RBD interactions formed by the middle segment of the RBM interface (K417, Y453, L455, F456, Y489 and Q493) as mutations of these residues resulted in a significant loss of binding affinity (Figure 5A-C). These results are consistent with recent functional studies, indicating that mutations of the SARS-CoV-2 RBD residues (L455/Y442, F456/L443, F486/L472, F490/W476, Q493/N479) result in a significant reduction of their binding affinity with ACE2.^37^

The alanine scanning of the SARS-CoV-2 RBD residues in the complex with REGN-COV2 antibody cocktail revealed large destabilization effects and strong peaks at positions K444, V445, F456, F486, and Y489 ((Supporting Information, Table S4, Figure 5D). The largest free energy changes exceeding 3 kcal/mol were observed for alanine modifications of F456 and F486 residues which is consistent with the prominent role these residues play in eliciting antibody-escaping mutations.^70^ We also specifically highlighted binding free energy changes caused by alanine modifications in sites targeted by circulating and antibody-escaping variants E406, K417, N439, E484, and N501. The results showed that alanine substitutions in these positions induced only minor destabilization changes in the RBD-ACE2 and S1-ACE2 complexes, and these values were particularly small when mutations were introduced in E484 and N501 positions (Figure 5A-C). A slightly different pattern was seen in the RBD-REGN-COV2 complex where alanine modifications in K417, N439 and E484 residues led to appreciable > 1.0 kcal/mol binding free energy loss, while mutations in E406 and N501 positions produced only small destabilization effect. These patterns indicated that antibody binding may induce changes in the binding interactions and distribution of the binding energy hotspots.

To further quantify the effects of the binding energy hotspots, we followed up with a complete mutational sensitivity analysis of these RBD positions (Figure 6). The profiling showed that all mutations in Y453, F456, F486 and Y505 positions were highly destabilizing (Figure 6A-D), while for L455 and F490 positions the loss of binding affinity was only moderately destabilizing for majority of substitutions (Figure 6E,F). The destabilization pattern observed for all modifications of F456 and F486 residues is consistent with the functional experiments^37^ highlighting the importance of these positions in binding affinity. We also conducted mutational sensitivity scanning of K417, E484 and N501 residues that are targeted by circulating variants in the UK (B.1.1.7/501Y.V1), South Africa (501Y.V2) and Brazil (B1.1.28/501.V3) lineages^59–65^ as well as profiling of E406 and N439 sites that are of considerable interest due newly emerging circulating variants and antibody-escaping mutations (Figure 7). Of special interest was analysis of the protein stability and binding free energy changes incurred by N501Y mutation that is prominently featured in the UK B.1.1.7 variant and mutations K417N, E484K that together with N501Y modifications are central to the increased transmission and infectivity effects seen in the South Africa (501Y.V2) and Brazil (B1.1.28/501.V3) lineages. In addition, we considered other important sites of newly circulating mutations N439 and the unique position E406 giving rise to unique antibody-escaping mutations.^70, 81^

**Figure 6.**
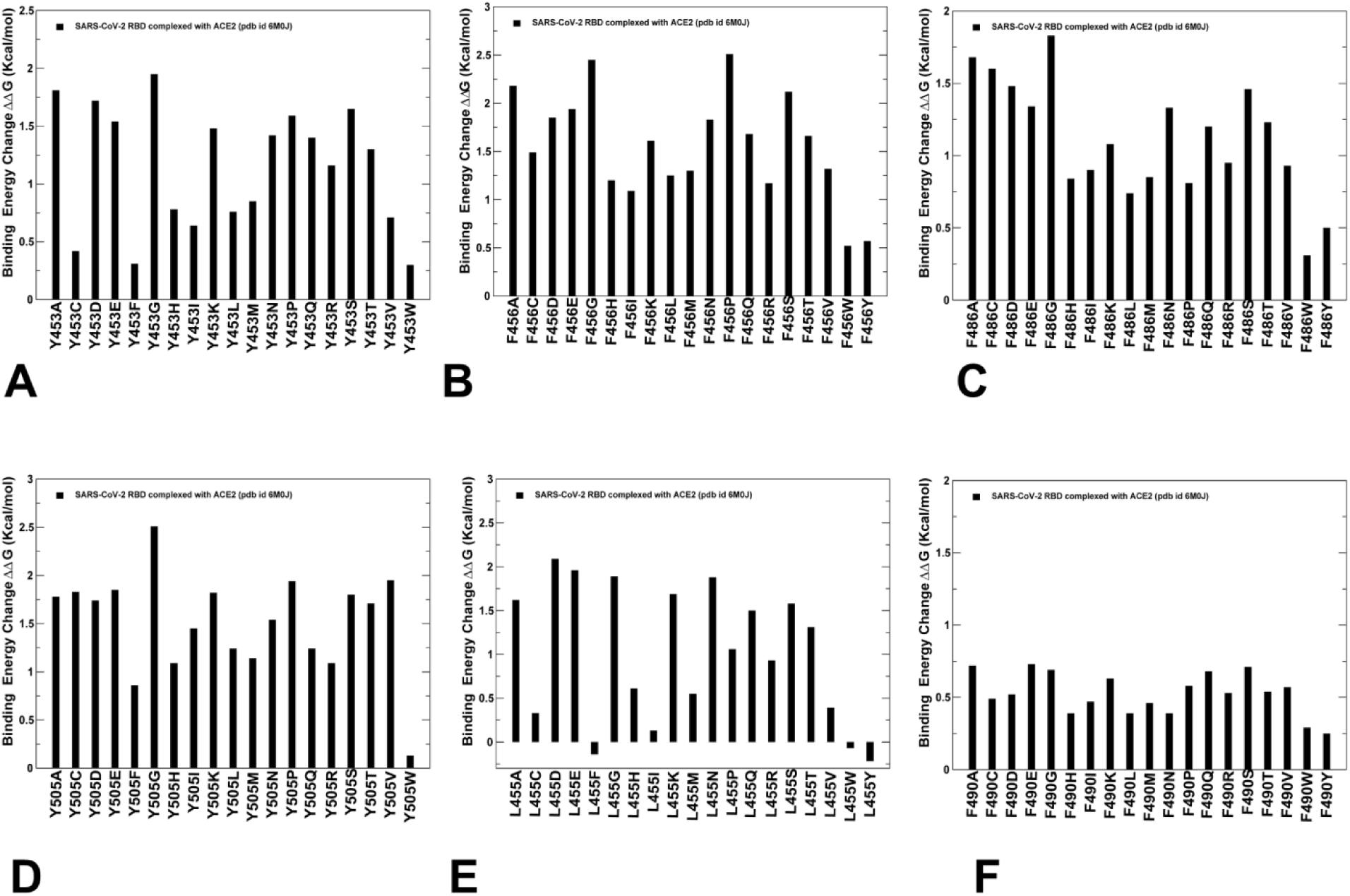
Mutational sensitivity analysis of binding free energy hotspots in the SARS-CoV-2 S-RBD complex with ACE2 (pdb id 6M0J). (A) Mutational sensitivity scanning of Y453 residue. (B) Mutational sensitivity scanning of F456 residue. (C) Mutational sensitivity scanning of F486 residue. (D) Mutational sensitivity scanning of Y505 residue. (E) Mutational sensitivity scanning of L455 residue. (F) Mutational sensitivity scanning of F490 residue. The protein stability changes are shown in maroon-colored filled bars.

**Figure 7.**
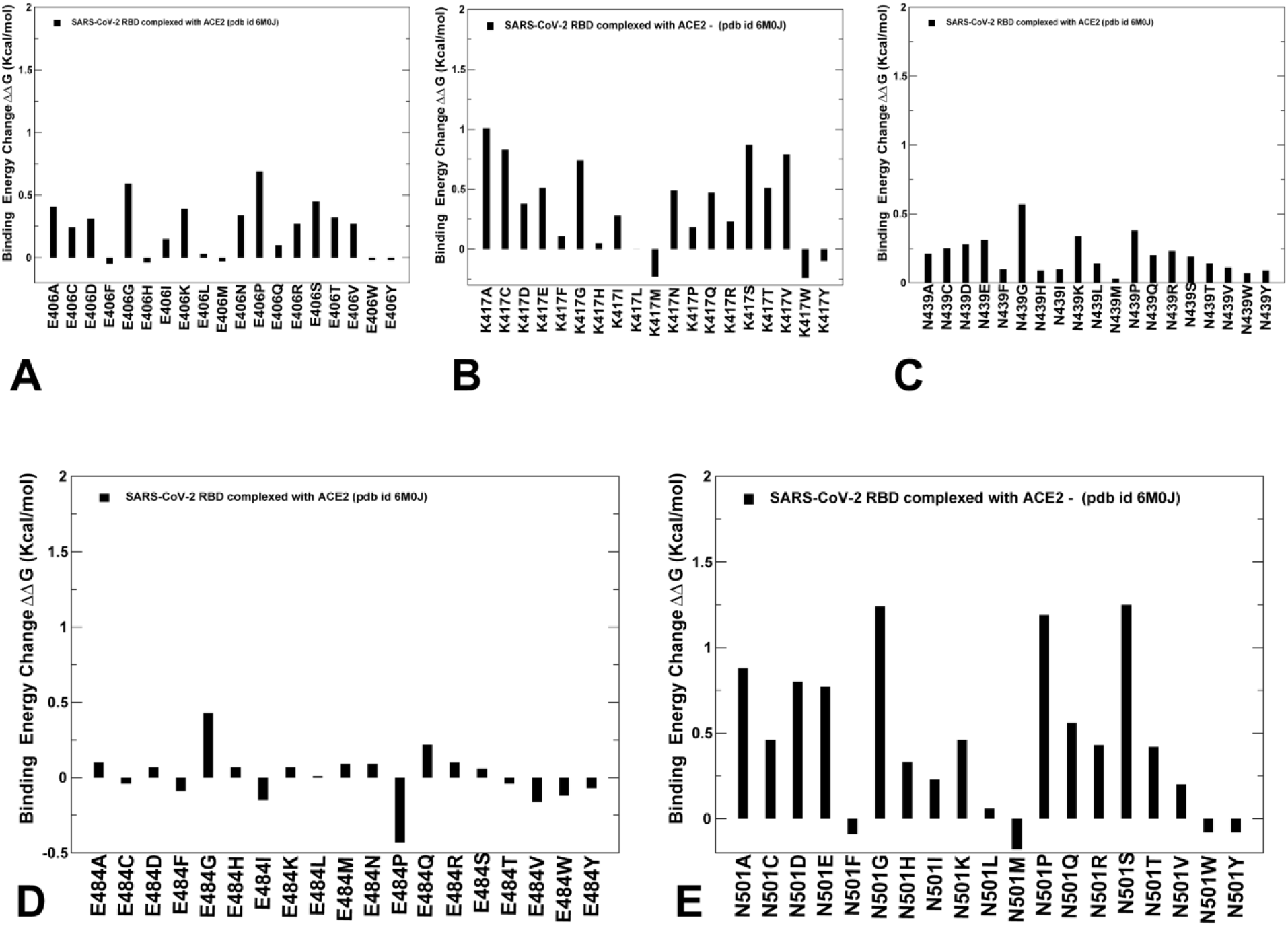
Mutational sensitivity analysis of functional RBD residues targeted by novel mutational variants and antibody-escaping mutations in the SARS-CoV-2 S-RBD complex with ACE2 (pdb id 6M0J). (A) Mutational sensitivity scanning of E406 residue. (B) Mutational sensitivity scanning of K417 residue. (C) Mutational sensitivity scanning of N439 residue. (D) Mutational sensitivity scanning of E484 residue. (E) Mutational sensitivity scanning of N501 residue. The protein stability changes are shown in maroon-colored filled bars.

Consistent with the deep mutational scanning experiments^35, 66^ we found that E406, N439, and E484 sites are energetically adaptable and can effectively tolerate different mutations without incurring significant changes in protein stability and binding affinity (Figure 7 A,C,D). Somewhat larger but still relatively tolerable were binding free energy changes induced by mutations in K417 and N501 positions (Figure 7B,E). K417 is a unique ACE2-interacting residue that forms favorable contacts with central residues of the ACE2 interface H34 and D30 (Supporting Information, Table S1-S3). However, an appreciable energetic plasticity could be seen in mutational sensitivity profiling of K417 residue (Figure 7B). Although K417 mutations to alanine or glycine produced fairly significant destabilization changes, K417N and K417D mutations led to only small perturbations (∼0.4-0.5 kcal/mol). Indeed, deep mutational scanning suggests that the K417N mutation has minimal impact on binding affinity with ACE2.^35^

The results also predicted the marginal improvement in the binding free energy mediated by E484K mutation and only very modest increase in the binding affinity upon K417N modification (Figure 7D). The experimental studies indicated that the E484K mutation may induce a moderate improvement in binding affinity and showed that other single mutations of E484 may only slightly compromise spike folding stability and binding affinity for ACE2.^35, 70^ According to our analysis, several hydrophobic substitutions in this position (E484I, E484V, E484F, E484W, and E484P) may in fact lead to the moderately improved affinity, while other mutations appeared to produce only marginal destabilization (Figure 7D). These results indicated a significant plasticity of this important RBD position that is relatively exposed and may favor hydrophobic residues in this position to improve both stability and binding. The mutational sensitivity profiling at N501 position is consistent with deep mutational scanning experiments^35, 66^ reproducing the improvements in binding mediated by N501F and N501Y mutations (Figure 7E). Indeed, deep mutation scanning showed that N501F, N501T and N501Y mutations may lead to moderate enhancement of binding with ACE2, while N501D is an affinity-decreasing mutation.^35^ We observed only a small destabilization effect for N501T and more significant destabilization upon N501D and N501A/G mutations (Figure 7E). Importantly, these results supported the notion that N501Y mutational variant could be beneficial for ACE2 binding, while escaping neutralizing antibodies targeting the same region.

We also performed mutational sensitivity analysis of the key functional positions in the SARS-CoV-2 RBD complex the REGN-COV2 (Figure 8A-D). The results revealed moderate changes upon mutations at E406, E484, N501 and N439 positions. Interestingly, although E406W escaped both individual REGN-COV2 antibodies,^70^ our results indicate that this mutation would not drastically perturb the RBD region and significantly affect binding interactions with the REGN-COV2 antibody cocktail (Figure 8A). Hence, antibody-escaping effect of E406W substitution may not be trivially linked to the local interaction effects. In this context, given the results of functional dynamics analysis, it is tempting to argue that mutations at E406 position may instead alter collective movements and compromise long-range allosteric couplings in the SARS-CoV-2 RBD. This may ultimately affect neutralization activity of the REGN-COV2 antibody combination. At the same time, a wide range of modifications at K417, K444 and F486 sites resulted in significant destabilization changes and loss of the binding affinity (Figure 8E-G). These results are in excellent agreement with functional mapping of the SARS-CoV-2 RBD residues that affect binding of the REGN-COV2 cocktail showing that F486 mutations are predominant for escaping neutralization by REGN10933 and mutations at K444 evade binding of REGN10987 antibody.^70^

**Figure 8.**
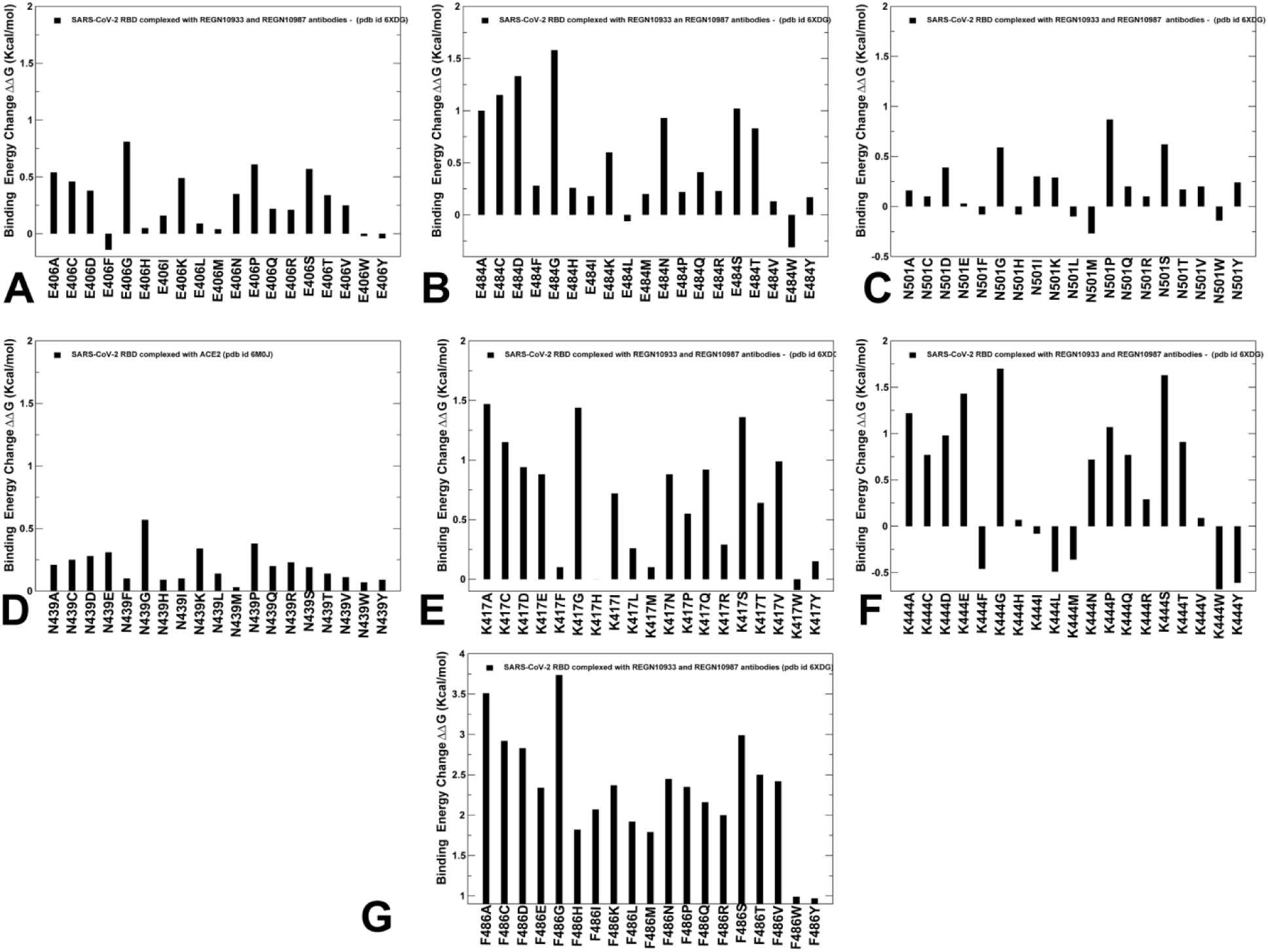
Mutational sensitivity analysis of functional RBD residues in the SARS-CoV-2 S-RBD complex with REGN-COV2 antibody cocktail (REGN10933 and REGN10987). (A) Mutational sensitivity scanning of E406 residue. (B) Mutational sensitivity scanning of E484 residue. (C) Mutational sensitivity scanning of N501 residue. (D) Mutational sensitivity scanning of N439 residue. (E) Mutational sensitivity scanning of K417 residue. (F) Mutational sensitivity scanning of K444 residue. (G) (E) Mutational sensitivity scanning of F486 residue. The protein stability changes are shown in maroon-colored filled bars.

Consistent with the functional analysis of the immune-selected mutational landscape in the S protein, we found that a wide spectrum of K444 modifications induced a significant loss of binding free energy (Figure 8E) including K444E and K444N mutations that showed a broad-range resistance against multiple antibodies.^69^ The large destabilization changes caused by F486 mutations can be contrasted to fairly small changes incurred by E484 mutations, indicating that E484 site is characterized by sufficient level of structural plasticity and energetic adaptability to readily accommodate mutations in complexes with ACE2 and REGN-COV2 cocktail. These findings may explain why single-site mutations of these residues can only slightly change binding affinity for ACE2 and folding stability, while double-site mutations of proximal E484 and F486 can significantly weaken the fitness of the SARS-CoV-2 RBD region and binding.^66^ To summarize, our results pointed to several interesting trends. First, mutational sensitivity profiling of the conserved hydrophobic binding energy hotspots Y453, L455, F456, F486, and Y505 consistently yielded large destabilization changes affecting folding stability and binding to ACE2 receptor, making these positions unlikely candidates for antibody escaping mutations as even small modifications in these positions could have a severe detrimental effect on the spike activity. Second, we found that SARS-CoV-2 binding affinity could be strongly influenced by the virus-binding hotspot K31 and H34 in the middle of the interface through an extensive interaction network with K417, Y453, L455, F456, and Q493 residues. Finally, mutational analysis of K417, E484 and N501 positions implicated in new mutational strains and antibody-escaping changes showed that these residues correspond to important interacting centers with a significant degree of structural and energetic plasticity. Indeed, N501Y, E484K and K417N mutations can result in the improved or only slightly decreased affinity with ACE2. These results suggest a hypothesis that antibody-escaping mutations target residues with sufficient plasticity and adaptability to preserve a sufficient spike activity while having a more detrimental effect on antibody recognition. These findings are particularly interesting in light of recent functional studies^66^ showing that escape mutations target a subset of sites in the antibody-RBD interfaces corresponding to binding energy hotspots. Importantly, these experiments suggested that escape mutations are consistently those that have significant deleterious effects on antibody binding but little negative impact on ACE2 binding and RBD folding. Based on our findings, we argue that escape mutations constrained by the requirements for ACE2 binding and preservation of RBD stability may preferentially select structurally plastic and energetically adaptable allosteric centers at the key interfacial regions to compromise antibody recognition through modulation of global motions and allosteric interactions in the complex.

### Perturbation Response Scanning Reveals Structurally Adaptable Allosteric Effector Hotspots in Sites Targeted by Global Circulating Mutations

Using the PRS method^124–127^ we quantified the allosteric effect of each residue in the SARS-CoV-2 complexes in response to external perturbations. The effector profiles estimate the propensities of a given residue to influence dynamic changes in other residues and can be applied to identify regulatory hotspots of allosteric interactions as the local maxima along the profile. First, we computed the residue-based effector response profiles for the SARS-CoV-2 RBD complex with ACE2 (Figure 9A) and the complexes formed by the dissociated S1 domain with ACE2 (Figure 9B,C). By comparing the PRS profiles in the ACE2 complexes with SARS-CoV-2 S1/RBD and REGN-COV2 antibody cocktail, we determined the distribution of regulatory allosteric centers and highlighted a potential role of sites targeted by global circulating mutations.

**Figure 9.**
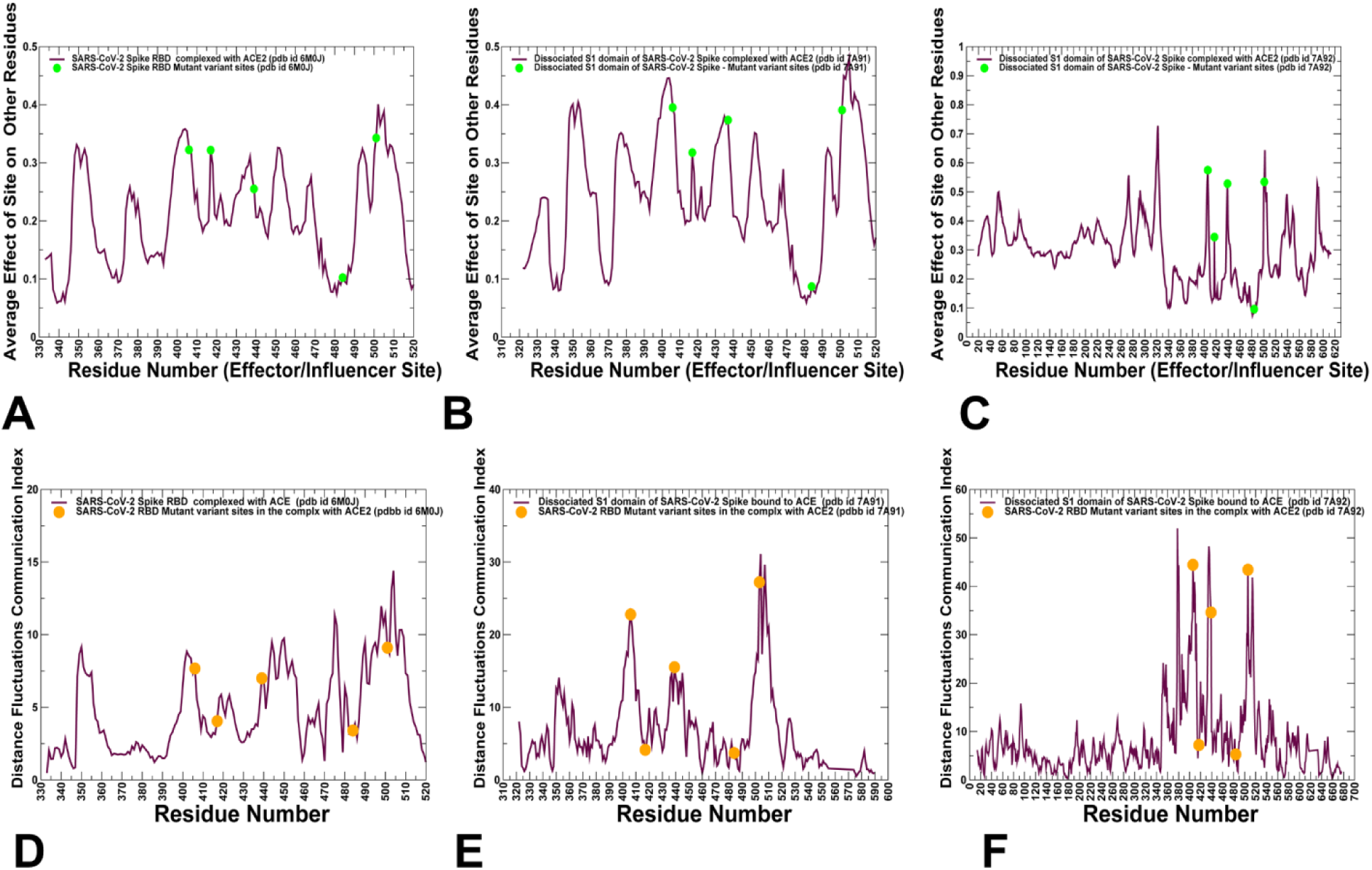
The PRS effector profiles and distance fluctuation communication indexes for the SARS-CoV-2 S-RBD and S1 domain complexes with ACE2. (A) The PRS effector distribution profile for the SARS-CoV-2 S-RBD complex with ACE2 (pdb id 6M0J). (B) The PRS effector profile for the S1-RBD complex with ACE2 (pdb id 7A91). (C) The PRS effector profile for the dissociated S1 domain complex with ACE2 (pdb id 7A92). The positions of functional residues E406, K417, N439, E484, and N501 are indicated by filled green circles. (D) The distance fluctuation communication index profile for the SARS-CoV-2 S-RBD complex with ACE2 (pdb id 6M0J). (B) The distance fluctuation communication index profile for the S1-RBD complex with ACE2 (pdb id 7A91). (C) The distance fluctuation communication index profile for the dissociated S1 domain complex with ACE2 (pdb id 7A92). The positions of functional residues E406, K417, N439, E484, and N501 are indicated by filled orange circles.

Strikingly, the effector profile of the SARS-CoV-2 RBD complex with ACE2 featured two major peaks corresponding to residues E606 and T500/N501, indicating that E406 and N501 positions are aligned with the regulatory centers that may control allosteric communications in the complex (Figure 9A). Several other notable peaks corresponded to W353, K417, N439 and L452 residues. Hence, all known positions targeted by novel circulating mutational variants with the exception of E484 corresponded to the effector peaks and are involved in coordination of allosteric communications in the SARS-CoV-2 RBD complex with ACE2. Moreover, the effector profiles indicated that these regulatory sites may function in a coordinated manner and maintain an allosteric cross-talk to control signal transmission “traffic” and long-range interactions in the RBD-ACE2 complex. Interestingly, several of these effector centers L452, N439 and N501 were among SARS-CoV-2 RBD residues whose mutations to the SARS-CoV RBD counterparts N439/R426, L452/K439, and N501/T487 enhanced the binding affinity.^37^ The prominent role of these residues as regulatory effector centers becomes even more apparent in the S1 domain complexes with ACE2 (Figure 9B,C). It is evident that E406, K417, N439 and especially N501 positions corresponded to sharp peaks of the effector profile. This implies that these sites may be collectively responsible for coordination of long-range communication in the system. The central result of this analysis is that circulating and escape mutations appeared to target residues corresponding to structurally and energetically adaptable regulatory control points that can tolerate individual mutations and often enhance ACE2 binding, while at the same time allowing for coordinated modulation of allosteric communications. We suggest that allosteric signaling in the SARS-CoV-2 RBD complex with ACE2 is adaptable where a mutation of a regulatory control point can be functionally compensated through energetic rebalancing of structurally plastic allosteric hotspots.

Using a protein mechanics-based approach^149–152^ we also employed distance fluctuations analysis of the conformational ensembles to further probe allosteric communication preferences of the RBD residues in the SARS-CoV-2 RBD and S1 complexes with ACE2. The residue-based distance fluctuation communication indexes measure the energy cost of the dynamic residue deformations and could serve as a robust metric for assessment of allosteric propensities of protein residues.^153–155^ In this model, dynamically correlated residues whose effective distances fluctuate with low or moderate intensity are expected to communicate with the higher efficiency than the residues that experience large fluctuations. Notably, structurally stable and densely interconnected residues as well as moderately flexible residues that serve as a source or sink of allosteric signals could feature high value of these indexes.

The distance fluctuation profile of the SARS-CoV-2 RBD and S1 domain complexes with ACE2 showed a small but important redistriubution of major peaks, pointing to sites E406, W436, N439, N501 and Y505 (Figure 9D). Notably, this group of residues is featured prominently among peaks of the profile when the entire S1 domain monomer forms complex with ACE2 (Figure 9E,F). We also noticed that the overall shape and distribution of the peaks are similar between the PRS effector profiles and distance fluctuation communication indexes profiles. Notably, E406, N439 and N501 sites were featured as recurring peaks in both distributions, strengening the proposed notion that positions targted by the emerging mutational variants can cooperate and play a central role in regulation of long-range couplings and allosteric communications in the complexes with the ACE2 host receptor. Hence, the distance fluctuation profiling and analysis of communication indexes provide important supporting evidence to the PRS modeling, suggesting that structurally stable positions and potential allosteric hotspot residues only partially overlap, and allosteric hubs may exhibit a certain degree of structural plasticity and energetic adaptability to enable balance between binding and signaling function. To understand a potential role of the E484 residue, it is instructive to analyze the PRS sensor profile (Supporting Information, Figure S2). A comparison between sensor profiles obtained for the unbound and bound forms of the SARS-CoV-2 RBD showed that E484 position is aligned with the dominant peak of the sensor profile in the unbound form (Supporting Information, Figure S2A). Interestingly, in the complex with ACE2, this site also corresponded to a major sensor peak at the binding interface (Supporting Information, Figure S2B). Structural mapping of sensor profiles in the unbound and bound RBD forms illustrated these observations, pointing to role of E484 residue as a major receiver site of allosteric signaling in the RBD-ACE2 complex (Supporting Information, Figure S2C,and D). Hence, the PRS analysis of the RBD-ACE2 and S1-ACE2 complexes demonstrated that while E406, K417, N439 an N501 are aligned with dominant effector positions representing source and regulatory points of allosteric signaling, E484 corresponded to a major sensor/receiver site that may absorb signal information. Collectively, these sites may represent key nodes of the allosteric interaction network in the functional ACE2-bound complexes and determine the robustness and efficiency of signal transmission.

We also computed the PRS effector profiles for the SARS-RBD complex with REGN-COV2 antibody combination (Figure 10). The effector profile revealed some redistribution of peaks, featuring V401/E406, N439, K444/G446 and G496 positions as major effector centers (Figure 10A). At the same time, residues E484/F486 and N501 were aligned with the local sensor peaks (Figure 10B). These results could provide a feasible rationale for a critical role of K444 and F486 positions in escaping antibody combinations.

**Figure 10.**
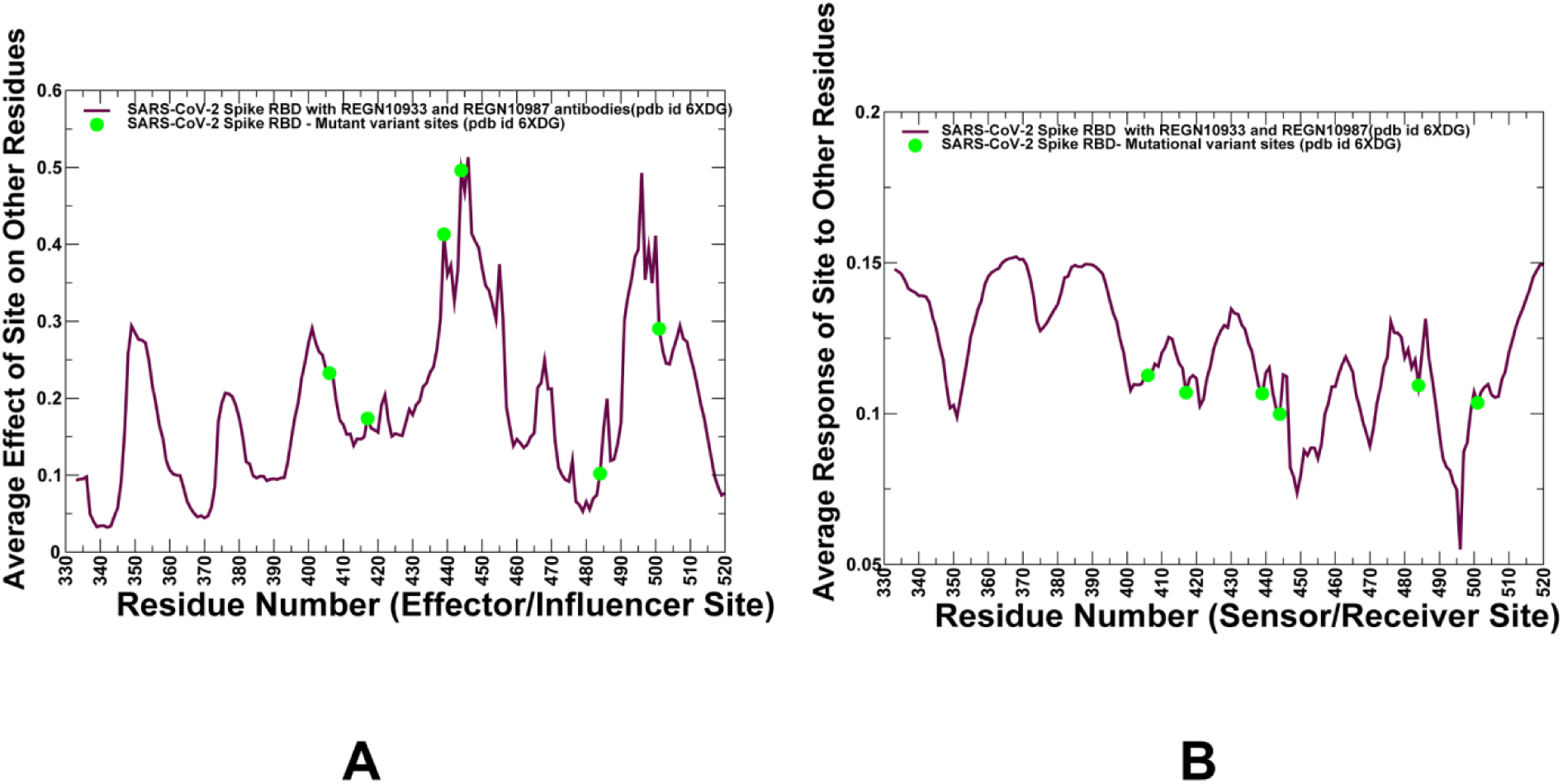
The PRS profiles for the SARS-CoV-2 S-RBD and S1 domain complexes with REGN-COV2 antibody cocktail. (A) The PRS effector profile for the SARS-CoV-2 S-RBD complex with REGN-COV2 antibodies (REGN10933 and REGN10987) (pdb id 6XDG). (B) The PRS sensor profile for the SARS-CoV-2 S-RBD complex with REGN-COV2 (pdb id 6XDG) The positions of functional residues E406, K417, N439, K444, E484, and N501 are indicated by filled green circles.

Indeed, K444 is a central epitope residue for REG10987 while F486 residue is fundamental for recognition of REG10933 antibody.^70^ Our findings also indicated that E406 and K444 are the dominant effector centers in the RBD complex with REGN-COV2 (Figure 10A) and may be functionally important not only for binding affinity but also for mediating signaling and long-range communications in the complex. In the context of perturbation-based PRS model, this implies that single mutations at these positions could affect collective movements and allosteric couplings between many residues in the system and potentially compromise functional activity of the REGN-COV2 cocktail. Interestingly, other positions targeted by antibody-escaping mutants E484 and F486 are the major sensor sites (Figure 10B). Based on these observations, we suggest that allosteric control of the RBD-REGN COV2 complex is provided through a cross-talk between major effector sites (E406, K444) and receiver sites (E484 and F486).

To summarize, perturbation-based modeling of the SARS-CoV-2 RBD complexes suggested that functional residues targeted by global circulating variants and antibody-escaping mutants could form a network of structurally adaptable allosteric hotspots that collectively coordinate allosteric interactions in the system. These results bear some significance and support the latest illuminating study suggesting a model functional plasticity and evolutionary adaptation of allosteric regulation.^156^ This function-centric model of allostery revealed a remarkable functional plasticity of allosteric switches allowing modulate and restore regulatory activity through mutational combinations or ligand interactions. Our results similarly suggested that functional plasticity and cross-talk of allosteric control points in the SARS-CoV-2 RBD region can allow for differential modulation of recognition and long-range communication with ACE2 and antibodies.

### Network Modeling Reveals that Sites Targeted by Circulating Mutations are Mediating Anchors of the Intermolecular Communities with ACE2 and REGN-COV2 Antibodies

Mechanistic network-based models allow for a quantitative analysis of allosteric molecular events in which conformational landscapes of protein systems can be remodeled by various perturbations such as mutations, ligand binding, or interactions with other proteins. The residue interaction networks in the SARS-CoV-2 RBD complexes with ACE2 and REGN-COV2 antibodies were built using a graph-based representation of protein structures in which residue nodes are interconnected through both dynamic^134^ and coevolutionary correlations.^135^ Using community decomposition, the residue interaction networks were further divided into local stable interaction modules in which residues are densely interconnected and highly correlated during simulations, while different communities are connected through long-range couplings.

Using this network-centric description of residue interactions, we compared the organization of stable local communities in the SARS-CoV-2 RBD complexes with ACE2 (Figure 11) and REGN-COV2 cocktail (Figure 12). In the RBD-ACE2 and S1-ACE2 complexes we found a deeply interconnected community organization (Figure 11) where stable modules in the RBD core are tightly linked with the interfacial clusters. A number of stable intramolecular communities in the RBD core contribute to stability of the RBD regions. Some of these communities are formed by hydrophobic core residues including F342-V511-F374-W436-F347-R509, W353-F400-Y423 as well as E406-Q409-I418 and N439-443-P499 centered around functional residues E406 and N439 (Figure 11). Of particular interest an importance was a more detailed comparative analysis of the inter-molecular communities. In the RBD-ACE2 and S1-ACE2 complexes, these modules are integrated around key anchor residues K417, F456, Y489, N501, and Y505. The largest and most stable community in which each node is strongly linked with each other is centered on N501 (K353-D38-Y41-Q498-N501-Y505) and engaged ACE2 hotspots K353, D38, and Y41 (Figure 11A). This interfacial module anchored by N501 allows for persistent interactions by N501, Y505 and Q498 RBD residues. Moreover, this community may be instrumental for signal transmission between RBD and ACE2 molecules, highlighting the important role of N501 position. By introducing N501Y mutation, we rebuilt the residue interaction network and performed community decomposition which revealed preservation of this major community. This observation supported our energetic analysis indicating structural plasticity and stability of the key intermolecular communities in the RBD-ACE2 complex.

**Figure 11.**
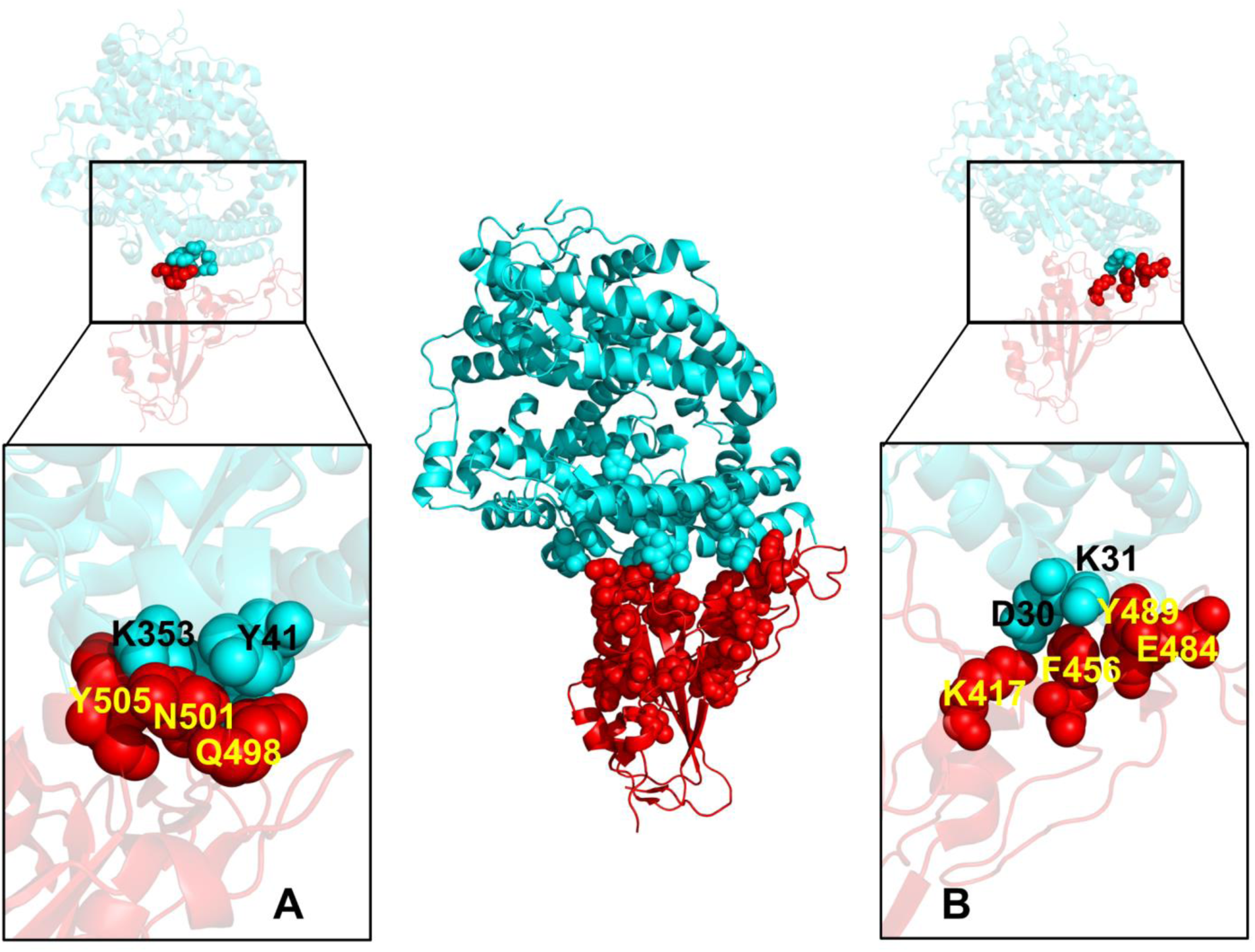
Structural maps of the intra- and intermolecular local communities formed in the SARS-CoV-2 S-RBD complex with ACE2. (Central panel) Molecular topography of major communities in the SARS-CoV-2 S-RBD complex with ACE2. SARS-CoV-2 RBD is shown in red ribbons and ACE2 in cyan ribbons. The RBD residues involved in local communities are shown in red spheres and ACE2 residues involved in the inter-molecular communities only are shown in cyan spheres (Left panel A) A close-up of the major intermolecular community that is anchored by N501 and includes Y505, Q498 residues of RBD and K353 and Y41 hotspot residues of ACE2 are highlighted in red and cyan spheres respectively and annotated. (Right panel B). A close-up of another dominant interfacial community anchored by F456 is shown. The community includes K417, F456, Y489 and E484 residues of RBD and K31/D30 hotspot residues of ACE2. The community residues are shown in spheres and annotated. Notably, these two major intermolecular communities that mediate communications and stability of the interface include several key functional sites K417, F456, E484, and N501 targeted by mutational variants and antibody-escaping mutations.

**Figure 12.**
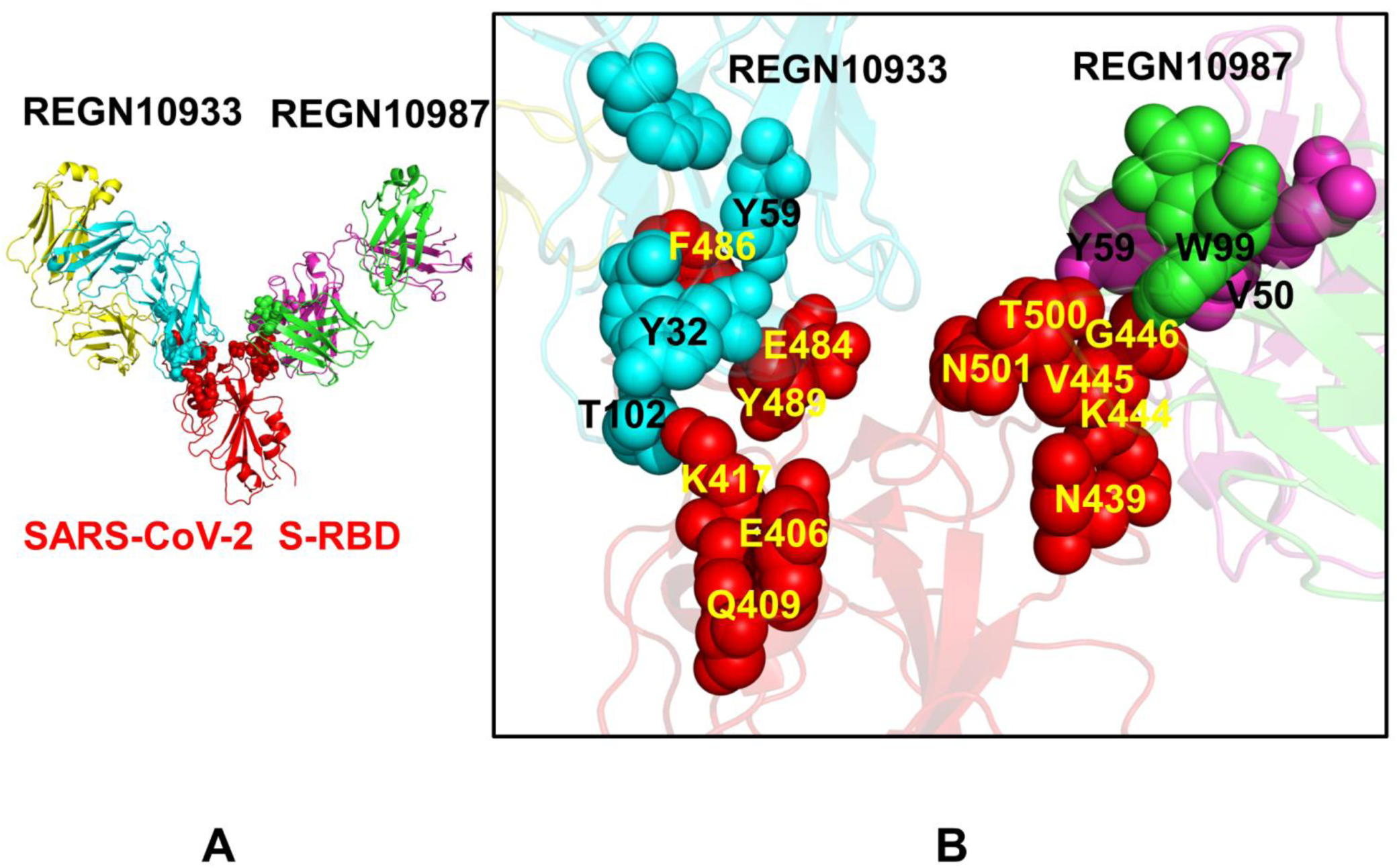
Structural maps of local communities in the SARS-CoV-2 S-RBD complex with REGN-COV2 cocktail of antibodies. (A) A general community overview of the SARS-CoV-2 S-RBD complex with REGN-COV2. The RBD is in red ribbons and interfacial communities are shown in red spheres. (B) A close-up of the intermolecular communities formed by the S-RBD. The communities at the interface with REGN10933 include F486(RBD)-E484(RBD)-Y489(RBD)-R100(REGN10933)-Y50(REGN10933)-Y59(REGN10933)-L94(REGN10933) and K417(RBD)-E406(RBD)-Q409(RBD)-Y32(REGN10933)-T102(REGN10933). The interfacial community with another antibody in the complex is W99(REGN10987)-W47(REGN10987)-G446(RBD)-V445(RBD)-K444(RBD)-Y59(REGN10987) that is anchored by K444, V445 and G446 residues connected with Y59 and W99 of REGN10987.

Another group of interfacial communities is anchored by K417 and F456 positions that couple modules Y489-F456-K31 and D30-K417-F456 (Figure 11B). Interestingly, these intermolecular communities are directly coupled through K417 with the intramolecular module I402-I418-E406-Y495-Q409 centered on the E406 residue. Hence, the community organization revealed strong interconnectivity between key functional sites E406, K417, F456, and N501 that integrate the residue interaction network and enable allosteric couplings between RBD and ACE2 molecules. The important revelation of this analysis is that only a fraction of the RBD residues anchor the intermolecular community organization and mediate long-range interactions in the RBD-ACE2 complex. Furthermore, the binding free energy hotspots are not necessarily involved in community-mediating function. Instead, a group of structurally plastic allosteric centers such as E406, K417, F456, and N501 play key roles in integrating local communities into a robust and adaptable global network that can mediate signal transmission and communication between SARS-CoV-2 RBD and ACE2.

The REGN-COV2 antibody binding induced a partial and yet significant reorganization of the interface communities (Figure 12A). We found that key RBD residues F486, Y489 and K417 anchor major intermolecular communities with REGN10933 including R100-Y50-F486-W47-L94, Y32-T102-K417 and Y33-T52-Y489 (Figure 12B). Interestingly, K417 and another site of escape mutations E406 are inter-connected in the local RBD community E406-Q409-K417-I418. E484 is involved in formation of the inter-molecular contacts with T52, Y53, T57, Y59 residues of the heavy chain of REGN10933 and could bridge several interfacial communities anchored by F486 and Y489 residues. Among major interfacial communities formed by the RBD with REGN10987 is W99-W47-V445-K444-Y59 module in which V445 and K444 play a key role in mediating intermolecular communication. Another notable community is anchored by T500 which engages N439, P499, K444 residues of the RBD and W99 of the heavy chain of REGN10987 (Figure 12B). These findings are consistent with the experimental deep mutational screening showing that K444 is a critical RBD residue for REGN10987, while F486 and Y489 sites are essential in inducing activity of REGN10933. According to these experiments, mutations F486I and Y489H escape REGN10933 and E484A/F486I combinations evade REGN10933. Interestingly, it was also established that mutations at F486 escaped neutralization only by REGN10933, whereas mutations of N439 and K444 escaped neutralization only by REGN10987.

E406 site is at the center of the largest intramolecular community in the SARS-CoV-2 RBD (R403-Y453-V350-I418-N422-Y423-Y495-F497-E406-Q409) that connects the inter-molecular interfaces with the RBD core (Supporting Information, Figure S3). This site could anchor large communities in the core with the interfacial regions facing both antibodies. Based on these observations, we suggested that unique escape mutations in E406 position may be largely determined by its allosteric mediating role in interconnecting functional regions of the RBD. Given the fact that E406 is only involved in several contacts with T28 of REGN10933, the strong mutational escape effect may be mainly driven by long-range allosteric effects and attributed to the strategic position of this residue in the global network. Noticeably, E406 is closely connected with Y453 residue where another REGN10933 specific escape-mutant Y453F was detected.^70^ Another key member of this community is F497 that effectively bridges local intermolecular modules interacting with REGN10933 and REGN10987 antibodies (Supporting Information, Figure S3). Recent studies indicated that SARS-CoV-2 neighbor residues G496 and F497 are critical for the RBD–ACE2 interaction and F497 may play important role for enhancing the RBD–ACE2 interaction for SARS-Cov-2 RBD. Our network analysis quantifies the structural hypothesis offered in the experimental study according to which E406 escape mutation may affect recognition by REGN10987 through cascading effect onto adjacent structural elements across the RBD and propagating changes through aromatic residues Y453, Y495, and F497.^70^ Importantly, network analysis revealed that these hydrophobic residues belong to the tightly packed stable community anchored by the E406 residue. Owing to the modular interconnectivity where each of these residues is connected with every community neighbor, it is likely that E406W mutation may simultaneously perturb multiple contacts and alter couplings between these residues, thus adversely affecting the fidelity of allosteric communication with REGN10987. Hence, although this residue makes no persistent contacts with either of the interacting antibodies, its position in the largest community anchored by E406 could be important for integrity of the network organization in the complex with antibodies. The network analysis showed that not only these residues are involved in favorable contacts with these antibodies but they also define key regulatory nodes that mediate stability and connectivity of the inter-molecular communities and may be responsible for control of signal transmission between SARS-CoV-2 RBD and REGN10933/REGN10987 antibodies. In agreement with functional data, the network community analysis singled out N439, K444, F486 and Y489 sites as allosteric network hubs where mutations would result in weakening of the entire interface and compromise the efficiency of allosteric interactions.

## Conclusions

This integrative computational investigation combined molecular simulations and functional dynamics analysis with mutational energetic profiling of the SARS-CoV-2 S protein binding and network-based community modeling to delineate specific allosteric signatures of functionally important residues that are subjected to novel circulating variants. A comparative perturbation-based modeling of the SARS-CoV-2 S complexes with ACE2 and REGN-COV2 antibody combination revealed several important trends and characterized the unique allosteric-centric signatures of functional RBD residues. First, we found that the RBM residues targeted by novel mutational variants may be fairly structurally adaptable and display a range of flexibility – from more dynamic positions at N439 and E486 to more constrained K417 and N501 residues. Functional dynamics analysis of global motions and identification of major hinge centers in the SARS-CoV-2 S complexes demonstrated that these sites play a central role in regulation of collective movements and long-range couplings, explaining why mutations in these positions could escape from antibody binding while maintaining and enhancing binding with the host receptor. Through comprehensive mutational scanning and sensitivity analysis of the SARS-CoV-2 RBD residues in the studied complexes, we accurately reproduced the binding affinity changes for N501Y, E484K and K417N mutations. Moreover, this analysis strongly indicated that these residues correspond to important interacting centers with a significant degree of structural and energetic plasticity. As a result, we proposed a model according to which circulating variants and antibody-escaping mutations tend to target residues with sufficient plasticity and adaptability to preserve a sufficient spike activity while having a more detrimental effect on antibody recognition. Our results also suggested that these functionally important for spike activity residues correspond to structurally plastic and energetically adaptable allosteric centers involved in modulation of global motions and allosteric interactions in the complexes. Using PRS analysis and community modeling of the SARS-CoV-2 S complexes, we characterized signatures of functional residues implicated in novel variants and demonstrated that these sites form a network of energetically adaptable regulatory centers modulating long-range communication of the SARS-CoV-2 S-RBD regions with ACE2 and antibodies. The results demonstrate that SARS-CoV-2 S protein may function as a versatile and functionally adaptable allosteric machine that exploits plasticity of allosteric regulatory centers to generate escape mutants that fine-tune response to antibody binding without compromising activity of the spike protein. This study provides a novel insight into allosteric regulatory mechanisms of SARS-CoV-2 S proteins, showing that antibodies can modulate allosteric interactions and signaling of spike proteins, providing a plausible strategy for therapeutic intervention by targeting specific hotspots of allosteric interactions in the SARS-CoV-2 proteins.

## ABBREVIATIONS

SARS: Severe Acute Respiratory Syndrome
RBD: Receptor Binding Domain
ACE2: Angiotensin-Converting Enzyme 2 (ACE2)
NTD: N-terminal domain
RBD: receptor-binding domain
CTD1: C-terminal domain 1
CTD2: C-terminal domain 2.

## SUPPORTING INFORMATION

Figure S1 presents the computations of the solvent accessible surface area (SASA) and RSA ratio for the SARS-CoV-2 S complexes with ACE2 and REGN-COV2 antibody cocktail. Figure S2 reports the PRS sensor profiles for the unbound and bound forms of the SARS-CoV-2 RBD with ACE2 host receptor. Figure S3 describes the structural maps of local communities in the SARS-CoV-2 S-RBD complex with REGN-COV2 cocktail of antibodies REGN10933 and REGN10987. Supporting information contains Tables S1, S2 and S3 that list the intermolecular contacts formed by the S-RBD residues in the SARS-CoV-2 S-RBD complex with ACE2 (pdb id 6M0J) and S1 domain complexes with the host receptor (pdb id 7A91 and 7A92) respectively. Table S4 lists the intermolecular contacts formed by the S-RBD residues in the SARS-CoV-2 S-RBD complex with REGN-COV2 antibody cocktail (pdb id 6XDG).

This material is available free of charge via the Internet at http://pubs.acs.org.

## AUTHOR INFORMATION

The authors declare no competing financial interest.

## Acknowledgment

This work was partly supported by institutional funding from Chapman University. The author acknowledges support by the Kay Family Foundation Grant A20-0032.

---

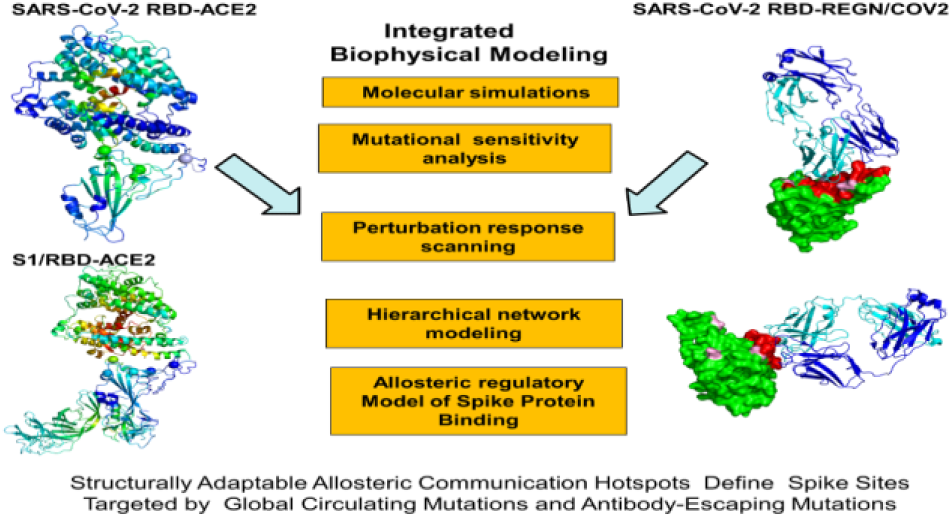
For Table of Contents Use Only.

## References

(1) Li, Q.; Guan, X.; Wu, P.; Wang, X.; Zhou, L.; Tong, Y.; Ren, R.; Leung, K. S. M.; Lau, E. H. Y.; Wong, J. Y.; Xing, X.; Xiang, N.; Wu, Y.; Li, C.; Chen, Q.; Li, D.; Liu, T.; Zhao, J.; Liu, M.; Tu, W.; Chen, C.; Jin, L.; Yang, R.; Wang, Q.; Zhou, S.; Wang, R.; Liu, H.; Luo, Y.; Liu, Y.; Shao, G.; Li, H.; Tao, Z.; Yang, Y.; Deng, Z.; Liu, B.; Ma, Z.; Zhang, Y.; Shi, G.; Lam, T. T. Y.; Wu, J. T.; Gao, G. F.; Cowling, B. J.; Yang, B.; Leung, G. M.; Feng, Z. Early transmission dynamics in Wuhan, China, of novel coronavirus-infected pneumonia. N. Engl. J. Med. 2020, 382, 1199–1207.

(2) Wang, C.; Horby, P. W.; Hayden, F. G.; Gao, G. F. A novel coronavirus outbreak of global health concern. Lancet 2020, 395, 470–473.

(3) Yi, Y.; Lagniton, P. N. P.; Ye, S.; Li, E.; Xu, R. H. COVID-19: what has been learned and to be learned about the novel coronavirus disease. Int. J. Biol. Sci. 2020, 16, 1753–1766.

(4) Wu, A.; Peng, Y.; Huang, B.; Ding, X.; Wang, X.; Niu, P.; Meng, J.; Zhu, Z.; Zhang, Z.; Wang, J.; Sheng, J.; Quan, L.; Xia, Z.; Tan, W.; Cheng, G.; Jiang, T. Genome composition and divergence of the novel coronavirus (2019-nCoV) originating in China. Cell Host Microbe 2020, 27, 325–328.

(5) Tai, W.; He, L.; Zhang, X.; Pu, J.; Voronin, D.; Jiang, S.; Zhou, Y.; Du, L. Characterization of the receptor-binding domain (RBD) of 2019 novel coronavirus: implication for development of RBD protein as a viral attachment inhibitor and vaccine. Cell. Mol. Immunol. 2020, 17, 613–620.

(6) Hoffmann, M.; Kleine-Weber, H.; Schroeder, S.; Krüger, N.; Herrler, T.; Erichsen, S.; Schiergens, T. S.; Herrler, G.; Wu, N. H.; Nitsche, A.; Müller, M. A.; Drosten, C.; Pöhlmann, S. SARS-CoV-2 cell entry depends on ACE2 and TMPRSS2 and is blocked by a clinically proven protease inhibitor. Cell 2020, 181, 271–280.e8.

(7) Lu, R.; Zhao, X.; Li, J.; Niu, P.; Yang, B.; Wu, H.; Wang, W.; Song, H.; Huang, B.; Zhu, N.; Bi, Y.; Ma, X.; Zhan, F.; Wang, L.; Hu, T.; Zhou, H.; Hu, Z.; Zhou, W.; Zhao, L.; Chen, J.; Meng, Y.; Wang, J.; Lin, Y.; Yuan, J.; Xie, Z.; Ma, J.; Liu, W. J.; Wang, D.; Xu, W.; Holmes, E. C.; Gao, G. F.; Wu, G.; Chen, W.; Shi, W.; Tan, W. Genomic characterisation and epidemiology of 2019 novel coronavirus: implications for virus origins and receptor binding. Lancet 2020, 395, 565–574.

(8) Duan, L.; Zheng, Q.; Zhang, H.; Niu, Y.; Lou, Y.; Wang, H. The SARS-CoV-2 spike glycoprotein biosynthesis, structure, function, and antigenicity: Implications for the design of spike-based vaccine immunogens. Front. Immunol. 2020, 11, 576622.

(9) Wang, Q.; Zhang, Y.; Wu, L.; Niu, S.; Song, C.; Zhang, Z.; Lu, G.; Qiao, C.; Hu, Y.; Yuen, K. Y.; Zhou, H.; Yan, J.; Qi, J. Structural and functional basis of SARS-CoV-2 entry by using human ACE2. Cell 2020, 181, 894–904.e9.

(10) Wan, Y.; Shang, J.; Graham, R.; Baric, R. S.; Li, F. Receptor recognition by the novel coronavirus from Wuhan: An analysis based on decade-long structural studies of SARS coronavirus. J. Virol. 2020, 94, e00127–20.

(11) Shang, J.; Wan, Y.; Luo, C.; Ye, G.; Geng, Q.; Auerbach, A.; Li, F. Cell entry mechanisms of SARS-CoV-2. Proc. Natl. Acad. Sci. U. S. A. 2020, 117, 11727–11734.

(12) Wang, Q.; Zhang, Y.; Wu, L.; Niu, S.; Song, C.; Zhang, Z.; Lu, G.; Qiao, C.; Hu, Y.; Yuen, K. Y.; Zhou, H.; Yan, J.; Qi, J. Structural and Functional Basis of SARS-CoV-2 Entry by Using Human ACE2. Cell 2020, 181, 894–904.e9.

(13) Wan, Y.; Shang, J.; Graham, R.; Baric, R. S.; Li, F. Receptor Recognition by the Novel Coronavirus from Wuhan: an Analysis Based on Decade-Long Structural Studies of SARS Coronavirus. J. Virol. 2020, 94, e00127–20.

(14) Li, F.; Li, W.; Farzan, M.; Harrison, S. C. Structure of SARS coronavirus spike receptor-binding domain complexed with receptor. Science 2005, 309, 1864–1868.

(15) Lan, J.; Ge, J.; Yu, J.; Shan, S.; Zhou, H.; Fan, S.; Zhang, Q.; Shi, X.; Wang, Q.; Zhang, L.; Wang, X. Structure of the SARS-CoV-2 spike receptor-binding domain bound to the ACE2 receptor. Nature 2020, 581, 215–220.

(16) Shang, J.; Ye, G.; Shi, K.; Wan, Y.; Luo, C.; Aihara, H.; Geng, Q.; Auerbach, A.; Li, F. Structural basis of receptor recognition by SARS-CoV-2. Nature 2020, 581, 221–224.

(17) Kirchdoerfer, R. N.; Cottrell, C. A.; Wang, N.; Pallesen, J.; Yassine, H. M.; Turner, H. L.; Corbett, K. S.; Graham, B. S.; McLellan, J. S.; Ward, A. B. Pre-fusion structure of a human coronavirus spike protein. Nature 2016, 531, 118–121.

(18) Walls, A. C.; Tortorici, M. A.; Bosch, B. J.; Frenz, B.; Rottier, P. J. M.; DiMaio, F.; Rey, F. A.; Veesler, D. Cryo-electron microscopy structure of a coronavirus spike glycoprotein trimer. Nature 2016, 531, 114–117.

(19) Gui, M.; Song, W.; Zhou, H.; Xu, J.; Chen, S.; Xiang, Y.; Wang, X. Cryo-electron microscopy structures of the SARS-CoV spike glycoprotein reveal a prerequisite conformational state for receptor binding. Cell Res. 2017, 27, 119–129.

(20) Walls, A. C.; Xiong, X.; Park, Y. J.; Tortorici, M. A.; Snijder, J.; Quispe, J.; Cameroni, E.; Gopal, R.; Dai, M.; Lanzavecchia, A.; Zambon, M.; Rey, F. A.; Corti, D.; Veesler, D. Unexpected receptor functional mimicry elucidates activation of coronavirus fusion. Cell 2019, 176, 1026–1039.e15.

(21) Yuan, Y.; Cao, D.; Zhang, Y.; Ma, J.; Qi, J.; Wang, Q.; Lu, G.; Wu, Y.; Yan, J.; Shi, Y.; Zhang, X.; Gao, G. F. Cryo-EM structures of MERS-CoV and SARS-CoV spike glycoproteins reveal the dynamic receptor binding domains. Nat. Commun. 2017, 8, 15092.

(22) Song, W.; Gui, M.; Wang, X.; Xiang, Y. Cryo-EM structure of the SARS coronavirus spike glycoprotein in complex with its host cell receptor ACE2. PLoS Pathog. 2018, 14, e1007236.

(23) Kirchdoerfer, R. N.; Wang, N.; Pallesen, J.; Wrapp, D.; Turner, H. L.; Cottrell, C. A.; Corbett, K. S.; Graham, B. S.; McLellan, J. S.; Ward, A. B. Stabilized coronavirus spikes are resistant to conformational changes induced by receptor recognition or proteolysis. Sci. Rep. 2018, 8, 15701.

(24) Walls, A. C.; Park, Y. J.; Tortorici, M. A.; Wall, A.; McGuire, A. T.; Veesler, D. Structure, Function, and Antigenicity of the SARS-CoV-2 Spike Glycoprotein. Cell 2020, 181, 281–292.e6.

(25) Wrapp, D.; Wang, N.; Corbett, K. S.; Goldsmith, J. A.; Hsieh, C. L.; Abiona, O.; Graham, B. S.; McLellan, J. S. Cryo-EM structure of the 2019-nCoV spike in the prefusion conformation. Science 2020, 367, 1260–1263.

(26) Cai, Y.; Zhang, J.; Xiao, T.; Peng, H.; Sterling, S. M.; Walsh, R. M., Jr.; Rawson, S.; Rits-Volloch, S.; Chen, B. Distinct conformational states of SARS-CoV-2 spike protein. Science 2020, 369, 1586–1592.

(27) Hsieh, C. L.; Goldsmith, J. A.; Schaub, J. M.; DiVenere, A. M.; Kuo, H. C.; Javanmardi, K.; Le, K. C.; Wrapp, D.; Lee, A. G.; Liu, Y.; Chou, C. W.; Byrne, P. O.; Hjorth, C. K.; Johnson, N. V.; Ludes-Meyers, J.; Nguyen, A. W.; Park, J.; Wang, N.; Amengor, D.; Lavinder, J. J.; Ippolito, G. C.; Maynard, J. A.; Finkelstein, I. J.; McLellan, J. S. Structure-based design of prefusion-stabilized SARS-CoV-2 spikes. Science 2020, 369, 1501–1505.

(28) Henderson, R.; Edwards, R. J.; Mansouri, K.; Janowska, K.; Stalls, V.; Gobeil, S. M. C.; Kopp, M.; Li, D.; Parks, R.; Hsu, A. L.; Borgnia, M. J.; Haynes, B. F.; Acharya, P. Controlling the SARS-CoV-2 spike glycoprotein conformation. Nat. Struct. Mol. Biol. 2020, 27, 925–933.

(29) McCallum, M.; Walls, A. C.; Bowen, J. E.; Corti, D.; Veesler, D. Structure-guided covalent stabilization of coronavirus spike glycoprotein trimers in the closed conformation. Nat. Struct. Mol. Biol. 2020, 27, 942–949.

(30) Xiong, X.; Qu, K.; Ciazynska, K. A.; Hosmillo, M.; Carter, A. P.; Ebrahimi, S.; Ke, Z.; Scheres, S. H. W.; Bergamaschi, L.; Grice, G. L.; Zhang, Y.; Nathan, J. A.; Baker, S.; James, L. C.; Baxendale, H. E.; Goodfellow, I.; Doffinger, R.; Briggs, J. A. G. A thermostable, closed SARS-CoV-2 spike protein trimer. Nat. Struct. Mol. Biol. 2020, 27, 934–941.

(31) Yurkovetskiy, L.; Wang, X.; Pascal, K. E.; Tomkins-Tinch, C.; Nyalile, T. P.; Wang, Y.; Baum, A.; Diehl, W. E.; Dauphin, A.; Carbone, C.; Veinotte, K.; Egri, S. B.; Schaffner, S. F.; Lemieux, J. E.; Munro, J. B.; Rafique, A.; Barve, A.; Sabeti, P. C.; Kyratsous, C. A.; Dudkina, N. V.; Shen, K.; Luban, J. Structural and functional analysis of the D614G SARS-CoV-2 spike protein variant. Cell 2020, 183, 739–751.e8.

(32) Turoňová, B.; Sikora, M.; Schürmann, C.; Hagen, W. J. H.; Welsch, S.; Blanc, F. E. C.; von Bülow, S.; Gecht, M.; Bagola, K.; Hörner, C.; van Zandbergen, G.; Landry, J.; de Azevedo, N. T. D.; Mosalaganti, S.; Schwarz, A.; Covino, R.; Mühlebach, M. D.; Hummer, G.; Krijnse Locker, J.; Beck, M. In situ structural analysis of SARS-CoV-2 spike reveals flexibility mediated by three hinges. Science 2020, 370, 203–208.

(33) Xu, C.; Wang, Y.; Liu, C.; Zhang, C.; Han, W.; Hong, X.; Wang, Y.; Hong, Q.; Wang, S.; Zhao, Q.; Wang, Y.; Yang, Y.; Chen, K.; Zheng, W.; Kong, L.; Wang, F.; Zuo, Q.; Huang, Z., Cong, Y. Conformational dynamics of SARS-CoV-2 trimeric spike glycoprotein in complex with receptor ACE2 revealed by cryo-EM. Sci Adv 2021, 7, eabe5575.

(34) Benton, D. J.; Wrobel, A. G.; Xu, P.; Roustan, C.; Martin, S. R.; Rosenthal, P. B.; Skehel, J. J.; Gamblin, S. J. Receptor binding and priming of the spike protein of SARS-CoV-2 for membrane fusion. Nature 2020, 588, 327–330.

(35) Starr, T. N.; Greaney, A. J.; Hilton, S. K.; Ellis, D.; Crawford, K. H. D.; Dingens, A. S.; Navarro, M. J.; Bowen, J. E.; Tortorici, M. A.; Walls, A. C.; King, N. P.; Veesler, D.; Bloom, J. D. Deep Mutational Scanning of SARS-CoV-2 Receptor Binding Domain Reveals Constraints on Folding and ACE2 Binding. Cell 2020, 182, 1295–1310.e20.

(36) Chan, K. K.; Dorosky, D.; Sharma, P.; Abbasi, S. A.; Dye, J. M.; Kranz, D. M.; Herbert, A. S.; Procko, E. Engineering human ACE2 to optimize binding to the spike protein of SARS coronavirus 2. Science 2020, 369, 1261–1265.

(37) Yi, C.; Sun, X.; Ye, J.; Ding, L.; Liu, M.; Yang, Z.; Lu, X.; Zhang, Y.; Ma, L.; Gu, W.; Qu, A.; Xu, J.; Shi, Z.; Ling, Z.; Sun, B. Key residues of the receptor binding motif in the spike protein of SARS-CoV-2 that interact with ACE2 and neutralizing antibodies. Cell Mol Immunol 2020, 17, 621–630.

(38) Wrapp, D.; De Vlieger, D.; Corbett, K. S.; Torres, G. M.; Wang, N.; Van Breedam, W.; Roose, K.; van Schie, L.; Hoffmann, M.; Pöhlmann, S.; Graham, B. S.; Callewaert, N.; Schepens, B.; Saelens, X.; McLellan, J. S. Structural Basis for Potent Neutralization of Betacoronaviruses by Single-Domain Camelid Antibodies. Cell 2020, 181, 1004–1015.e15.

(39) Yu, F.; Xiang, R.; Deng, X.; Wang, L.; Yu, Z.; Tian, S.; Liang, R.; Li, Y.; Ying, T.; Jiang, S. Receptor-binding domain-specific human neutralizing monoclonal antibodies against SARS-CoV and SARS-CoV-2. Signal Transduct. Target. Ther. 2020, 5, 212.

(40) Wang, C.; Li, W.; Drabek, D.; Okba, N. M. A.; van Haperen, R.; Osterhaus, A.; van Kuppeveld, F. J. M.; Haagmans, B. L.; Grosveld, F.; Bosch, B. J. A human monoclonal antibody blocking SARS-CoV-2 infection. Nat. Commun. 2020, 11, 2251.

(41) Hwang, W. C.; Lin, Y.; Santelli, E.; Sui, J.; Jaroszewski, L.; Stec, B.; Farzan, M.; Marasco, W. A.; Liddington, R. C. Structural basis of neutralization by a human anti-severe acute respiratory syndrome spike protein antibody, 80R. J. Biol. Chem. 2006, 281, 34610–34616.

(42) Yuan, M.; Wu, N. C.; Zhu, X.; Lee, C. D.; So, R. T. Y.; Lv, H.; Mok, C. K. P.; Wilson, I. A. A highly conserved cryptic epitope in the receptor binding domains of SARS-CoV-2 and SARS-CoV. Science 2020, 368, 630–633.

(43) Huo, J.; Zhao, Y.; Ren, J.; Zhou, D.; Duyvesteyn, H. M. E.; Ginn, H. M.; Carrique, L.; Malinauskas, T.; Ruza, R. R.; Shah, P. N. M.; Tan, T. K.; Rijal, P.; Coombes, N.; Bewley, K. R.; Tree, J. A.; Radecke, J.; Paterson, N. G.; Supasa, P.; Mongkolsapaya, J.; Screaton, G. R.; Carroll, M.; Townsend, A.; Fry, E. E.; Owens, R. J.; Stuart, D. I. Neutralization of SARS-CoV-2 by destruction of the prefusion spike. Cell Host Microbe 2020, 28, 445–454.e6.

(44) Brouwer, P. J. M.; Caniels, T. G.; van der Straten, K.; Snitselaar, J. L.; Aldon, Y.; Bangaru, S.; Torres, J. L.; Okba, N. M. A.; Claireaux, M.; Kerster, G.; Bentlage, A. E. H.; van Haaren, M. M.; Guerra, D.; Burger, J. A.; Schermer, E. E.; Verheul, K. D.; van der Velde, N.; van der Kooi, A.; van Schooten, J.; van Breemen, M. J.; Bijl, T. P. L.; Sliepen, K.; Aartse, A.; Derking, R.; Bontjer, I.; Kootstra, N. A.; Wiersinga, W. J.; Vidarsson, G.; Haagmans, B. L.; Ward, A. B.; de Bree, G. J.; Sanders, R. W.; van Gils, M. J. Potent neutralizing antibodies from COVID-19 patients define multiple targets of vulnerability. Science 2020, 369, 643–650.

(45) Lv, Z.; Deng, Y. Q.; Ye, Q.; Cao, L.; Sun, C. Y.; Fan, C.; Huang, W.; Sun, S.; Sun, Y.; Zhu, L.; Chen, Q.; Wang, N.; Nie, J.; Cui, Z.; Zhu, D.; Shaw, N.; Li, X. F.; Li, Q.; Xie, L.; Wang, Y.; Rao, Z.; Qin, C. F.; Wang, X. Structural basis for neutralization of SARS-CoV-2 and SARS-CoV by a potent therapeutic antibody. Science 2020, 369, 1505–1509.

(46) Pinto, D.; Park, Y. J.; Beltramello, M.; Walls, A. C.; Tortorici, M. A.; Bianchi, S.; Jaconi, S.; Culap, K.; Zatta, F.; De Marco, A.; Peter, A.; Guarino, B.; Spreafico, R.; Cameroni, E.; Case, J. B.; Chen, R. E.; Havenar-Daughton, C.; Snell, G.; Telenti, A.; Virgin, H. W.; Lanzavecchia, A.; Diamond, M. S.; Fink, K.; Veesler, D.; Corti, D. Cross-neutralization of SARS-CoV-2 by a human monoclonal SARS-CoV antibody. Nature 2020, 583, 290–295.

(47) Tortorici, M. A.; Beltramello, M.; Lempp, F. A.; Pinto, D.; Dang, H. V.; Rosen, L. E.; McCallum, M.; Bowen, J.; Minola, A.; Jaconi, S.; Zatta, F.; De Marco, A.; Guarino, B.; Bianchi, S.; Lauron, E. J.; Tucker, H.; Zhou, J.; Peter, A.; Havenar-Daughton, C.; Wojcechowskyj, J. A.; Case, J. B.; Chen, R. E.; Kaiser, H.; Montiel-Ruiz, M.; Meury, M.; Czudnochowski, N.; Spreafico, R.; Dillen, J.; Ng, C.; Sprugasci, N.; Culap, K.; Benigni, F.; Abdelnabi, R.; Foo, S. C.; Schmid, M. A.; Cameroni, E.; Riva, A.; Gabrieli, A.; Galli, M.; Pizzuto, M. S.; Neyts, J.; Diamond, M. S.; Virgin, H. W.; Snell, G.; Corti, D.; Fink, K.; Veesler, D. Ultrapotent human antibodies protect against SARS-CoV-2 challenge via multiple mechanisms. Science 2020, 370, 950–957.

(48) Chi, X.; Yan, R.; Zhang, J.; Zhang, G.; Zhang, Y.; Hao, M.; Zhang, Z.; Fan, P.; Dong, Y.; Yang, Y.; Chen, Z.; Guo, Y.; Li, Y.; Song, X.; Chen, Y.; Xia, L.; Fu, L.; Hou, L.; Xu, J.; Yu, C.; Li, J.; Zhou, Q.; Chen, W. A neutralizing human antibody binds to the N-terminal domain of the Spike protein of SARS-CoV-2. Science 2020, 369, 650–655.

(49) Schoof, M.; Faust, B.; Saunders, R. A.; Sangwan, S.; Rezelj, V.; Hoppe, N.; Boone, M.; Billesbølle, C. B.; Puchades, C.; Azumaya, C. M.; Kratochvil, H. T.; Zimanyi, M.; Deshpande, I.; Liang, J.; Dickinson, S.; Nguyen, H. C.; Chio, C. M.; Merz, G. E.; Thompson, M. C.; Diwanji, D.; Schaefer, K.; Anand, A. A.; Dobzinski, N.; Zha, B. S.; Simoneau, C. R.; Leon, K.; White, K. M.; Chio, U. S.; Gupta, M.; Jin, M.; Li, F.; Liu, Y.; Zhang, K.; Bulkley, D.; Sun, M.; Smith, A. M.; Rizo, A. N.; Moss, F.; Brilot, A. F.; Pourmal, S.; Trenker, R.; Pospiech, T.; Gupta, S.; Barsi-Rhyne, B.; Belyy, V.; Barile-Hill, A. W.; Nock, S.; Krogan, N. J.; Ralston, C. Y.; Swaney, D. L.; García-Sastre, A.; Ott, M.; Vignuzzi, M.; Walter, P.; Manglik, A. An ultrapotent synthetic nanobody neutralizes SARS-CoV-2 by stabilizing inactive Spike. Science 2020, 370, 1473–1479.

(50) Gavor, E.; Choong, Y. K.; Er, S. Y.; Sivaraman, H.; Sivaraman, J. Structural Basis of SARS-CoV-2 and SARS-CoV Antibody Interactions. Trends Immunol 2020, 41, 1006–1022.

(51) Barnes, C. O.; Jette, C. A.; Abernathy, M. E.; Dam, K. A.; Esswein, S. R.; Gristick, H. B.; Malyutin, A. G.; Sharaf, N. G.; Huey-Tubman, K. E.; Lee, Y. E.; Robbiani, D. F.; Nussenzweig, M. C.; West, A. P., Jr.; Bjorkman, P. J. SARS-CoV-2 neutralizing antibody structures inform therapeutic strategies. Nature 2020, 588, 682–687.

(52) Wu, Y.; Wang, F.; Shen, C.; Peng, W.; Li, D.; Zhao, C.; Li, Z.; Li, S.; Bi, Y.; Yang, Y.; Gong, Y.; Xiao, H.; Fan, Z.; Tan, S.; Wu, G.; Tan, W.; Lu, X.; Fan, C.; Wang, Q.; Liu, Y.; Zhang, C.; Qi, J.; Gao, G. F.; Gao, F.; Liu, L. A noncompeting pair of human neutralizing antibodies block COVID-19 virus binding to its receptor ACE2. Science 2020, 368, 1274–1278.

(53) Ju, B.; Zhang, Q.; Ge, J.; Wang, R.; Sun, J.; Ge, X.; Yu, J.; Shan, S.; Zhou, B.; Song, S.; Tang, X.; Lan, J.; Yuan, J.; Wang, H.; Zhao, J.; Zhang, S.; Wang, Y.; Shi, X.; Liu, L.; Wang, X.; Zhang, Z.; Zhang, L. Human neutralizing antibodies elicited by SARS-CoV-2 infection. Nature 2020, 584, 115–119.

(54) Shi, R.; Shan, C.; Duan, X.; Chen, Z.; Liu, P.; Song, J.; Song, T.; Bi, X.; Han, C.; Wu, L.; Gao, G.; Hu, X.; Zhang, Y.; Tong, Z.; Huang, W.; Liu, W. J.; Wu, G.; Zhang, B.; Wang, L.; Qi, J.; Feng, H.; Wang, F. S.; Wang, Q.; Gao, G. F.; Yuan, Z.; Yan, J. A human neutralizing antibody targets the receptor-binding site of SARS-CoV-2. Nature 2020, 584,120–124.

(55) Yuan, M.; Liu, H.; Wu, N. C.; Lee, C. D.; Zhu, X.; Zhao, F.; Huang, D.; Yu, W.; Hua, Y.; Tien, H.; Rogers, T. F.; Landais, E.; Sok, D.; Jardine, J. G.; Burton, D. R.; Wilson, I. A., Structural basis of a shared antibody response to SARS-CoV-2. Science 2020, 369, 1119–1123.

(56) Barnes, C. O.; West, A. P., Jr.; Huey-Tubman, K. E.; Hoffmann, M. A. G.; Sharaf, N. G.; Hoffman, P. R.; Koranda, N.; Gristick, H. B.; Gaebler, C.; Muecksch, F.; Lorenzi, J. C. C.; Finkin, S.; Hägglöf, T.; Hurley, A.; Millard, K. G.; Weisblum, Y.; Schmidt, F.; Hatziioannou, T.; Bieniasz, P. D.; Caskey, M.; Robbiani, D. F.; Nussenzweig, M. C.; Bjorkman, P. J. Structures of Human Antibodies Bound to SARS-CoV-2 Spike Reveal Common Epitopes and Recurrent Features of Antibodies. Cell 2020, 182, 828–842.e16.

(57) Cao, Y.; Su, B.; Guo, X.; Sun, W.; Deng, Y.; Bao, L.; Zhu, Q.; Zhang, X.; Zheng, Y.; Geng, C.; Chai, X.; He, R.; Li, X.; Lv, Q.; Zhu, H.; Deng, W.; Xu, Y.; Wang, Y.; Qiao, L.; Tan, Y.; Song, L.; Wang, G.; Du, X.; Gao, N.; Liu, J.; Xiao, J.; Su, X. D.; Du, Z.; Feng, Y.; Qin, C.; Jin, R.; Xie, X. S. Potent Neutralizing Antibodies against SARS-CoV-2 Identified by High-Throughput Single-Cell Sequencing of Convalescent Patients’ B Cells. Cell 2020, 182, 73–84.e16.

(58) Zhou, D.; Duyvesteyn, H. M. E.; Chen, C. P.; Huang, C. G.; Chen, T. H.; Shih, S. R.; Lin, Y. C.; Cheng, C. Y.; Cheng, S. H.; Huang, Y. C.; Lin, T. Y.; Ma, C.; Huo, J.; Carrique, L.; Malinauskas, T.; Ruza, R. R.; Shah, P. N. M.; Tan, T. K.; Rijal, P.; Donat, R. F.; Godwin, K.; Buttigieg, K. R.; Tree, J. A.; Radecke, J.; Paterson, N. G.; Supasa, P.; Mongkolsapaya, J.; Screaton, G. R.; Carroll, M. W.; Gilbert-Jaramillo, J.; Knight, M. L.; James, W.; Owens, R. J.; Naismith, J. H.; Townsend, A. R.; Fry, E. E.; Zhao, Y.; Ren, J.; Stuart, D. I.; Huang, K. A. Structural basis for the neutralization of SARS-CoV-2 by an antibody from a convalescent patient. Nat Struct Mol Biol 2020, 27, 950–958.

(59) Fiorentini, S.; Messali, S.; Zani, A.; Caccuri, F.; Giovanetti, M.; Ciccozzi, M.; Caruso, A., First detection of SARS-CoV-2 spike protein N501 mutation in Italy in August, 2020. Lancet Infect. Dis. 2021. S1473-3099(21)00007-4. doi: 10.1016/S1473-3099(21)00007-4.

(60) Davies, N.G.; Barnard, R.C.; Jarvis, C.I. Kucharski, A.J.; Munday, J.; Pearson, C.A.B.; Russell, T.W.; Tully, D.C.; Abbott, S.; Gimma, A.; Waites, W.; Wong, K.L.M.; van Zandvoort, K.; CMMID COVID-19 Working Group, Eggo, R.M.; Funk, S.; Jit, M.; Atkins, K.E.; Edmunds, W.J. Estimated transmissibility and severity of novel SARS-CoV-2 variant of concern 202012/01 in England. medRxiv 2021 doi.org/10.1101/2020.12.24.20248822.

(61) Davies, N. G.; Jarvis, C. I.; CMMID COVID-19 Working Group, Edmunds, W. J.; Jewell, N. P.; Diaz-Ordaz, K.; Keogh, R. H.. Increased hazard of death in community tested cases of SARS-CoV-2 Variant of Concern 202012/01. medRxiv 2021 doi.org/10.1101/2021.02.01.21250959.

(62) Tegally, H.; Wilkinson E.; Giovanetti M., et al. Emergence and rapid spread of a new severe acute respiratory syndrome-related coronavirus 2 (SARS-CoV-2) lineage with multiple spike mutations in South Africa. medRxiv 2020 doi: https://doi.org/10.1101/2020.12.21.20248640.

(63) Tegally, H.; Wilkinson, E.; Lessells, R. J.; Giandhari, J.; Pillay, S.; Msomi, N.; Mlisana, K.; Bhiman, J. N.; von Gottberg, A.; Walaza, S.; Fonseca, V.; Allam, M.; Ismail, A.; Glass, A. J.; Engelbrecht, S.; Van Zyl, G.; Preiser, W.; Williamson, C.; Petruccione, F.; Sigal, A.; Gazy, I.; Hardie, D.; Hsiao, N. Y.; Martin, D.; York, D.; Goedhals, D.; San, E. J.; Giovanetti, M.; Lourenço, J.; Alcantara, L. C. J.; de Oliveira, T., Sixteen novel lineages of SARS-CoV-2 in South Africa. Nat Med 2021. doi: 10.1038/s41591-021-01255-3.

(64) Voloch CM, da Silva F Jr R, de Almeida LGP, et al. Genomic characterization of a novel SARS-CoV-2 lineage from Rio de Janeiro, Brazil. 2020. medRxiv 2020 doi: https://doi.org/10.1101/2020.12.23.20248598.

(65) Sabino, E. C.; Buss, L. F.; Carvalho, M. P. S.; Prete, C. A., Jr.; Crispim, M. A. E.; Fraiji, N. A.; Pereira, R. H. M.; Parag, K. V.; da Silva Peixoto, P.; Kraemer, M. U. G.; Oikawa, M. K.; Salomon, T.; Cucunuba, Z. M.; Castro, M. C.; de Souza Santos, A. A.; Nascimento, V. H.; Pereira, H. S.; Ferguson, N. M.; Pybus, O. G.; Kucharski, A.; Busch, M. P.; Dye, C.; Faria, N. R., Resurgence of COVID-19 in Manaus, Brazil, despite high seroprevalence. Lancet 2021, 397, 452–455.

(66) Greaney, A. J.; Starr, T. N.; Gilchuk, P.; Zost, S. J.; Binshtein, E.; Loes, A. N.; Hilton, S. K.; Huddleston, J.; Eguia, R.; Crawford, K. H. D.; Dingens, A. S.; Nargi, R. S.; Sutton, R. E.; Suryadevara, N.; Rothlauf, P. W.; Liu, Z.; Whelan, S. P. J.; Carnahan, R. H.; Crowe, J. E., Jr.; Bloom, J. D. Complete Mapping of Mutations to the SARS-CoV-2 Spike Receptor-Binding Domain that Escape Antibody Recognition. Cell Host Microbe 2021, 29, 44–57.e9.

(67) Greaney, A.J.; Loes, A.N.; Crawford, K.H.D.; Starr, T.N.; Malone, K.D.; Chu, H.Y.; Bloom, J.D. Comprehensive mapping of mutations to the SARS-CoV-2 receptor-binding domain that affect recognition by polyclonal human serum antibodies Cell Host Microbe 2021, S1931-3128(21)00082-2. doi: 10.1016/j.chom.2021.02.003.

(68) Andreano, E., Piccini, G., Licastro, D., Casalino, L., Johnson, N.V., Paciello, I., Monego, S.D., Pantano, E., Manganaro, N., Manenti, A., et al. SARS-CoV-2 escape in vitro from a highly neutralizing COVID-19 convalescent plasma. bioRxiv 2020 doi: 10.1101/2020.12.28.424451.

(69) Liu, Z.; VanBlargan, L.A.; Bloyet, L.M.; Rothlauf, P.W.; Chen, R.E.; Stumpf, S.; Zhao, H.; Errico, J.M.; Theel, E.S.; Liebeskind, M.J.; Alford, B.; Buchser, W.J.; Ellebedy, A.H.; Fremont, D.H.; Diamond, M.S.; Whelan, S.P.J. Landscape analysis of escape variants identifies SARS-CoV-2 spike mutations that attenuate monoclonal and serum antibody neutralization. bioRxiv 2020 doi: 10.1101/2020.11.06.372037.

(70) Starr, T. N.; Greaney, A. J.; Addetia, A.; Hannon, W. W.; Choudhary, M. C.; Dingens, A. S.; Li, J. Z.; Bloom, J. D. Prospective mapping of viral mutations that escape antibodies used to treat COVID-19. Science 2021, 371,850–854.

(71) Weisblum, Y.; Schmidt, F.; Zhang, F.; DaSilva, J.; Poston, D.; Lorenzi, J. C.; Muecksch, F.; Rutkowska, M.; Hoffmann, H. H.; Michailidis, E.; Gaebler, C.; Agudelo, M.; Cho, A.; Wang, Z.; Gazumyan, A.; Cipolla, M.; Luchsinger, L.; Hillyer, C. D.; Caskey, M.; Robbiani, D. F.; Rice, C. M.; Nussenzweig, M. C.; Hatziioannou, T.; Bieniasz, P. D. Escape from neutralizing antibodies by SARS-CoV-2 spike protein variants. Elife 2020, 9, e61312. doi: 10.7554/eLife.61312.

(72) Baum, A.; Fulton, B. O.; Wloga, E.; Copin, R.; Pascal, K. E.; Russo, V.; Giordano, S.; Lanza, K.; Negron, N.; Ni, M.; Wei, Y.; Atwal, G. S.; Murphy, A. J.; Stahl, N.; Yancopoulos, G. D.; Kyratsous, C. A. Antibody cocktail to SARS-CoV-2 spike protein prevents rapid mutational escape seen with individual antibodies. Science 2020, 369, 1014–1018.

(73) Zost, S. J.; Gilchuk, P.; Case, J. B.; Binshtein, E.; Chen, R. E.; Nkolola, J. P.; Schäfer, A.; Reidy, J. X.; Trivette, A.; Nargi, R. S.; Sutton, R. E.; Suryadevara, N.; Martinez, D. R.; Williamson, L. E.; Chen, E. C.; Jones, T.; Day, S.; Myers, L.; Hassan, A. O.; Kafai, N. M.; Winkler, E. S.; Fox, J. M.; Shrihari, S.; Mueller, B. K.; Meiler, J.; Chandrashekar, A.; Mercado, N. B.; Steinhardt, J. J.; Ren, K.; Loo, Y. M.; Kallewaard, N. L.; McCune, B. T.; Keeler, S. P.; Holtzman, M. J.; Barouch, D. H.; Gralinski, L. E.; Baric, R. S.; Thackray, L. B.; Diamond, M. S.; Carnahan, R. H.; Crowe, J. E., Jr. Potently neutralizing and protective human antibodies against SARS-CoV-2. Nature 2020, 584, 443–449.

(74) Du, S.; Cao, Y.; Zhu, Q.; Yu, P.; Qi, F.; Wang, G.; Du, X.; Bao, L.; Deng, W.; Zhu, H.; Liu, J.; Nie, J.; Zheng, Y.; Liang, H.; Liu, R.; Gong, S.; Xu, H.; Yisimayi, A.; Lv, Q.; Wang, B.; He, R.; Han, Y.; Zhao, W.; Bai, Y.; Qu, Y.; Gao, X.; Ji, C.; Wang, Q.; Gao, N.; Huang, W.; Wang, Y.; Xie, X. S.; Su, X. D.; Xiao, J.; Qin, C. Structurally Resolved SARS-CoV-2 Antibody Shows High Efficacy in Severely Infected Hamsters and Provides a Potent Cocktail Pairing Strategy. Cell 2020, 183, 1013–1023.e13.

(75) Ku, Z.; Xie, X.; Davidson, E.; Ye, X.; Su, H.; Menachery, V. D.; Li, Y.; Yuan, Z.; Zhang, X.; Muruato, A. E.; Ag, I. E.; Tyrell, B.; Doolan, K.; Doranz, B. J.; Wrapp, D.; Bates, P. F.; McLellan, J. S.; Weiss, S. R.; Zhang, N.; Shi, P. Y.; An, Z. Molecular determinants and mechanism for antibody cocktail preventing SARS-CoV-2 escape. Nat Commun 2021, 12, 469.

(76) Wibmer, C. K.; Ayres, F.; Hermanus, T.; Madzivhandila, M.; Kgagudi, P.; Lambson, B. E.; Vermeulen, M.; van den Berg, K.; Rossouw, T.; Boswell, M.; Ueckermann, V.; Meiring, S.; von Gottberg, A.; Cohen, C.; Morris, L.; Bhiman, J. N.; Moore, P. L. SARS-CoV-2 501Y.V2 escapes neutralization by South African COVID-19 donor plasma. bioRxiv 2021. doi: 10.1101/2021.01.18.427166.

(77) Wang, Z.; Schmidt, F.; Weisblum, Y.; Muecksch, F.; Barnes, C. O.; Finkin, S.; Schaefer-Babajew, D.; Cipolla, M.; Gaebler, C.; Lieberman, J. A.; Oliveira, T. Y.; Yang, Z.; Abernathy, M. E.; Huey-Tubman, K. E.; Hurley, A.; Turroja, M.; West, K. A.; Gordon, K.; Millard, K. G.; Ramos, V.; Silva, J. D.; Xu, J.; Colbert, R. A.; Patel, R.; Dizon, J.; Unson-O’Brien, C.; Shimeliovich, I.; Gazumyan, A.; Caskey, M.; Bjorkman, P. J.; Casellas, R.; Hatziioannou, T.; Bieniasz, P. D.; Nussenzweig, M. C. mRNA vaccine-elicited antibodies to SARS-CoV-2 and circulating variants. Nature 2021 doi: 10.1038/s41586-021-03324-6.

(78) Xie, X.; Liu, Y.; Liu, J.; Zhang, X.; Zou, J.; Fontes-Garfias, C. R.; Xia, H.; Swanson, K. A.; Cutler, M.; Cooper, D.; Menachery, V. D.; Weaver, S. C.; Dormitzer, P. R.; Shi, P. Y., Neutralization of SARS-CoV-2 spike 69/70 deletion, E484K and N501Y variants by BNT162b2 vaccine-elicited sera. Nat Med 2021doi: 10.1038/s41591-021-01270-4.

(79) Muik, A.; Wallisch, A. K.; Sänger, B.; Swanson, K. A.; Mühl, J.; Chen, W.; Cai, H.; Maurus, D.; Sarkar, R.; Türeci, Ö.; Dormitzer, P. R.; Şahin, U., Neutralization of SARS-CoV-2 lineage B.1.1.7 pseudovirus by BNT162b2 vaccine-elicited human sera. Science 2021 doi: 10.1126/science.abg6105.

(80) Wu, K.; Werner, A. P.; Moliva, J. I.; Koch, M.; Choi, A.; Stewart-Jones, G. B. E.; Bennett, H.; Boyoglu-Barnum, S.; Shi, W.; Graham, B. S.; Carfi, A.; Corbett, K. S.; Seder, R. A.; Edwards, D. K. mRNA-1273 vaccine induces neutralizing antibodies against spike mutants from global SARS-CoV-2 variants. bioRxiv 2021 doi: 10.1101/2021.01.25.427948.

(81) Thomson, E. C.; Rosen, L. E.; Shepherd, J. G.; Spreafico, R.; da Silva Filipe, A.; Wojcechowskyj, J. A.; Davis, C., et al. Circulating SARS-CoV-2 spike N439K variants maintain fitness while evading antibody-mediated immunity. Cell 2021 https://doi.org/10.1016/j.cell.2021.01.037.

(82) Woo, H.; Park, S. J.; Choi, Y. K.; Park, T.; Tanveer, M.; Cao, Y.; Kern, N. R.; Lee, J.; Yeom, M. S.; Croll, T. I.; Seok, C.; Im, W. Developing a fully glycosylated full-length SARS-CoV-2 spike protein model in a viral membrane. J. Phys. Chem. B 2020, 124, 7128–7137.

(83) Casalino, L.; Gaieb, Z.; Goldsmith, J.A.; Hjorth, C.K.; Dommer, A. C.; Harbison, A. M.; Fogarty, C. A.; Barros, E. P.; Taylor, B. C.; McLellan, J.S.; Fadda, E.; Amaro, R. E. Beyond shielding: The roles of glycans in the SARS-CoV-2 spike potein. ACS Cent. Sci. 2020, 6, 1722–1734.

(84) Kalathiya, U.; Padariya, M.; Mayordomo, M.; Lisowska, M.; Nicholson, J.; Singh, A.; Baginski, M.; Fahraeus, R.; Carragher, N.; Ball, K.; Haas, J.; Daniels, A.; Hupp, T. R.; Alfaro, J. A. Highly conserved homotrimer cavity formed by the SARS-CoV-2 spike glycoprotein: A novel binding site. J. Clin. Med. 2020, 9, 1473.

(85) Brielle, E. S.; Schneidman-Duhovny, D.; Linial, M. The SARS-CoV-2 exerts a distinctive strategy for interacting with the ACE2 human receptor. Viruses 2020, 12,497.

(86) Spinello, A.; Saltalamacchia, A.; Magistrato, A. Is the Rigidity of SARS-CoV-2 spike receptor-binding motif the hallmark for Its enhanced infectivity? Insights from all-atom simulations. J. Phys. Chem. Lett. 2020, 11, 4785–4790.

(87) Chen, Y.; Guo, Y.; Pan, Y.; Zhao, Z. J. Structure analysis of the receptor binding of 2019-nCoV. Biochem. Biophys. Res. Commun. 2020, 525, 135–40.

(88) Othman, H.; Bouslama, Z.; Brandenburg, J. T.; da Rocha, J.; Hamdi, Y.; Ghedira, K.; Srairi-Abid, N.; Hazelhurst, S. Interaction of the spike protein RBD from SARS-CoV-2 with ACE2: Similarity with SARS-CoV, hot-spot analysis and effect of the receptor polymorphism. Biochem. Biophys. Res. Commun. 2020, 527, 702–708.

(89) Ghorbani, M.; Brooks, B. R.; Klauda, J. B. Critical Sequence Hot-spots for Binding of nCOV-2019 to ACE2 as Evaluated by Molecular Simulations. bioRxiv 2020, doi: 10.1101/2020.06.27.175448.

(90) Veeramachaneni, G. K.; Thunuguntla, V.; Bobbillapati, J.; Bondili, J. S. Structural and simulation analysis of hotspot residues interactions of SARS-CoV 2 with human ACE2 receptor. J. Biomol. Struct. Dyn. 2020, 1–11.

(91) Wang, Y.; Liu, M.; Gao, J. Enhanced receptor binding of SARS-CoV-2 through networks of hydrogen-bonding and hydrophobic interactions. Proc. Natl. Acad. Sci. U. S. A. 2020, 117, 13967–13974.

(92) Laurini, E.; Marson, D.; Aulic, S.; Fermeglia, M.; Pricl, S. Computational alanine scanning and structural analysis of the SARS-CoV-2 Spike protein/angiotensin-converting enzyme 2 complex. ACS Nano 2020, 14, 11821–11830.

(93) Ali, A.; Vijayan, R. Dynamics of the ACE2-SARS-CoV-2/SARS-CoV spike protein interface reveal unique mechanisms. Sci. Rep. 2020, 10, 14214.

(94) Verkhivker, G.M. Coevolution, dynamics and allostery conspire in shaping cooperative binding and signal transmission of the SARS-CoV-2 spike protein with human angiotensin-converting enzyme 2. Int. J. Mol. Sci. 2020, 21, 8268.

(95) Di Paola, L.; Hadi-Alijanvand, H.; Song, X.; Hu, G.; Giuliani, A. The discovery of a putative allosteric site in the SARS-CoV-2 spike protein using an integrated structural/dynamic approach. J. Proteome Res. 2020, 19, 4576–4586.

(96) Verkhivker, G.M. Molecular simulations and network modeling reveal an allosteric signaling in the SARS-CoV-2 spike proteins. J. Proteome Res. 2020, 19, 4587–4608.

(97) Teruel, N.; Mailhot, O.; Najmanovich, R.J. Modeling conformational state dynamics and its role on infection for SARS-CoV-2 Spike protein variants. bioRxiv 2020 doi: https://doi.org/10.1101/2020.12.16.423118

(98) Ray, D.; Le, L.; Andricioaei, I. Distant Residues Modulate Conformational Opening in SARS-CoV-2 Spike Protein. bioRxiv 2020 doi: https://doi.org/10.1101/2020.12.07.415596

(99) Cheng, M. H.; Krieger, J.M.; Kaynak, B.; Arditi, M.; Bahar, I. Impact of South African 501.V2 variant on SARS-CoV-2 spike infectivity and neutralization: A Structure-based computational assessment bioRxiv 2021 doi: https://doi.org/10.1101/2021.01.10.426143.

(100) Kolinski, A. Protein modeling and structure prediction with a reduced representation. Acta Biochim. Pol. 2004, 51, 349–371.

(101) Kmiecik, S.; Gront, D.; Kolinski, M.; Wieteska, L.; Dawid, A.E.; Kolinski, A. Coarse-grained protein models and their applications. Chem. Rev. 2016, 116, 7898–7936.

(102) Kmiecik, S.; Kouza, M.; Badaczewska-Dawid, A.E.; Kloczkowski, A.; Kolinski, A. Modeling of protein structural flexibility and large-scale dynamics: Coarse-grained simulations and elastic network models. Int. J. Mol. Sci. 2018, 19, e3496.

(103) Ciemny, M.P.; Badaczewska-Dawid, A.E.; Pikuzinska, M.; Kolinski, A.; Kmiecik, S. Modeling of disordered protein structures using monte carlo simulations and knowledge-based statistical force fields. Int. J. Mol. Sci. 2019, 20, e606.

(104) Kurcinski, M.; Oleniecki, T.; Ciemny, M.P.; Kuriata, A.; Kolinski, A.; Kmiecik, S. CABS-flex standalone: A simulation environment for fast modeling of protein flexibility. Bioinformatics 2019, 35, 694–695.

(105) Hansen, J.; Baum, A.; Pascal, K. E.; Russo, V.; Giordano, S.; Wloga, E.; Fulton, B. O.; Yan, Y.; Koon, K.; Patel, K.; Chung, K. M.; Hermann, A.; Ullman, E.; Cruz, J.; Rafique, A.; Huang, T.; Fairhurst, J.; Libertiny, C.; Malbec, M.; Lee, W. Y.; Welsh, R.; Farr, G.; Pennington, S.; Deshpande, D.; Cheng, J.; Watty, A.; Bouffard, P.; Babb, R.; Levenkova, N.; Chen, C.; Zhang, B.; Romero Hernandez, A.; Saotome, K.; Zhou, Y.; Franklin, M.; Sivapalasingam, S.; Lye, D. C.; Weston, S.; Logue, J.; Haupt, R.; Frieman, M.; Chen, G.; Olson, W.; Murphy, A. J.; Stahl, N.; Yancopoulos, G. D.; Kyratsous, C. A., Studies in humanized mice and convalescent humans yield a SARS-CoV-2 antibody cocktail. Science 2020, 369, 1010–1014.

(106) Berman, H.M.; Westbrook, J.; Feng, Z.; Gilliland, G.; Bhat, T.N.; Weissig, H.; Shindyalov, I.N.; Bourne, P.E. The Protein Data Bank. Nucleic Acids Res. 2000, 28, 235–242.

(107) Rose, P. W.; Prlic, A.; Altunkaya, A.; Bi, C.; Bradley, A. R.; Christie, C. H.; Costanzo, L. D.; Duarte, J. M.; Dutta, S.; Feng, Z.; Green, R. K.; Goodsell, D. S.; Hudson, B.; Kalro, T.; Lowe, R.; Peisach, E.; Randle, C.; Rose, A. S.; Shao, C.; Tao, Y. P.; Valasatava, Y.; Voigt, M.; Westbrook, J. D.; Woo, J.; Yang, H.; Young, J. Y.; Zardecki, C.; Berman, H. M.; Burley, S. K. The RCSB protein data bank: integrative view of protein, gene and 3D structural information. Nucleic Acids Res. 2017, 45, D271–D281.

(108) Hooft, R. W.; Sander, C.; Vriend, G. Positioning hydrogen atoms by optimizing hydrogen-bond networks in protein structures. Proteins 1996, 26, 363–376.

(109) Hekkelman, M. L.; Te Beek, T. A.; Pettifer, S. R.; Thorne, D.; Attwood, T. K.; Vriend, G. WIWS: A protein structure bioinformatics web service collection. Nucleic Acids Res. 2010, 38, W719–W723.

(110) Fiser, A.; Sali, A. ModLoop: Automated modeling of loops in protein structures. Bioinformatics 2003, 19, 2500–25001.

(111) Fernandez-Fuentes, N.; Zhai, J.; Fiser, A. ArchPRED: A template based loop structure prediction server. Nucleic Acids Res. 2006, 34, W173–W176.

(112) Ko, J.; Lee, D.; Park, H.; Coutsias, E. A.; Lee, J.; Seok, C. The FALC-Loop web server for protein loop modeling. Nucleic Acids Res. 2011, 39, W210–W214.

(113) Krivov, G. G.; Shapovalov, M. V.; Dunbrack, R. L., Jr. Improved prediction of protein side-chain conformations with SCWRL4. Proteins 2009, 77, 778–795.

(114) Watanabe, Y.; Berndsen, Z. T.; Raghwani, J.; Seabright, G. E.; Allen, J. D.; Pybus, O. G.; McLellan, J. S.; Wilson, I. A.; Bowden, T. A.; Ward, A. B.; Crispin, M. Vulnerabilities in coronavirus glycan shields despite extensive glycosylation. Nat. Commun. 2020, 11, 2688.

(115) Watanabe, Y.; Allen, J. D.; Wrapp, D.; McLellan, J. S.; Crispin, M. Site-specific glycan analysis of the SARS-CoV-2 spike. Science 2020, 369, 330–333.

(116) Koukos, P.I.; Glykos, N.M. Grcarma: A fully automated task-oriented interface for the analysis of molecular dynamics trajectories. J. Comput. Chem. 2013, 34, 2310–2312.

(117) Bahar, I.; Lezon, T. R.; Yang, L. W.; Eyal, E. Global dynamics of proteins: bridging between structure and function. Annu. Rev. Biophys. 2010, 39, 23–42.

(118) Yang, L. W.; Rader, A. J.; Liu, X.; Jursa, C. J.; Chen, S. C.; Karimi, H. A.; Bahar, I. oGNM: online computation of structural dynamics using the Gaussian Network Model. Nucleic Acids Res. 2006, 34, W24–W31.

(119) Eyal, E.; Lum, G.; Bahar, I. The anisotropic network model web server at 2015 (ANM 2.0). Bioinformatics 2015, 31, 1487–1489.

(120) Li, H.; Chang, Y. Y.; Lee, J. Y.; Bahar, I.; Yang, L. W. DynOmics: dynamics of structural proteome and beyond. Nucleic Acids Res. 2017, 45, W374–W380.

(122) Dehouck, Y.; Kwasigroch, J. M.; Rooman, M.; Gilis, D. BeAtMuSiC: Prediction of changes in protein-protein binding affinity on mutations. Nucleic Acids Res. 2013, 41, W333–W339.

(123) Dehouck, Y.; Gilis, D.; Rooman, M. A new generation of statistical potentials for proteins. Biophys. J. 2006, 90, 4010–4017.

(124) Atilgan, C.; Atilgan, A. R., Perturbation-response scanning reveals ligand entry-exit mechanisms of ferric binding protein. PLoS Comput. Biol. 2009, 5, e1000544.

(125) Atilgan, C.; Gerek, Z. N.; Ozkan, S. B.; Atilgan, A. R., Manipulation of conformational change in proteins by single-residue perturbations. Biophys. J. 2010, 99, 933–943.

(126) General, I. J.; Liu, Y.; Blackburn, M. E.; Mao, W.; Gierasch, L. M.; Bahar, I., ATPase subdomain IA is a mediator of interdomain allostery in Hsp70 molecular chaperones. PLoS Comput. Biol. 2014, 10, e1003624.

(127) Dutta, A.; Krieger, J.; Lee, J. Y.; Garcia-Nafria, J.; Greger, I. H.; Bahar, I., Cooperative Dynamics of Intact AMPA and NMDA Glutamate Receptors: Similarities and Subfamily-Specific Differences. Structure 2015, 23, 1692–1704.

(128) Penkler, D.; Sensoy, O.; Atilgan, C.; Tastan Bishop, O., Perturbation-Response Scanning Reveals Key Residues for Allosteric Control in Hsp70. J. Chem. Inf. Model. 2017, 57, 1359–1374.

(129) Penkler, D. L.; Atilgan, C.; Tastan Bishop, O., Allosteric Modulation of Human Hsp90alpha Conformational Dynamics. J. Chem. Inf. Model. 2018, 58, 383–404.

(130) Stetz, G.; Tse, A.; Verkhivker, G. M., Dissecting Structure-Encoded Determinants of Allosteric Cross-Talk between Post-Translational Modification Sites in the Hsp90 Chaperones. Sci. Rep. 2018, 8, 19.

(131) Jalalypour, F.; Sensoy, O.; Atilgan, C., Perturb-Scan-Pull: A Novel Method Facilitating Conformational Transitions in Proteins. J. Chem. Theory Comput. 2020, 16, 3825–3841.

(132) Brinda, K.V.; Vishveshwara, S. A network representation of protein structures: Implications for protein stability. Biophys. J. 2005, 89, 4159–4170.

(133) Vijayabaskar, M.S.; Vishveshwara, S. Interaction energy based protein structure networks. Biophys. J. 2010, 99, 3704–3715.

(134) Sethi, A.; Eargle, J.; Black, A.A.; Luthey-Schulten, Z. Dynamical networks in tRNA:protein complexes. Proc. Natl. Acad. Sci. U.S.A. 2009, 106, 6620–6625.

(135) Stetz, G.; Verkhivker, G. M. Computational analysis of residue interaction networks and coevolutionary relationships in the Hsp70 chaperones: A community-hopping model of allosteric regulation and communication. Plos Comput. Biol. 2017, 13, e1005299.

(136) Czemeres, J.; Buse, K.; Verkhivker, G. M. Atomistic simulations and network-based modeling of the Hsp90-Cdc37 chaperone binding with Cdk4 client protein: A mechanism of chaperoning kinase clients by exploiting weak spots of intrinsically dynamic kinase domains. PLoS One 2017, 12, e0190267.

(137) Stetz, G.; Verkhivker, G. M. Dancing through life: Molecular dynamics simulations and network-centric modeling of allosteric mechanisms in Hsp70 and Hsp110 chaperone proteins. PLoS One 2015, 10, e0143752.61.

(138) Floyd, R.W. Algorithm 97: Shortest path. Commun. A.C.M. 1962, 5, 345.

(139) Hagberg, A.A.; Schult, D.A.; Swart, P.J. Exploring network structure, dynamics, and function using NetworkX, in : G. Varoquaux, T. Vaught, J. Millman (Eds.), Proceedings of the 7th Python in Science Conference (SciPy2008), Pasadena, 2008, pp. 11–15.

(140) Girvan, M.; Newman, M. E. Community Structure in Social and Biological Networks. Proc. Natl. Acad. Sci. U. S. A. 2002, 99, 7821–7826.

(141) Newman, M. E.; Girvan, M. Finding and Evaluating Community Structure in Networks. Phys. Rev. E Stat. Nonlin. Soft Matter Phys. 2004, 69, 026113.

(142) Newman, M. E. Modularity and Community Structure in Networks. Proc. Natl. Acad. Sci. U. S. A. 2006, 103, 8577–8582.

(143) Astl, L.; Verkhivker, G. M. Atomistic modeling of the ABL kinase regulation by allosteric modulators using structural perturbation analysis and community-based network reconstruction of allosteric communications. J. Chem. Theory Comput. 2019, 15, 3362–3380.

(144) Astl, L.; Verkhivker, G.M. Dynamic view of allosteric regulation in the Hsp70 chaperones by J-Domain cochaperone and post-translational modifications: Computational analysis of Hsp70 mechanisms by exploring conformational landscapes and residue interaction networks. J. Chem. Inf. Model. 2020, 60, 1614–1631.

(145) Shannon, P.; Markiel, A.; Ozier, O.; Baliga, N. S.; Wang, J. T.; Ramage, D.; Amin, N.; Schwikowski, B.; Ideker, T. Cytoscape: A software environment for integrated models of biomolecular interaction networks. Genome Res 2003, 13, 2498–2504.

(146) Kovács, I.A.; Palotai, R.; Szalay, M.S.; Csermely, P. Community landscapes: an integrative approach to determine overlapping network module hierarchy, identify key nodes and predict network dynamics. PLoS One 2010, 5, e12528.

(147) Ni, D.; Lau, K.; Lehmann, F.; Fränkl, A.; Hacker, D.; Pojer, F.; Stahlberg, H. Structural investigation of ACE2 dependent disassembly of the trimeric SARS-CoV-2 Spike glycoprotein. bioRxiv 2020 doi: https://doi.org/10.1101/2020.10.12.336016.

(148) Verkhivker, G. M.; Di Paola, L., Dynamic Network Modeling of Allosteric Interactions and Communication Pathways in the SARS-CoV-2 Spike Trimer Mutants: Differential Modulation of Conformational Landscapes and Signal Transmission via Cascades of Regulatory Switches. J Phys Chem B 2021, 125, 850–873.

(149) Navizet, I.; Cailliez, F.; Lavery, R., Probing protein mechanics: residue-level properties and their use in defining domains. Biophys. J. 2004, 87, 1426–1435.

(150) Sacquin-Mora, S.; Lavery, R., Investigating the local flexibility of functional residues in hemoproteins. Biophys. J. 2006, 90, 2706–2717.

(151) Sacquin-Mora, S.; Laforet, E.; Lavery, R., Locating the active sites of enzymes using mechanical properties. Proteins 2007, 67, 350–359.

(152) Lavery, R.; Sacquin-Mora, S., Protein mechanics: a route from structure to function. J. Biosci. 2007, 32, 891–898.

(153) Blacklock, K.; Verkhivker, G. M., Computational modeling of allosteric regulation in the hsp90 chaperones: a statistical ensemble analysis of protein structure networks and allosteric

(154) Rader, A. J.; Brown, S. M., Correlating allostery with rigidity. Mol Biosyst 2011, 7, 464–471.

(155) Rader, A.J.; Yennamalli, R.M.; Harter, A.K.; Sen, T.Z. A rigid network of long-range contacts increases thermostability in a mutant endoglucanase. J. Biomol. Struct. Dyn. 2012, 30, 628–637.

(156) Leander, M.; Yuan, Y.; Meger, A.; Cui, Q.; Raman, S., Functional plasticity and evolutionary adaptation of allosteric regulation. Proc Natl Acad Sci U S A 2020, 117 (41), 25445–25454.

