## Supplemental Information for "Comparative Perturbation-Based Modeling of the SARS-CoV-2 Spike Protein Binding with Host Receptor and Neutralizing Antibodies : Structurally Adaptable Allosteric Communication Hotspots Define Spike Sites Targeted by Global Circulating Mutations"

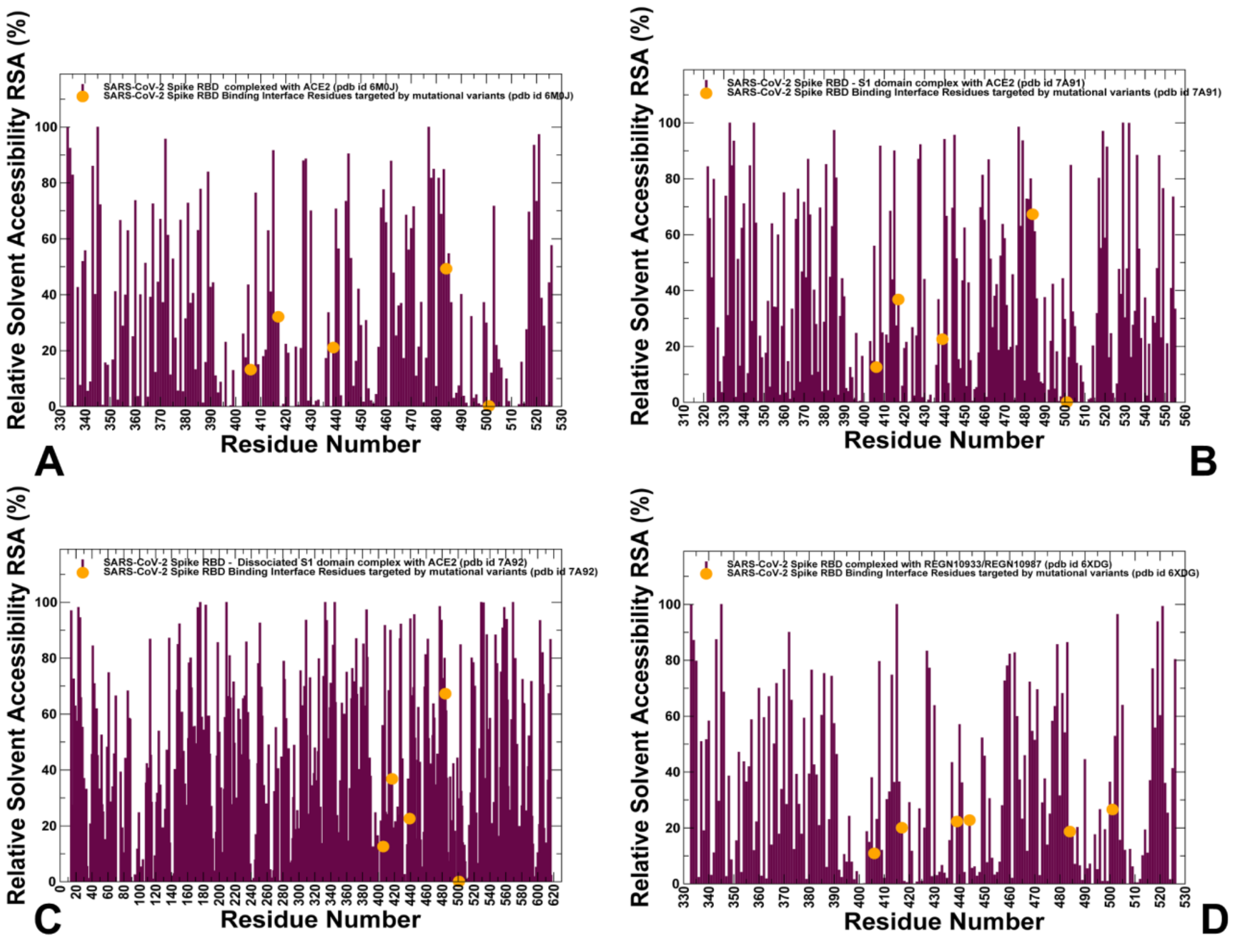


**Figure S1.** The average relative solvent accessibility (RSA) percentage of the protein residues for the SARS-CoV-2 S-RBD and S1 domain complexes with ACE2 and REGN-COV2 antibody cocktail. (A) The RSA profile of the bound SARS-CoV-2 S-RBD structure in the complex with ACE2 ( pdb id 6M0J). (B) The RSA profile of the bound SARS-CoV-2 S1-RBD structure in the complex with ACE2 ( pdb id 7A91). (C) The RSA profile of the bound S1 domain structure in the complex with ACE2 ( pdb id 7A92). (D) The RSA profile of the bound SARS-CoV-2 S-RBD structure in the complex with the REGN-COV2 antibody combination ( pdb id 6XDG). The RSA profiles are shown in maroon lines and positions of key functional residues E406, K417, N439, E484, and N501 are highlighted by filled green circles.

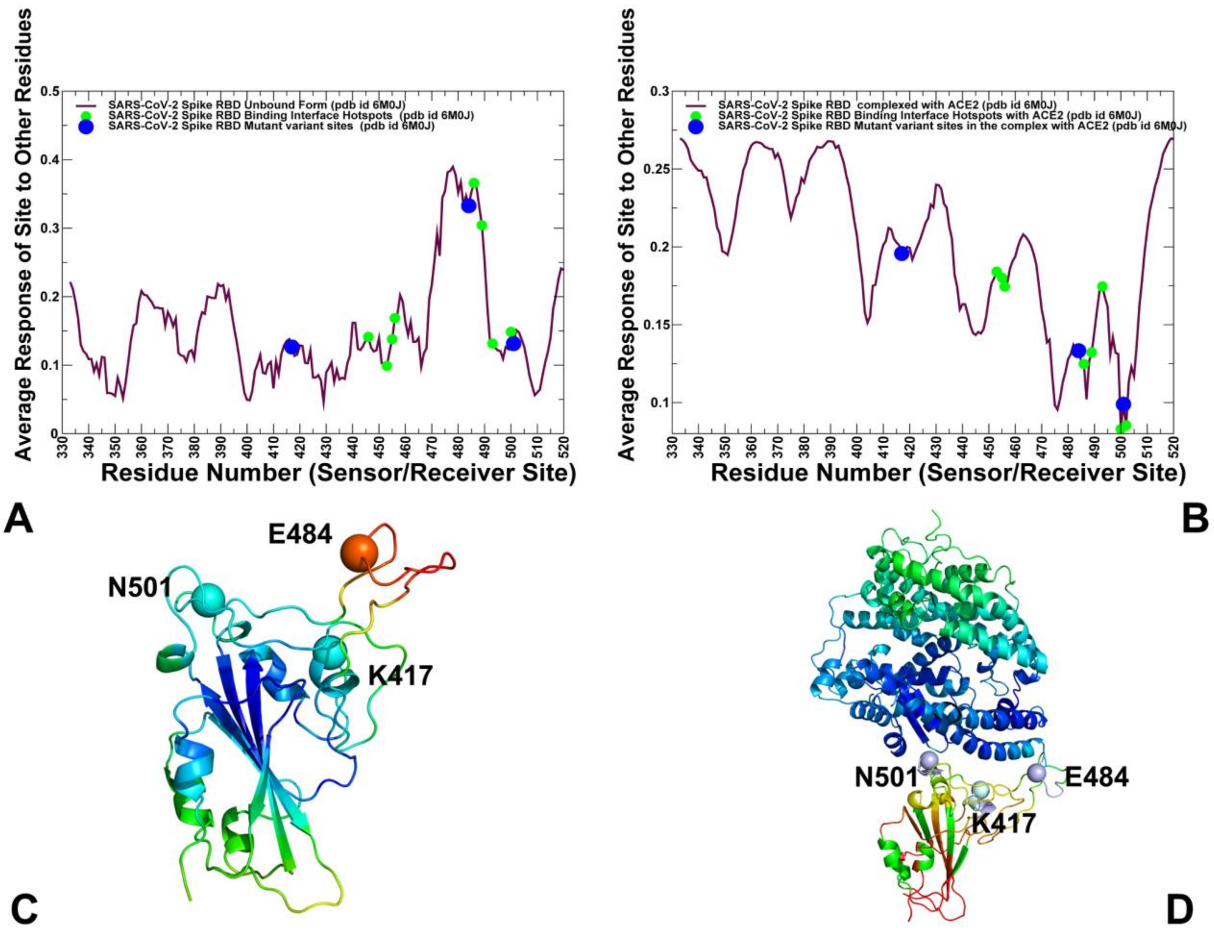


**Figure S2.** The PRS sensor profiles for the unbound and bound forms of the SARS-CoV-2 S-RBD with ACE2. (A) The PRS sensor profile for the unbound SARS-CoV-2 S-RBD (pdb id 6M0J). (B) The PRS sensor profile for the SARSCoV-2 S-RBD complex with ACE2 (pdb id 6M0J). (C) Structural map of the sensor profile for the unbound SARS-CoV-2 S-RBD. (D) Structural map of the sensor profile for the SARSCoV-2 S-RBD complex with ACE2. Structural maps of the PRS sensor profiles are shown with the color gradient from blue to red indicating the increasing sensor propensities. The positions of functional residues K417, E484, and N501 are indicated by spheres colored according to the sensor propensity level.

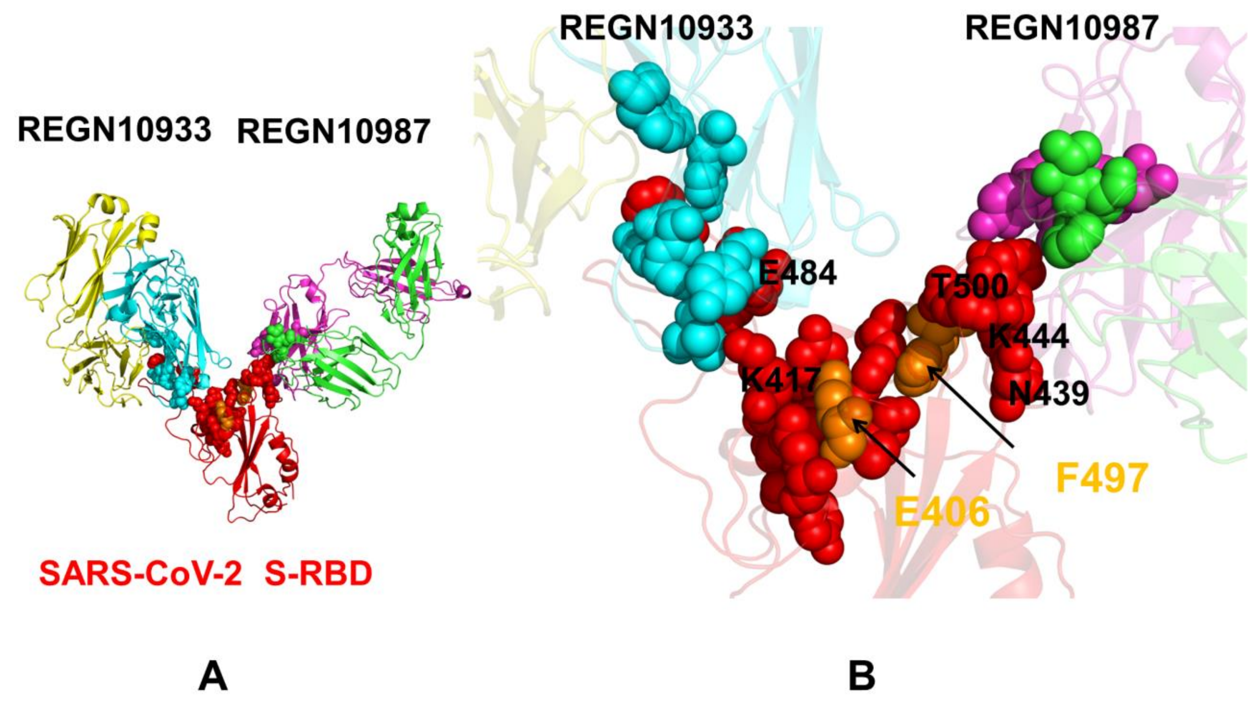


**Figure S3.** Structural maps of local communities in the SARS-CoV-2 S-RBD complex with REGN-COV2 cocktail of antibodies. (A) A general community overview of the SARS-CoV-2 S-RBD complex with REGN-COV2. The RBD is in red ribbons and interfacial communities are shown in red spheres. (B) A close-up of the intermolecular communities formed by the S-RBD. The key inter-community bridging sites E406 and F497 that effectively link local intermolecular modules interacting with REGN10933 and REGN10987 antibodies are shown in orange spheres and annotated. The positions of positions targeted by circulating and antibody-escaping mutations K417, E484, N439, and K444 are annotated.

**Table S1.** The list of the intermolecular contacts in the structure of SARS-CoV-2 RBD complex with ACE2 (pdb id 6M0J).

| **Residue Name** | **Residue Number** | **SARS-RBD** | **Residue Name** | **Residue Number** | **ACE2** |
|---|---|---|---|---|---|
| LYS | 417 | E | ASP | 30 | A |
| LYS | 417 | E | HIS | 34 | A |
| GLY | 446 | E | LEU | 45 | A |
| GLY | 446 | E | GLN | 42 | A |
| GLY | 447 | E | GLN | 42 | A |
| TYR | 449 | E | GLN | 42 | A |
| TYR | 449 | E | ASP | 38 | A |
| TYR | 453 | E | HIS | 34 | A |
| LEU | 455 | E | LYS | 31 | A |
| LEU | 455 | E | HIS | 34 | A |
| LEU | 455 | E | ASP | 30 | A |
| PHE | 456 | E | THR | 27 | A |
| PHE | 456 | E | ASP | 30 | A |
| PHE | 456 | E | LYS | 31 | A |
| TYR | 473 | E | THR | 27 | A |
| ALA | 475 | E | SER | 19 | A |
| ALA | 475 | E | GLN | 24 | A |
| ALA | 475 | E | THR | 27 | A |
| GLY | 476 | E | GLN | 24 | A |
| SER | 477 | E | GLN | 24 | A |
| GLU | 484 | E | LYS | 31 | A |
| PHE | 486 | E | GLN | 24 | A |
| PHE | 486 | E | TYR | 83 | A |
| PHE | 486 | E | MET | 82 | A |
| PHE | 486 | E | LEU | 79 | A |
| ASN | 487 | E | GLN | 24 | A |
| ASN | 487 | E | TYR | 83 | A |
| ASN | 487 | E | PHE | 28 | A |
| TYR | 489 | E | PHE | 28 | A |
| TYR | 489 | E | GLN | 24 | A |
| TYR | 489 | E | TYR | 83 | A |
| TYR | 489 | E | THR | 27 | A |
| TYR | 489 | E | LYS | 31 | A |
| PHE | 490 | E | LYS | 31 | A |

| | | | | | |
|---|---|---|---|---|---|
| GLN | 493 | E | LYS | 31 | A |
| GLN | 493 | E | HIS | 34 | A |
| GLN | 493 | E | GLU | 35 | A |
| TYR | 495 | E | LYS | 353 | A |
| GLY | 496 | E | LYS | 353 | A |
| GLY | 496 | E | ASP | 38 | A |
| PHE | 497 | E | LYS | 353 | A |
| GLN | 498 | E | LEU | 45 | A |
| GLN | 498 | E | TYR | 41 | A |
| GLN | 498 | E | LYS | 353 | A |
| GLN | 498 | E | GLN | 42 | A |
| GLN | 498 | E | ASP | 38 | A |
| THR | 500 | E | LEU | 45 | A |
| THR | 500 | E | GLY | 354 | A |
| THR | 500 | E | TYR | 41 | A |
| THR | 500 | E | ASN | 330 | A |
| THR | 500 | E | ARG | 357 | A |
| THR | 500 | E | LYS | 353 | A |
| THR | 500 | E | ASP | 355 | A |
| ASN | 501 | E | LYS | 353 | A |
| ASN | 501 | E | TYR | 41 | A |
| ASN | 501 | E | GLY | 354 | A |
| ASN | 501 | E | ASP | 355 | A |
| GLY | 502 | E | LYS | 353 | A |
| GLY | 502 | E | GLY | 354 | A |
| GLY | 502 | E | ASP | 355 | A |
| VAL | 503 | E | GLY | 354 | A |
| TYR | 505 | E | GLY | 354 | A |
| TYR | 505 | E | ARG | 393 | A |
| TYR | 505 | E | ALA | 386 | A |
| TYR | 505 | E | LYS | 353 | A |
| TYR | 505 | E | GLU | 37 | A |

**Table S2.** The list of the intermolecular contacts in the structure of SARS-CoV-2 S1 domain complex with ACE2 (pdb id 7A91).

| Residue Name | Residue Number | SARS-RBD | Residue Name | Residue Number | ACE2 |
|---|---|---|---|---|---|
| LYS | 417 | A | ASP | 30 | D |
| GLY | 446 | A | GLN | 42 | D |
| TYR | 449 | A | LYS | 353 | D |
| TYR | 449 | A | GLN | 42 | D |
| TYR | 449 | A | ASP | 38 | D |
| TYR | 453 | A | HIS | 34 | D |
| LEU | 455 | A | LYS | 31 | D |
| LEU | 455 | A | ASP | 30 | D |
| LEU | 455 | A | HIS | 34 | D |
| PHE | 456 | A | ASP | 30 | D |
| PHE | 456 | A | LYS | 31 | D |
| PHE | 456 | A | THR | 27 | D |
| TYR | 473 | A | THR | 27 | D |
| ALA | 475 | A | GLU | 23 | D |
| ALA | 475 | A | GLN | 24 | D |
| ALA | 475 | A | THR | 27 | D |
| ALA | 475 | A | SER | 19 | D |
| GLY | 476 | A | SER | 19 | D |
| GLY | 476 | A | GLN | 24 | D |
| SER | 477 | A | SER | 19 | D |
| SER | 477 | A | GLN | 24 | D |
| GLY | 485 | A | LEU | 79 | D |
| PHE | 486 | A | TYR | 83 | D |
| PHE | 486 | A | LEU | 79 | D |
| PHE | 486 | A | MET | 82 | D |
| ASN | 487 | A | GLN | 24 | D |
| ASN | 487 | A | TYR | 83 | D |
| TYR | 489 | A | THR | 27 | D |
| TYR | 489 | A | PHE | 28 | D |
| TYR | 489 | A | TYR | 83 | D |
| TYR | 489 | A | LEU | 79 | D |
| TYR | 489 | A | LYS | 31 | D |
| PHE | 490 | A | LYS | 31 | D |
| GLN | 493 | A | GLU | 35 | D |
| GLN | 493 | A | HIS | 34 | D |
| GLN | 493 | A | LYS | 31 | D |

| | | | | | |
|---|---|---|---|---|---|
| SER | 494 | A | HIS | 34 | D |
| TYR | 495 | A | HIS | 34 | D |
| GLY | 496 | A | LYS | 353 | D |
| PHE | 497 | A | LYS | 353 | D |
| GLN | 498 | A | LYS | 353 | D |
| GLN | 498 | A | TYR | 41 | D |
| GLN | 498 | A | ASP | 38 | D |
| GLN | 498 | A | GLN | 42 | D |
| GLN | 498 | A | LEU | 45 | D |
| THR | 500 | A | TYR | 41 | D |
| THR | 500 | A | ASN | 330 | D |
| THR | 500 | A | ASP | 355 | D |
| THR | 500 | A | LYS | 353 | D |
| THR | 500 | A | LEU | 45 | D |
| THR | 500 | A | GLY | 354 | D |
| THR | 500 | A | ARG | 357 | D |
| ASN | 501 | A | ASP | 355 | D |
| ASN | 501 | A | TYR | 41 | D |
| ASN | 501 | A | LYS | 353 | D |
| ASN | 501 | A | GLY | 354 | D |
| GLY | 502 | A | LYS | 353 | D |
| GLY | 502 | A | GLY | 354 | D |
| GLY | 502 | A | ASP | 355 | D |
| VAL | 503 | A | GLY | 354 | D |
| TYR | 505 | A | GLU | 37 | D |
| TYR | 505 | A | ALA | 386 | D |
| TYR | 505 | A | ARG | 393 | D |
| TYR | 505 | A | LYS | 353 | D |
| TYR | 505 | A | GLY | 354 | D |

**Table S3.** The list of the intermolecular contacts in the structure of SARS-CoV-2 S1 domain complex with ACE2 (pdb id 7A92).

| **Residue Name** | **Residue Number** | **SARS-RBD** | **Residue Name** | **Residue Number** | **ACE2** |
|---|---|---|---|---|---|
| LYS | 417 | A | ASP | 30 | D |
| GLY | 446 | A | GLN | 42 | D |
| TYR | 449 | A | LYS | 353 | D |
| TYR | 449 | A | ASP | 38 | D |
| TYR | 449 | A | GLN | 42 | D |
| TYR | 453 | A | HIS | 34 | D |
| LEU | 455 | A | HIS | 34 | D |
| LEU | 455 | A | LYS | 31 | D |
| LEU | 455 | A | ASP | 30 | D |
| PHE | 456 | A | ASP | 30 | D |
| PHE | 456 | A | LYS | 31 | D |
| PHE | 456 | A | THR | 27 | D |
| TYR | 473 | A | THR | 27 | D |
| ALA | 475 | A | THR | 27 | D |
| ALA | 475 | A | GLN | 24 | D |
| ALA | 475 | A | SER | 19 | D |
| ALA | 475 | A | GLU | 23 | D |
| GLY | 476 | A | SER | 19 | D |
| GLY | 476 | A | GLN | 24 | D |
| SER | 477 | A | GLN | 24 | D |
| SER | 477 | A | SER | 19 | D |
| GLY | 485 | A | LEU | 79 | D |
| PHE | 486 | A | LEU | 79 | D |
| PHE | 486 | A | TYR | 83 | D |
| PHE | 486 | A | MET | 82 | D |
| ASN | 487 | A | TYR | 83 | D |
| ASN | 487 | A | GLN | 24 | D |
| TYR | 489 | A | LYS | 31 | D |
| TYR | 489 | A | PHE | 28 | D |
| TYR | 489 | A | TYR | 83 | D |
| TYR | 489 | A | LEU | 79 | D |
| TYR | 489 | A | THR | 27 | D |
| PHE | 490 | A | LYS | 31 | D |
| GLN | 493 | A | HIS | 34 | D |
| GLN | 493 | A | GLU | 35 | D |
| GLN | 493 | A | LYS | 31 | D |

| | | | | | |
|---|---|---|---|---|---|
| SER | 494 | A | HIS | 34 | D |
| TYR | 495 | A | HIS | 34 | D |
| GLY | 496 | A | LYS | 353 | D |
| PHE | 497 | A | LYS | 353 | D |
| GLN | 498 | A | LYS | 353 | D |
| GLN | 498 | A | ASP | 38 | D |
| GLN | 498 | A | LEU | 45 | D |
| GLN | 498 | A | GLN | 42 | D |
| GLN | 498 | A | TYR | 41 | D |
| THR | 500 | A | TYR | 41 | D |
| THR | 500 | A | LYS | 353 | D |
| THR | 500 | A | GLY | 354 | D |
| THR | 500 | A | LEU | 45 | D |
| THR | 500 | A | ARG | 357 | D |
| THR | 500 | A | ASN | 330 | D |
| THR | 500 | A | ASP | 355 | D |
| ASN | 501 | A | TYR | 41 | D |
| ASN | 501 | A | LYS | 353 | D |
| ASN | 501 | A | GLY | 354 | D |
| ASN | 501 | A | ASP | 355 | D |
| GLY | 502 | A | GLY | 354 | D |
| GLY | 502 | A | LYS | 353 | D |
| GLY | 502 | A | ASP | 355 | D |
| VAL | 503 | A | GLY | 354 | D |
| TYR | 505 | A | ARG | 393 | D |
| TYR | 505 | A | GLU | 37 | D |
| TYR | 505 | A | GLY | 354 | D |
| TYR | 505 | A | ALA | 386 | D |
| TYR | 505 | A | LYS | 353 | D |

**Table S4.** The list of the intermolecular contacts in the structure of SARS-CoV-2 RBD domain complex with REGN-COV2 antibody cocktail (pdb id 6XDG).

Chain E: SPIKE PROTEIN S1
Chain D: REGN10933 ANTIBODY FAB FRAGMENT LIGHT CHAIN
Chain B: REGN10933 ANTIBODY FAB FRAGMENT HEAVY CHAIN
Chain C: REGN10987 ANTIBODY FAB FRAGMENT HEAVY CHAIN
Chain A: REGN10987 ANTIBODY FAB FRAGMENT LIGHT CHAIN

| **Residue Name** | **Residue Number** | **SARS-RBD** | **Residue Name** | **Residue Name** | **REGN-COV2** |
|---|---|---|---|---|---|
| ARG | 346 | E | ASN | 31 | C |
| ARG | 403 | E | THR | 28 | B |
| GLU | 406 | E | THR | 28 | B |
| LYS | 417 | E | ASP | 31 | B |
| LYS | 417 | E | THR | 102 | B |
| LYS | 417 | E | TYR | 32 | B |
| LYS | 417 | E | THR | 28 | B |
| TYR | 421 | E | THR | 103 | B |
| TYR | 421 | E | THR | 102 | B |
| ASN | 439 | E | GLY | 103 | C |
| ASN | 439 | E | ASP | 104 | C |
| ASN | 440 | E | ASP | 101 | C |
| ASN | 440 | E | GLY | 103 | C |
| ASN | 440 | E | TYR | 102 | C |
| ASN | 440 | E | ASP | 104 | C |
| LEU | 441 | E | GLY | 103 | C |
| LEU | 441 | E | TYR | 102 | C |
| LEU | 441 | E | ASP | 101 | C |
| SER | 443 | E | GLY | 103 | C |
| SER | 443 | E | ASP | 104 | C |
| LYS | 444 | E | TYR | 32 | C |
| LYS | 444 | E | ASN | 31 | C |
| LYS | 444 | E | TYR | 53 | C |
| VAL | 445 | E | ILE | 51 | C |
| VAL | 445 | E | ALA | 33 | C |
| VAL | 445 | E | SER | 52 | C |
| VAL | 445 | E | TRP | 99 | A |
| VAL | 445 | E | ASN | 57 | C |
| VAL | 445 | E | TYR | 53 | C |
| VAL | 445 | E | TYR | 59 | C |

| | | | | | |
|---|---|---|---|---|---|
| VAL | 445 | E | TYR | 35 | C |
| VAL | 445 | E | TYR | 105 | C |
| VAL | 445 | E | VAL | 50 | C |
| GLY | 446 | E | SER | 52 | C |
| GLY | 446 | E | TYR | 59 | C |
| GLY | 446 | E | ASN | 57 | C |
| GLY | 447 | E | TYR | 59 | C |
| GLY | 447 | E | ASP | 54 | C |
| GLY | 447 | E | ASN | 57 | C |
| GLY | 447 | E | TYR | 53 | C |
| GLY | 447 | E | SER | 52 | C |
| ASN | 448 | E | TYR | 53 | C |
| TYR | 449 | E | TYR | 53 | C |
| TYR | 449 | E | ASN | 57 | C |
| TYR | 449 | E | SER | 56 | C |
| TYR | 449 | E | ASP | 54 | C |
| ASN | 450 | E | TYR | 53 | C |
| TYR | 453 | E | SER | 30 | B |
| TYR | 453 | E | ASP | 31 | B |
| LEU | 455 | E | THR | 102 | B |
| LEU | 455 | E | SER | 30 | B |
| LEU | 455 | E | ASP | 31 | B |
| LEU | 455 | E | TYR | 32 | B |
| PHE | 456 | E | GLY | 101 | B |
| PHE | 456 | E | MET | 104 | B |
| PHE | 456 | E | ARG | 100 | B |
| PHE | 456 | E | THR | 103 | B |
| PHE | 456 | E | ASP | 31 | B |
| PHE | 456 | E | THR | 102 | B |
| TYR | 473 | E | MET | 104 | B |
| ALA | 475 | E | MET | 104 | B |
| ALA | 475 | E | TYR | 32 | D |
| GLY | 476 | E | TYR | 32 | D |
| GLY | 476 | E | ASP | 92 | D |
| SER | 477 | E | ASP | 92 | D |
| SER | 477 | E | TYR | 32 | D |
| THR | 478 | E | ASN | 93 | D |
| GLU | 484 | E | THR | 52 | B |
| GLU | 484 | E | TYR | 53 | B |
| GLU | 484 | E | SER | 56 | B |
| GLU | 484 | E | TYR | 59 | B |

| | | | | | |
|---|---|---|---|---|---|
| GLU | 484 | E | SER | 54 | B |
| GLU | 484 | E | THR | 57 | B |
| GLY | 485 | E | TYR | 59 | B |
| GLY | 485 | E | THR | 57 | B |
| GLY | 485 | E | TYR | 33 | B |
| GLY | 485 | E | THR | 52 | B |
| PHE | 486 | E | LEU | 96 | D |
| PHE | 486 | E | TYR | 33 | B |
| PHE | 486 | E | ARG | 100 | B |
| PHE | 486 | E | TYR | 59 | B |
| PHE | 486 | E | ASP | 92 | D |
| PHE | 486 | E | THR | 57 | B |
| PHE | 486 | E | LEU | 94 | D |
| PHE | 486 | E | ASN | 93 | D |
| PHE | 486 | E | TYR | 50 | B |
| PHE | 486 | E | TYR | 91 | D |
| ASN | 487 | E | ASP | 92 | D |
| ASN | 487 | E | ARG | 100 | B |
| ASN | 487 | E | TYR | 59 | B |
| ASN | 487 | E | TYR | 33 | B |
| CYS | 488 | E | ARG | 100 | B |
| CYS | 488 | E | TYR | 33 | B |
| TYR | 489 | E | SER | 30 | B |
| TYR | 489 | E | THR | 52 | B |
| TYR | 489 | E | TYR | 33 | B |
| TYR | 489 | E | TYR | 53 | B |
| TYR | 489 | E | ASP | 31 | B |
| TYR | 489 | E | TYR | 32 | B |
| PHE | 490 | E | TYR | 53 | B |
| LEU | 492 | E | TYR | 53 | B |
| GLN | 493 | E | SER | 30 | B |
| GLN | 493 | E | ASP | 31 | B |
| GLN | 493 | E | SER | 54 | B |
| GLN | 493 | E | TYR | 53 | B |
| GLN | 493 | E | ASN | 74 | B |
| SER | 494 | E | ASN | 74 | B |
| GLN | 498 | E | ALA | 75 | B |
| GLN | 498 | E | TYR | 59 | C |
| PRO | 499 | E | TYR | 105 | C |
| PRO | 499 | E | LEU | 93 | A |
| PRO | 499 | E | TYR | 34 | A |

| | | | | | |
|---|---|---|---|---|---|
| PRO | 499 | E | ASP | 104 | C |
| THR | 500 | E | SER | 95 | A |
| THR | 500 | E | LEU | 93 | A |
| THR | 500 | E | TRP | 99 | A |
| THR | 500 | E | TYR | 32 | A |
| ASN | 501 | E | TYR | 32 | A |
